# Integrated phenotypic screening and chemical proteomics identifies ETF1 ligands that modulate viral translation and replication

**DOI:** 10.1101/2025.08.27.672717

**Authors:** Arthur S. Kim, Kevin Ma, Christopher J. Reinhardt, Daniel C. Lazar, Daisuke Ogasawara, Teressa M. Shaw, Juan Carlos de la Torre, Adam L. Bailey, Bruno Melillo, John R. Teijaro, Benjamin F. Cravatt

## Abstract

Emerging and re-emerging viruses pose a significant threat to global health. Although direct-acting antivirals have shown success, their efficacy is limited by the rapid emergence of drug-resistant viral variants. Hence, there is an urgent need for additional broad spectrum antiviral therapeutic strategies. Here, we identify by phenotypic screening a set of stereochemically defined photoreactive small molecules (photo-stereoprobes) that stereoselectively suppress SARS-CoV-2 replication in human lung epithelial cells. Structure-activity relationship-guided chemical proteomics identified the eukaryotic translation termination factor 1 (ETF1) as a target of the photo-stereoprobes, and this interaction was recapitulated with recombinant purified ETF1. We found that the photo-stereoprobes modulate programmed ribosomal frameshifting mechanisms essential for SARS-CoV-2 infection without causing ETF1 degradation, thus distinguishing the photo-stereoprobes from other known ETF1-directed small molecules. We finally show that the photo-stereoprobes also inhibit the replication of additional viruses with non-canonical ribosomal frameshifting mechanisms. Our findings thus identify a mechanistically distinct class of ETF1 ligands that implicate host translation termination processes as a potential target for antiviral development.

**SIGNIFICANCE STATEMENT:** The identification of broad-spectrum antivirals that target host proteins is a desirable strategy to combat emerging viral infections given the rapid escape potential of viruses and the need to develop new countermeasures for clinically significant pathogens. Here, we integrate phenotypic screening and chemical proteomics to identify photo-stereoprobe small molecules that inhibit the replication of multiple viruses. We show that these compounds bind the protein eukaryotic peptide chain release factor subunit 1 (ETF1) and modulate programmed ribosomal frameshifting. Unlike previously described ligands for ETF1, which lead to proteasomal destruction of the protein, we did not find that the photo-stereoprobes altered ETF1 content in cells. Our findings thus point to an opportunity to pharmacologically modulate a host protein implicated in programmed ribosomal frameshifting as a strategy to combat infection of viruses from different families.

## INTRODUCTION

Emerging infectious diseases pose a significant threat to global health, as evidenced by the SARS-CoV-2 pandemic. Although the deployment of COVID-19 vaccines and therapeutics mitigate the risk of severe COVID-19 disease, the continuous emergence of SARS-CoV-2 variants along with other emerging viruses highlights the need for additional therapeutic countermeasures. Current antiviral strategies against SARS-CoV-2 are predominantly directed against a small subset of viral proteins, particularly those that are essential for viral replication. Examples include the RNA-dependent RNA polymerase inhibitors remdesivir and molnupiravir and the main protease (Mpro) inhibitors nirmatrelvir and ensitrelvir (1–4). However, these direct-acting antivirals have limitations, in particular, the emergence of viral variants that confer drug resistance (5, 6).

An alternative strategy would be to develop antivirals directed against host protein targets, which might allow for: 1) broad-spectrum antiviral efficacy, as viruses often hijack shared host proteins for replication (7, 8); and 2) reduced emergence of drug-resistant viral mutations given the decreased genetic variability of host factors (9). Strategies to identify host factors important for viral infection have included functional genomic screens (10–14), biochemical approaches (15, 16), and comparing host factors in cell lines that are permissive or non-permissive to viral infection (17, 18). Each of these approaches has different strengths and limitations, but they collectively share the shortcoming that additional work is required to identify chemical tools to pharmacologically characterize the nominated antiviral host proteins, which is an important step in the drug discovery process.

Large-scale phenotypic screens have identified small molecules that display antiviral activity against SARS-CoV-2, but often the proteins targeted by these compounds remain unknown (19). Chemical proteomics has emerged as a powerful approach to facilitate target identification in phenotypic screening assays (20–22), especially when performed with “fully-functionalized” compound libraries that are designed to possess i) electrophilic or photoreactive groups for covalent modification of proteins targets in cells; and ii) latent bioorthogonal affinity handles (e.g., alkynes) for conjugation to reporter tags for protein enrichment and identification.

Here, we perform integrated phenotypic screening and chemical proteomics using a stereochemically defined library of fully-functionalized photoreactive compounds (photo-stereoprobes) to identify and characterize small molecules that stereoselectively inhibit SARS-CoV-2 infection in human cells. The hit photo-stereoprobes were found to engage the translation termination factor ETF1 with a structure-activity relationship matching their antiviral activity. We further found that the active photo-stereoprobes perturb viral programmed ribosomal frameshifting (PRF) without causing ETF1 degradation or gross host cell cytotoxicity. Finally, we show that the active stereoprobes inhibited the replication of multiple viruses from different families that also utilize PRF mechanisms for protein production.

## RESULTS

### Phenotypic screening identifies photo-stereoprobes that inhibit SARS-CoV-2 infection

We have previously generated chemical proteomic ligandability maps of both fragment (23) and elaborated (24) photo-stereoprobes, which revealed that these compounds interact with a diverse array of proteins in human cells. The elaborated photo-stereoprobes were designed to contain structurally distinct cores (azetidine, pyrrolidine, tryptoline) with two stereocenters (to give a total of four stereoisomers per chemotype) and to incorporate a diazirine and alkyne handle for UV light-mediated crosslinking to interacting proteins and affinity enrichment via copper-catalyzed azide-alkyne cycloaddition (CuAAC or click (25, 26)) chemistry to a biotin tag, respectively.

We surmised that the larger size and structural diversity of the elaborated photo-stereoprobes would make them better suited for phenotypic screening compared to original fragment photo-stereoprobes, and we accordingly performed a screen of an in-house 56-member elaborated photo-stereoprobe library to identify compounds that inhibit SARS-CoV-2 infection (Fig. 1A-B). Human Calu-3 lung epithelial cells were pre-treated with photo-stereoprobes at 20 µM for 3 h, inoculated with the SARS-CoV-2 (strain WA1/2020) at an MOI of 0.1, and viral supernatant titers were assessed 48 h post infection with a focus-forming assay. In parallel, photo-stereoprobes were assessed for cytotoxicity 48 h post-treatment using a luminescence-based cell viability assay and compounds that reduced cell viability by >20% were deprioritized. From this screen, we identified two compounds that production of infectious SARS-CoV-2 progeny in human lung epithelial cells by at least one order of magnitude, with one hit (WX-02-31) exhibiting activity similar to the positive control compound EIDD-2801 (molnupiravir) (Fig. 1C).

**Figure 1.**
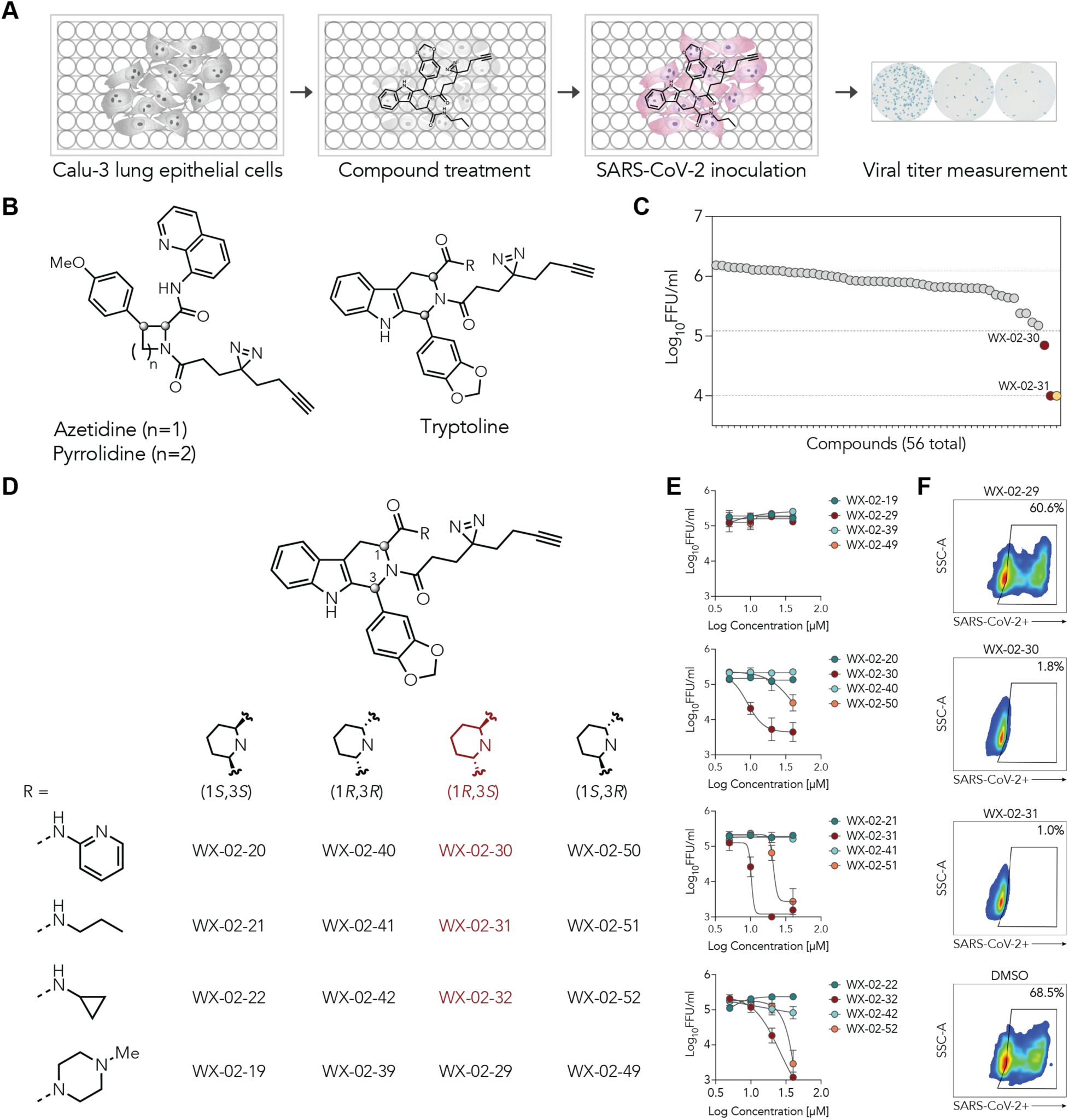
Phenotypic screen to identify photo-stereoprobes that inhibit SARS-CoV-2 infection. (A) Schematic for a phenotypic screen to identify small molecule modulators of SARS-CoV-2 infection. Calu-3 cells were pre-treated with photo-stereoprobes (3 h), inoculated with SARS-CoV-2 (MOI 0.1) in the presence of compounds, and SARS-CoV-2 titers were assessed using a focus-forming assay 48 h post infection. (B) Representative structures of core scaffolds (azetidine, pyrrolidine, and tryptoline) of the photo-stereoprobe library, where R designates variable appendage groups. (C) Screening results, where dotted lines indicate DMSO control (top line), one log_10_-fold reduction in infectivity relative to the DMSO control (middle line), and limit of detection (bottom line). Prioritized photo-stereoprobe hits from the phenotypic screen are shown in red. The positive control antiviral agent EIDD-2801 (molnupiravir) (2) is shown in yellow. Compounds that cause a >20% reduction in cell viability were excluded from the analysis. (D) Structures of photo-stereoprobe hits (red) and their stereoisomers (black). (E) Concentration-dependent inhibition of SARS-CoV-2 infection by photo-stereoprobe hits in Calu-3 cells. Calu-3 cells were pre-treated with indicated concentrations of photo-stereoprobes for 3 h, inoculated with SARS-CoV-2 (MOI 0.1) in the presence of compounds, and viral titers were assessed using a focus-forming assay 48 h post infection. (F) Results from a second SARS-CoV-2 infection model where HeLa-ACE2 cells were pre-treated with the indicated photo-stereoprobes (10 µM, 3 h), inoculated with SARS-CoV-2 in the presence of compounds or DMSO, and viral antigen levels assessed 24 h later by flow cytometry. For (E), data are average values ± SD for 2-3 independent experiments. For (F), data are from a single experiment representative of two independent experiments.

The two hit compounds WX-02-30 and WX-02-31 shared the same core scaffold and (1*R*,3*S*) stereoconfiguration, only differing in their amide appendages (derived from 2-aminopyridine and *n*-propylamine, respectively, Fig. 1D). Secondary concentration-dependent assays confirmed the antiviral activity of WX-02-30 and WX-02-31, with each compound showing half-maximal activity at ∼10 µM. In contrast, the corresponding enantiomeric compounds - WX-02-50 and WX-02-51 - showed lower activity (Fig. 1D-E), and the diastereomers were inactive (Fig. 1D-E). We further tested additional analogs and found that an N-cyclopropyl substitution (WX-02-32) retained stereoselective activity while an N-methypiperazine derivative (WX-02-29) was inactive (Fig. 1D-E). None of the active photo-stereoprobes exhibited cytotoxicity as measured in a luminescence-based cell viability assay (Fig. S1A-B). We also examined the kinetics of viral inhibition for the most potent photo-stereoprobe WX-02-31, its stereoisomers, and the inactive photo-stereoprobe WX-02-29 using multi-step growth curves. WX-02-31, but not WX-02-29, stereoselectively reduced SARS-CoV-2 titers by two and four orders of magnitude at 24 and 48 h post infection, respectively (Fig S1C-D). To confirm that this activity was not limited to Calu-3 cells, we tested the ability of the (1*R*,3*S*) analogs to inhibit SARS-CoV-2 infection in HeLa-ACE2 cells. Consistent with the inhibition seen in Calu-3 cells, the active photo-stereoprobes, WX-02-30, WX-02-31, and WX-02-32, but not WX-02-29, abrogated viral infection in HeLa-ACE2 when assessed by flow cytometry (Fig. 1F and S1E-G).

### Human ETF1 is stereoselectively engaged by antiviral photo-stereoprobes

In anticipation of mapping the relevant protein targets of the active photo-stereoprobes, we first determined whether an analog lacking the diazirine-alkyne tag retained antiviral effects, as such a compound could then be used as a competitor to block the photo-stereoprobe enrichment of proteins in chemical proteomic experiments. However, we found that a compound bearing a propanamide in place of the diazirine-alkyne tag (WX-02-654) did not inhibit SARS-CoV-2 infection in Calu-3 or HeLa-ACE2 cells (Fig. S2A-E). We therefore focused our chemical proteomic experiments on mapping proteins that showed stereoselective enrichment by the active photo-stereoprobes.

We initially assessed the general protein interaction profiles of active photo-stereoprobes and their stereoisomer controls by SDS-PAGE. Calu-3 and HEK293T cells were treated with the photo-stereoprobes (20 µM, 3 h), exposed to UV light, lysed, and stereoprobe-modified proteins were conjugated to a fluorescent AF647-azide tag by CuAAC chemistry and detected by SDS-PAGE and in-gel fluorescence scanning (Fig. S2F-G). Similar overall protein interaction profiles were observed for the active photo-stereoprobes and their respective stereoisomers, indicating that these compounds did not show gross differences in cellular uptake or distribution in cells. We next identified photo-stereoprobe-interacting proteins in Calu-3 cells using multiplexed tandem mass tagging (TMT) proteomics following the general protocol outlined above except that, after cell lysis, stereoprobe-modified proteins were conjugated to a biotin-PEG_4_-azide tag by CuAAC chemistry, enriched with streptavidin-conjugated beads, digested on-bead with trypsin, and the resulting tryptic peptides reacted with TMT tags and analyzed by mass spectrometry (MS)-based proteomics (Dataset S1).

Each active photo-stereoprobe and its stereoisomers exhibited stereoselective binding to discrete sets of proteins as revealed through quadrant plot displays (Fig. 2A and S2H-J). We considered that candidate protein targets of interest should: 1) show substantially higher (> 2.5 fold) signals in cells treated with active-photo-stereoprobes versus their respective stereoisomer controls, and 2) not be stereoselectively enriched by the inactive analog WX-02-29. As another filter to prioritize candidate targets of interest, we performed a more quantitative comparison of the structure-activity relationships (SARs) between the relative protein engagement and inhibitory activity of SARS-CoV-2 infection that included additional structurally related inactive photo-stereoprobes (Fig. S3A-B). We then performed pairwise Pearson correlations between the stereoselective enrichment of proteins quantified by the 19 photo-stereoprobes in proteomics experiments of Calu-3 cells and their respective log_10_-transformed fold-decreases in SARS-CoV-2 infectivity. From this analysis, the translation termination factor ETF1 showed the strongest correlation (r = 0.8) (Fig. 2B and S3B), with this protein displaying clear stereoselective enrichment by each of the active photo-stereoprobes WX-02-30, WX-02-31, and WX-02-32 but not by inactive analogs (Fig. 2C-D). To account for the possibility that the active photo-stereoprobes might additionally engage viral proteins, we performed chemical proteomic experiments in SARS-CoV-2-infected cells (Fig. S4A and Dataset S1). These experiments again showed strong stereoselective interactions between active photo-stereoprobes and ETF1 (Fig. S4B-C) and did not reveal stereoselective binding of these compounds to viral proteins (even though multiple viral proteins were detected as background signals in the proteomic experiments) (Dataset S1).

**Figure 2.**
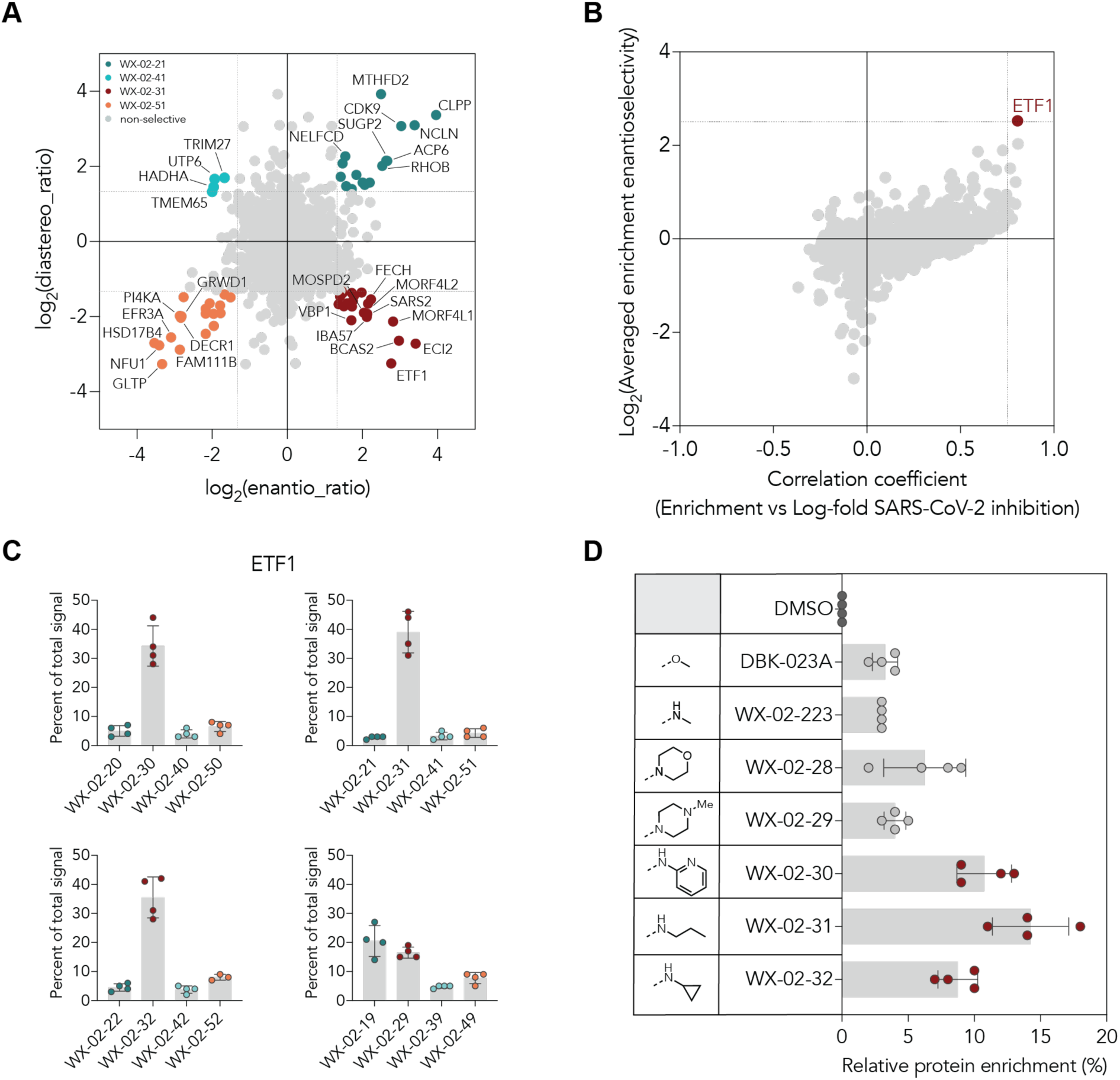
Mass spectrometry (MS)-based proteomic analysis of proteins that interact with photo-stereoprobes in Calu-3 cells. (A) Quadrant plots (24, 55) displaying stereoselectively liganded proteins by the active stereoprobe WX-02-31 compared to stereoisomeric control compounds WX-02-21/-41/-51 (20 µM, 3 h) in Calu-3 cells. The enantio-ratio (x-axis) shows the enrichment ratio value for the stereoisomer with maximum engagement of a protein (probe^max^) compared to its enantiomer. The diastereo-ratio (y-axis) shows the ratio value of probe^max^ to the diastereomer with higher relative engagement. Colored dots: proteins with an enantio-ratio ;= 2.5 and diastereo-ratio ;= 2.5; gray dots: other proteins. Data represent average values from two independent experiments, each with two technical replicates. (B) Correlation plot comparing protein engagement by active photo-stereoprobes and SARS-CoV-2 inhibition activity, revealing the strongest correlation for the protein ETF1. The Pearson correlation coefficient (x-axis) was calculated using the relative protein enrichment in MS-based proteomic experiments and the log _10_-fold decrease value in SARS-CoV-2 infection from Fig. 1C (see Fig. S3 for further details on this calculation). The average enrichment enantioselectivity (y-axis) is defined as the mean of the log_2_-transformed enantio-ratios of the three active photo-stereoprobes (WX-02-30, WX-02-31, and WX-02-32) compared to their respective enantiomeric inactive controls (WX-02-50, WX-02-51, and WX-02-52). Only proteins quantified in two independent proteomic experiments are shown. (C-D) Bar graphs showing the MS-based proteomic enrichment profile for ETF1 by active photo-stereoprobes (WX-02-30, WX-02-31, and WX-02-32) relative to their stereoisomeric controls (20 µM, 3 h) (C) or relative to inactive analogs of the same (1*R*, 3*S*) stereoconfiguration (D) in Calu-3 cells. The inactive analog WX-02-29 is shown in both C-D. For (C-D), data represent average values for two independent experiments.

### Recombinant, purified ETF1 binds photo-stereoprobes

To verify the stereoselective binding of active photo-stereoprobes to ETF1, we recombinantly expressed an epitope (FLAG)-tagged version of ETF1 in HEK293T cells by transient transfection. The cells were treated with photo-stereoprobes (20 µM, 3 h), UV crosslinked, conjugated to an AF647-azide by CuAAC chemistry, and analyzed by SDS-PAGE and in-gel fluorescence scanning. A clear signal was observed at the predicted MW of ETF1 (∼50 kDa) in ETF1-transfected cells treated with active photo-stereoprobes WX-02-30, −31, and −32, but not with inactive stereoprobe controls (stereoisomers or WX-02-29) (Fig. 3A and S5A). We also observed clear increases in fluorescent signals for recombinant ETF1 across a broad concentration range of WX-02-31 (2.5-20 µM) (Fig. 3B and S5B-C), indicating that the photo-stereoprobe-ETF1 interaction was low affinity. Nonetheless, the stereoselectivity of the WX-02-31-ETF1 interaction was preserved at the highest test concentration compared to the enantiomer WX-02-51. We additionally found that co-treatment with the propanamide analog WX-02-654 (40 µM) failed to block interactions between WX-02-31 and recombinant ETF1 (Fig. S5D-E). We finally expressed and purified human ETF1 from an *E. coli* expression system and confirmed that the purified protein also stereoselectively interacted with WX-02-30, WX-02-31, and WX-02-32, but not WX-02-29 (Fig. 3C-D).

**Figure 3.**
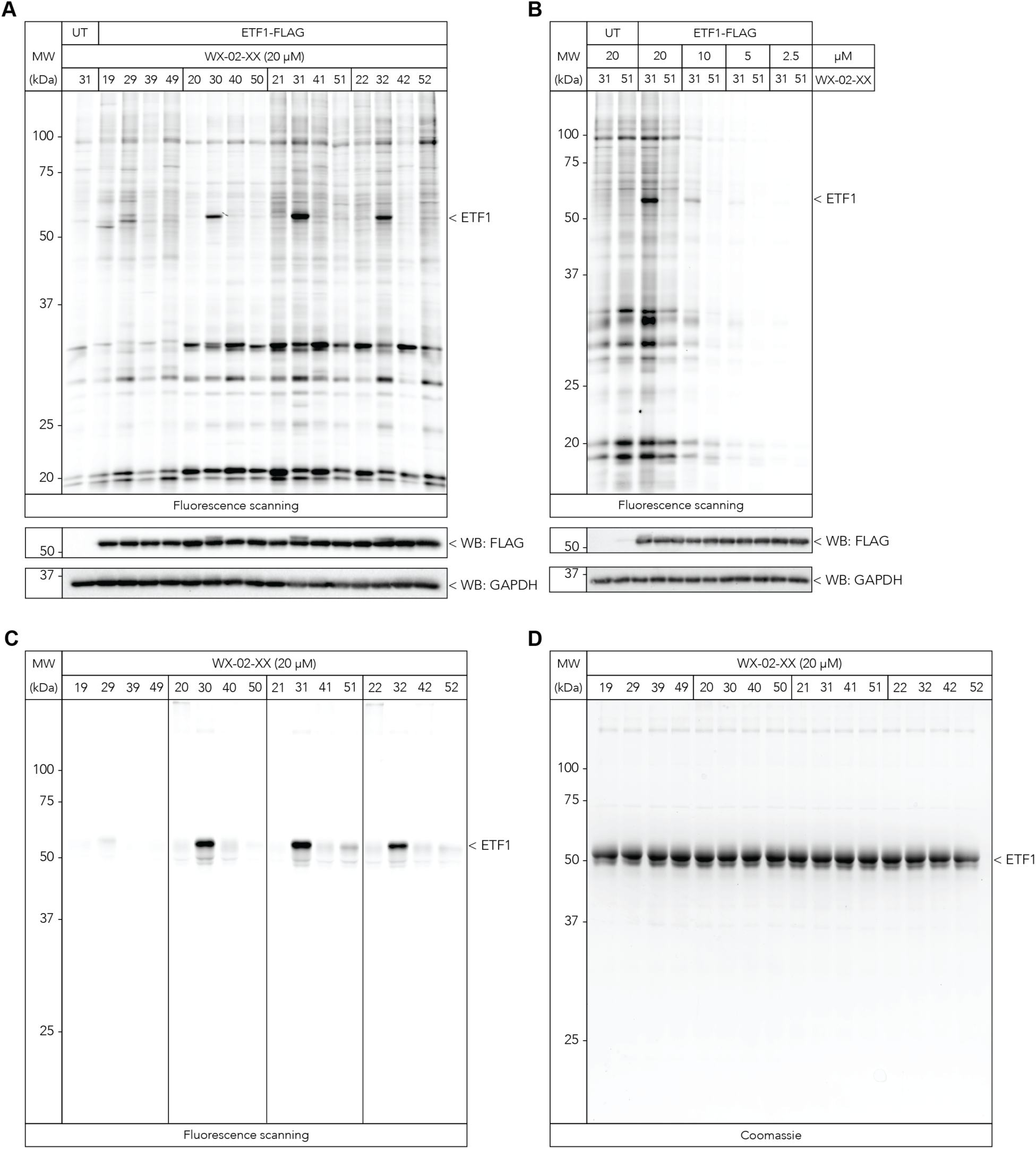
Confirmation of stereoselective binding of hit photo-stereoprobes to recombinant and purified ETF1. (A, B) Fluorescent gel (top) and western blot (bottom) images showing stereoselective engagement of recombinant FLAG epitope-tagged ETF1 in transiently transfected HEK293T cells by the active photo-stereoprobes WX-02-30, WX-02-31, and WX-02-32 (20 µM, 3 h, (A); indicated concentrations of WX-02-31, 3 h (B)). Anti-FLAG and anti-GAPDH western blot images confirm expression of recombinant FLAG epitope-tagged ETF1 and protein loading. UT, untransfected control cells. After stereoprobe treatment and UV light exposure, cells were lysed and stereoprobe-modified proteins conjugated to an AF647-azide tag reporter tag by CuAAC chemistry and analyzed by SDS-PAGE and in-gel fluorescence scanning. (C) Fluorescent gel image showing stereoselective engagement of recombinant ETF1 expressed and purified from *E. coli* by the active photo-stereoprobes WX-02-30, WX-02-31, and WX-02-32 (20 µM, 3 h,). Samples were processed as described in (A). (D) Coomassie blue stained gel showing purified ETF1 protein evaluated in (C). (A-D), data are representative of two independent experiments.

Taken together, these results indicate that the active photo-stereoprobes directly bind to purified ETF1 protein with an SAR that matches their antiviral activity, and these interactions do not require the presence of additional proteins.

### Active photo-stereoprobes modulate viral programmed ribosomal frameshifting

As ETF1 has previously been reported to modulate the translation of viral proteins (27–29), we tested whether the active photo-stereoprobes modulated programmed ribosomal frameshifting (PRF), a process upon which SARS-CoV-2 and other viruses (e.g., HIV) depend for replication (30–33). We generated a stable HeLa PRF reporter cell line with a construct containing mCherry, the SARS-CoV-2 frameshifting nucleotide element, and GFP in the −1 reading frame [mCherry-SARS2-FSE-GFP(−1)] (Fig. 4A). We also generated two additional reporter cell lines without the frameshifting nucleotide element [mCherry-FSE-GFP(No-FSE)] and a stop codon immediately after mCherry [mCherry-FSE-GFP(0)], which would allow for comparing compound effects on PRF versus general protein translation (Fig. 4A). The cell models were then treated with active and inactive control photo-stereoprobes and the ratio of GFP to mCherry assessed 48 h post-treatment. As a positive control, we included merafloxacin, a previously reported inhibitor of PRF and SARS-CoV-2 infection (34). The active photo-stereoprobes WX-02-31 and WX-02-32, but not their respective stereoisomers or the inactive compound WX-02-29, increased PRF in the mCherrry-FSE-GFP(−1) cell line (Fig. 4B and S6A), while producing minimal effects on general translation as measured in the mCherry-FSE-GFP(No-FSE) and mCherry-FSE-GFP(0) cell lines (Fig. S6B-C). Finally, we generated an additional HeLa reporter cell line containing a construct with mCherry, the HIV-1 frameshifting nucleotide element located between the Gag-Pol proteins, and GFP in the −1 reading frame [mCherry-HIV-FSE-GFP(−1)] and found that, in these cells, the active photo-stereoprobe WX-02-31, but not its stereoisomers, increased PRF (Fig. 4C). These data, taken together, indicate that the active photo-stereoprobes may produce antiviral effects, at least in part, through disrupting PRF.

**Figure 4.**
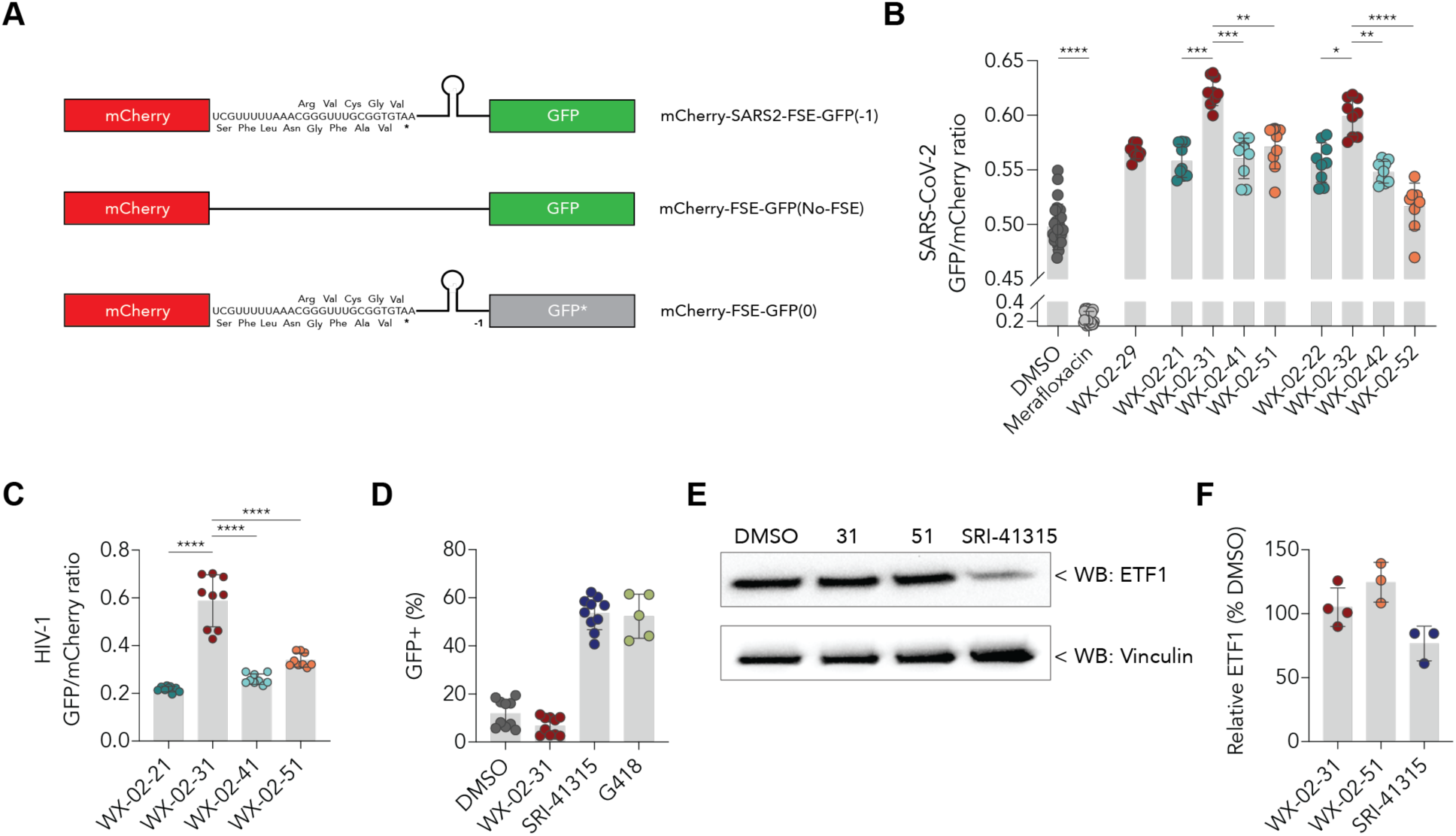
Photo-stereoprobes modulate SARS-CoV-2 programmed ribosomal frameshifting (PRF). (A) Schematic of the dual-reporter constructs used to examine SARS-CoV-2 PRF. mCherry and GFP are co-expressed upon frameshifting. Top: mCherry-SARS2-FSE-GFP(−1) which contains mCherry in the 0 reading frame followed by the SARS-CoV-2 PRF sequence and GFP in the −1 reading frame; middle: mCherry-FSE-GFP(No-FSE), which continuously translates mCherry and GFP; bottom: mCherry-FSE-GFP(0), which shifts GFP back to the 0 frame after the SARS-CoV-2 PRF sequence thus resulting in a stop codon after mCherry. (B) Effects of active photo-stereoprobes WX-02-31 and WX-02-32 (20 µM, 48 h) versus stereoisomeric compound, inactive compound (WX-02-29), and DMSO controls on HeLa cells stably expressing the mCherry-SARS2-FSE-GFP(−1) construct. Merafloxacin, an established inhibitor of SARS-CoV-2 PRF, was included as a positive control. Data are average values ± SD from three independent experiments. (C) Effects of active photo-stereoprobe WX-02-31 (20 µM, 48 h) versus stereoisomeric compound controls on HeLa cells stably expressing an mCherry-HIV-FSE-GFP(−1) which contains mCherry in the 0 reading frame followed by the HIV-1 PRF sequence and GFP in the −1 reading frame. Data are average values ± SD from three independent experiments. (D) Effect of active photo-stereoprobe WX-02-31 (20 µM, 24 h) on translational readthrough measured in HEK293T cells expressing a GFP construct with a premature termination codon at Trp57. Results for positive control compounds SRI-41315 (20 µM) and G418 (600 µM) are also shown. Data are average values ± SD for 2-4 independent experiments. (E) Western blot analysis of ETF1 protein in Calu-3 cells following treatment with indicated compounds (20 µM, 24 h). Data are from a single experiment representative of two independent experiments. (F) MS-based proteomic quantification of ETF1 abundance in Calu-3 cells treated with the indicated compounds (20 µM, 24 h) relative to DMSO control. Data are average values ± SD for 3-4 independent experiments.

### WX-02-31 acts differently than other ETF1 ligands

Previous phenotypic screens seeking to identify compounds that promote nonsense suppression, or translational readthrough, have identified additional ETF1 ligands (35, 36). We tested one of these compounds SRI-41315 and found that it increased PRF in the SARS-CoV-2 reporter cell model, albeit to a lesser extent than WX-02-31 (Fig. S6D). We then asked whether WX-02-31, like SRI-41315, promoted translational readthrough of premature termination codons (PTC) using a HEK293T cell model containing a GFP reporter plasmid with a stop codon at position 57 (37). Interestingly, WX-02-31 did not exhibit translational readthrough activity in this model, in contrast to the activity displayed by SRI-41315 and the aminoglycoside antibiotic G418 (38) (Fig. 4D).

The distinct activities of WX-02-31 and SRI-41315 in assays of translational frameshifting and readthrough suggested these compounds may differentially interact with ETF1. Also consistent with this hypothesis, recent cryo-EM work has revealed that SRI-41315 acts as a metal-dependent molecular glue binding to a composite pocket formed by ETF1 and the ribosome (39), while we had found, as shown above (Fig. 3C-D), that WX-02-31 retains stereoselective binding to recombinant, purified ETF1. Upon binding to the ETF1-ribosome complex, SRI-41315 and other previously described ligands promote the degradation of ETF1 through an E3 ligase-proteasome mechanism (36, 40). We found, however, that WX-02-31, unlike SRI-41315, did not reduce ETF1 protein content of Calu-3 cells as measured by western blotting (Fig. 4E) or MS-based proteomics (Fig. 4F). We therefore conclude that WX-02-31 binding to ETF1 does not promote the degradation of this protein in cells.

### Active photo-stereoprobes inhibit additional viruses that depend on PRF

We next tested the effects of active photo-stereoprobes in models for additional viruses that display variable dependency on frameshifting mechanisms. HIV-1 depends on PRF to generate Gag, which encodes viral structural proteins, and Pol, which encodes viral enzymes (30), and we found, using an HIV-1 NL4-3 luciferase pseudovirus reporter system (41), that WX-02-31, but not its stereoisomers or WX-02-29, reduced HIV-1 antigen expression (Fig 5A). A similar pharmacological profile was observed in infection assays for Sindbis virus, which uses frameshifting to produce the TF protein in lieu of the 6K protein (42) (Fig. 5B and S7A). Porcine reproductive and respiratory system virus (PRRSV) contains multiple PRF regions in the viral genome and mutation of these regions decreases PRRSV replication (43, 44). We again found that WX-02-31, but not inactive control compounds, inhibited PRRSV replication using a GFP reporter virus (Fig 5C and S7B). Finally, we also analyzed viruses that do not have known non-canonical translation mechanisms, including lymphocytic choriomeningitis virus (LCMV) and influenza A virus (IAV), and WX-02-31 did not inhibit the infectivity of these viruses (Figs 5D-E and S7C-E).

**Figure 5.**
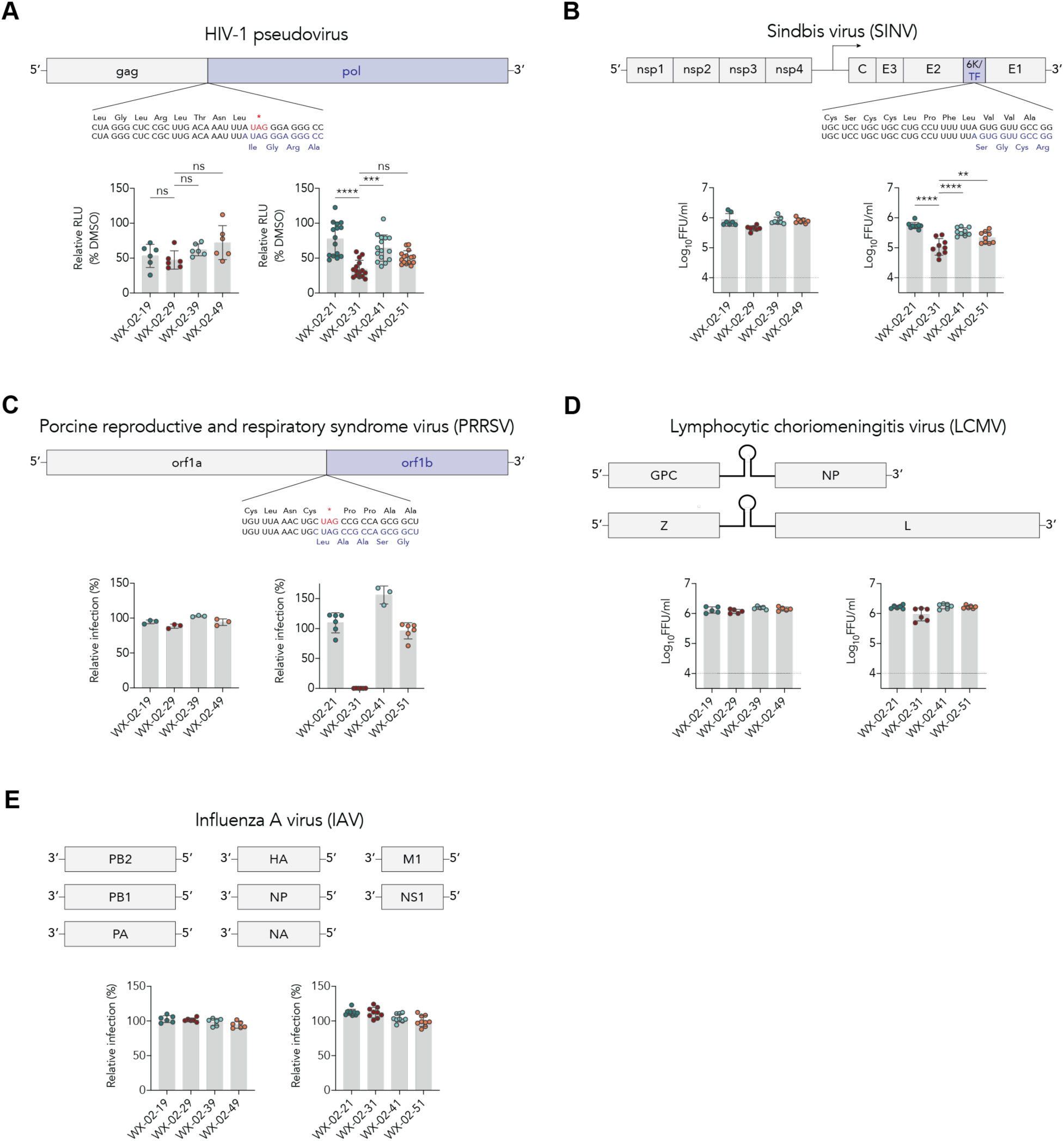
Photo-stereoprobes inhibit additional viruses that use PRF. (A) Effect of active stereoprobe WX-02-31 and control stereoprobes on HIV-1 pseudovirus production. Top, schematic of the HIV-1 ribosomal frameshift sequence. Bottom, HEK293T cells expressing the HIV-1 NL4-3 luciferase vector were treated with the indicated photo-stereoprobes (10 µM) and assessed for luciferase activity after 24 h (bottom). Data are average values ± SD for 2-5 independent experiments that each have three technical replicates. (B) Effect of active stereoprobe WX-02-31 and control stereoprobes on Sindbis virus (SINV) infection. Top, schematic of the SINV ribosomal frameshift sequence. Bottom, HeLa cells were pre-treated with indicated photo-stereoprobes (10 µM, 3 h), inoculated with SINV (MOI 3) in the presence of compounds, and viral supernatant titers were assessed after 48 hpi (bottom). Data are average values ± SD for three independent experiments that each have 2-3 technical replicates. (C) Effect of active stereoprobe WX-02-31 and control stereoprobes on porcine reproductive and respiratory syndrome virus (PRRSV) infection. Top, schematic of the PRRSV ribosomal frameshift sequence. Bottom, MA104 cells were pre-treated with indicated photo-stereoprobes (10 µM, 3 h), inoculated with PRRSV-GFP (MOI 0.1) in the presence of compounds, and viral antigen levels were assessed by GFP expression after 24 hpi (bottom). Data are representative of two independent experiments that each have three technical replicates. (D) Effect of active stereoprobe WX-02-31 and control stereoprobes on lymphocytic choriomeningitis virus (LCMV) infection. Top, schematic of the LCMV genome. Bottom, HeLa cells were pre-treated with indicated photo-stereoprobes (10 µM, 3 h), inoculated with LCMV (clone 13, MOI 0.01) in the presence of compounds and viral supernatant titers were assessed after 48 hpi (bottom). Data are average values ± SD for two independent experiments that each have 2-3 technical replicates. (E) Effect of active stereoprobe WX-02-31 and control stereoprobes on influenza A virus (IAV) infection. Top, schematic of the IAV genome. Bottom, Calu-3 cells were pre-treated with indicated photo-stereoprobes (20 µM, 3 h), inoculated with IAV (MOI 0.005) in the presence of compounds, and viral antigen levels were assessed by flow cytometry after 48 hpi (bottom). Data are the average values ± SD for two (left panel) and five (right panel) independent experiments that each have three technical replicates. For (A-E), one-way ANOVA with multiple comparisons: *, p < 0.05, **, p <0.01, ***, p < 0.001; ****, p < 0.0001; ns, not significant.

## DISCUSSION

Chemical proteomics has made substantive contributions to the identification of protein targets responsible for the biological effects of small molecules in phenotypic screens (20). Fully functionalized compound libraries that bear, for instance, photoreactive and clickable groups have an additional advantage for phenotypic screening in that they do not require chemical modification to proceed from screening hits to target identification (24, 45–47). For our SARS-CoV-2 infectivity screen, we leveraged a recently described class of elaborated photo-stereoprobes that incorporate defined stereocenters into the fully functionalized library design to facilitate the mapping of protein targets of compounds that display stereoselective phenotypic effects (24). Although this elaborated photo-stereoprobe library was small in size (56 compounds), it delivered a structurally related set of hit tryptoline compounds that suppressed SARS-CoV-2 replication. This same focused elaborated photo-stereoprobe library also identified structurally and stereochemically distinct hit compounds in an autophagy screen (24). We are unsure why such a small set of elaborated photo-stereoprobes has provided hit compounds in multiple phenotypic screens, but we anticipate that the library performance would be further enhanced by continuing to build out its core and appendage diversity.

The totality of our chemical proteomic and biological data point to ETF1 as a mechanistically relevant target for the antiviral effects of the tryptoline hit compounds. Previous research also supports this conclusion, as ETF1 has been implicated in HIV-1 infection through regulating PRF (28, 48). Nonetheless, how the tryptoline photo-stereoprobes alter ETF1 function to disrupt PRF remains unclear. Our findings indicate that, unlike a previously described ETF1 ligand SRI-41315, the tryptoline photo-stereoprobes do not cause degradation of ETF1 in cells. The tryptoline photo-stereoprobes also directly bind to purified ETF1, which would suggest a different mode of interaction from SRI-41315, which interacts with a composite pocket formed by ETF1 and the ribosome. Regardless of their precise mode of binding to ETF1, our data indicate that the tryptoline photo-stereoprobes are weak ligands, in that they show stereoselective increases in interaction with ETF1 across a concentration of 2-20 µM in cells. Future studies aimed at improving the binding affinity of compounds like WX-02-31 for ETF1 should help to clarify if weak binding is beneficial to the antiviral mechanism of action of the tryptoline photo-stereoprobes by, for instance, limiting the general cellular toxicity that might be an expected property of higher affinity ETF1 ligands (ETF1 has been identified as a common essential protein in large-scale genetic disruption screens (https://depmap.org/portal) .

PRF is a noncanonical translation mechanism utilized by viruses to regulate gene expression and to encode multiple open reading frames (ORFs) through controlled ribosomal slippage onto an alternative reading frame (49). PRF is a highly regulated process, and minor changes substantially affect viral translation and infectivity as seen with HIV-1 and SARS-CoV (50–52). For instance, a modest two-fold increase in PRF has been reported reduced HIV-1 infectivity by 250-1,000-fold and abrogated the production of infectious HIV-1 or MMLV viral particles (52–54). Subtle changes in PRF efficiency also affect coronavirus replication and infection (50). These genetic studies indicate that small perturbations to PRF can have large phenotypic effects on viral replication and raise the provocative question whether such effects are pharmacologically achievable in the absence of gross perturbations in host protein translation and cell physiology. While this hypothesis may be supported by the lack of cytotoxicity observed for the antiviral tryptoline photo-stereoprobes discovered herein, more work is needed to understand the extent to which these compounds might impact protein translational processes beyond PRF. In summary, we have used integrated phenotypic screening and chemical proteomics to identify compounds that stereoselectively suppress viral replication and translation through a mechanism that appears to involve binding to the translation termination factor ETF1 and corresponding perturbation of viral frameshifting. These compounds differentiate from previously described ligands for ETF1 by binding to the protein in the absence of associated ribosomes and sparing its degradation in cells. Our work thus provides an additional chemical tool for studying translational processes regulated by ETF1 and points to the potential for tailoring the pharmacological effects of small molecule-ETF1 interactions to address a wide range of pathogenic viruses.

## Supporting information

Synthetic Chemistry Procedures and Analytical Data

---

**Figure S1.**
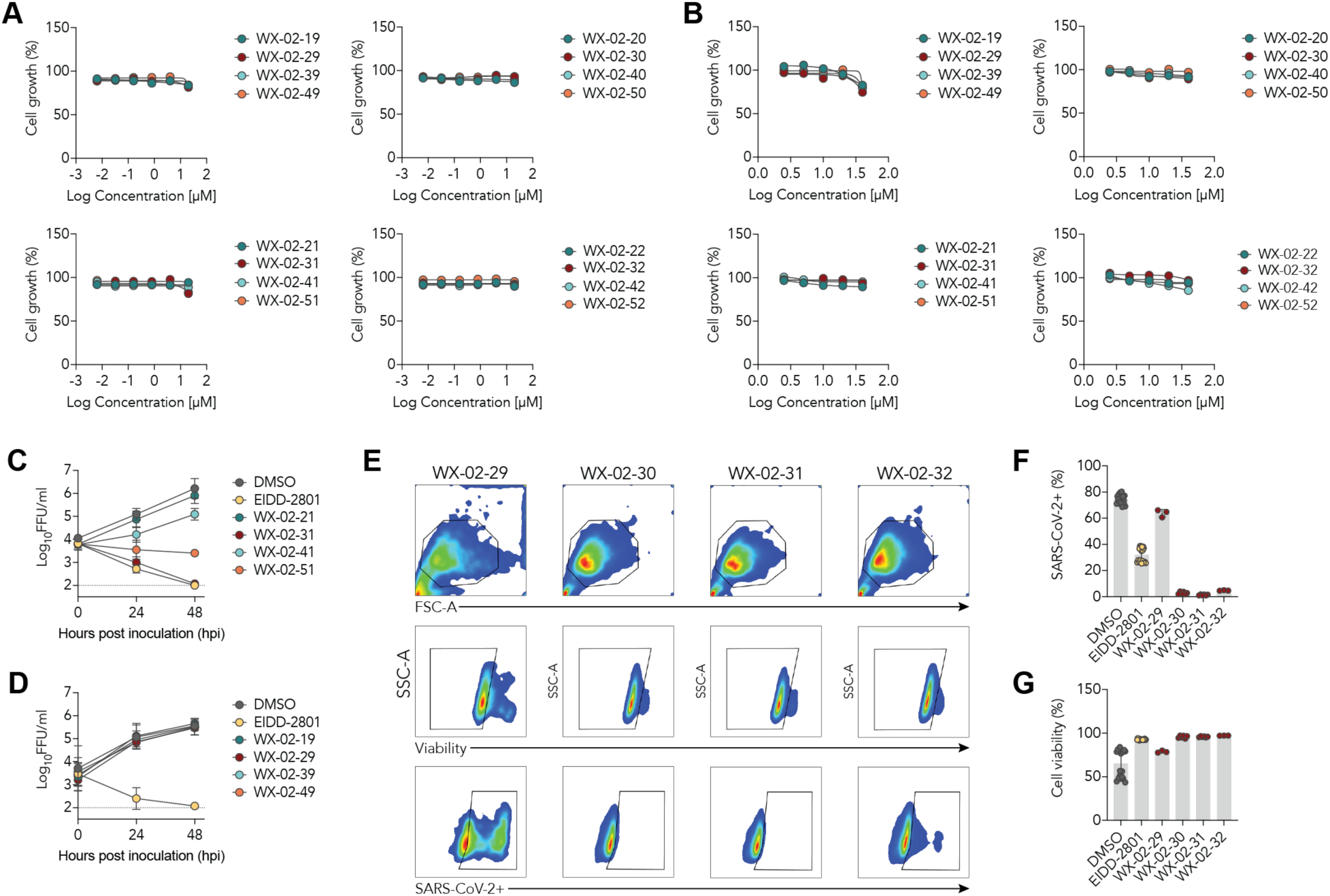
Further characterization of photo-stereoprobes that inhibit SARS-CoV-2 infection. (A-B) Effects of indicated photo-stereoprobes on the growth of Calu-3 (A) and 3T3 (B) cells 48 h post treatment by CellTiter-Glo luminescent cell viability assay. (C-D) Time-dependent effects of the indicated photo-stereoprobes (20 µM) on SARS-CoV-2 infection in Calu-3 cells. Cells were pre-treated with photo-stereoprobes for 3 h and then inoculated with SARS-CoV-2 and viral titers were measured by focus-forming assay after 24 and 48 hpi. The positive control SARS-CoV-2 inhibitor EIDD-2801 is also shown. Data are average values ± SD for two independent experiments. (E) Flow cytometry gating scheme for SARS-CoV-2 viral antigen levels in HeLa-ACE2 cells treated with photo-stereoprobes (10 µM) and analyzed after 24 h (related to Fig. 1F). (F-G) Effects of indicated photo-stereoprobes (10 µM) on SARS-CoV-2 antigen (F) and cell growth (G) in HeLa-ACE2 cells. Cells were pre-treated with photo-stereoprobes (10 µM) for 3 h and then inoculated with SARS-CoV-2 and viral antigen (F) and cell viability (G) were assessed by flow cytometry after 24 h. Data are average values ± SD for two independent experiments.

**Figure S2.**
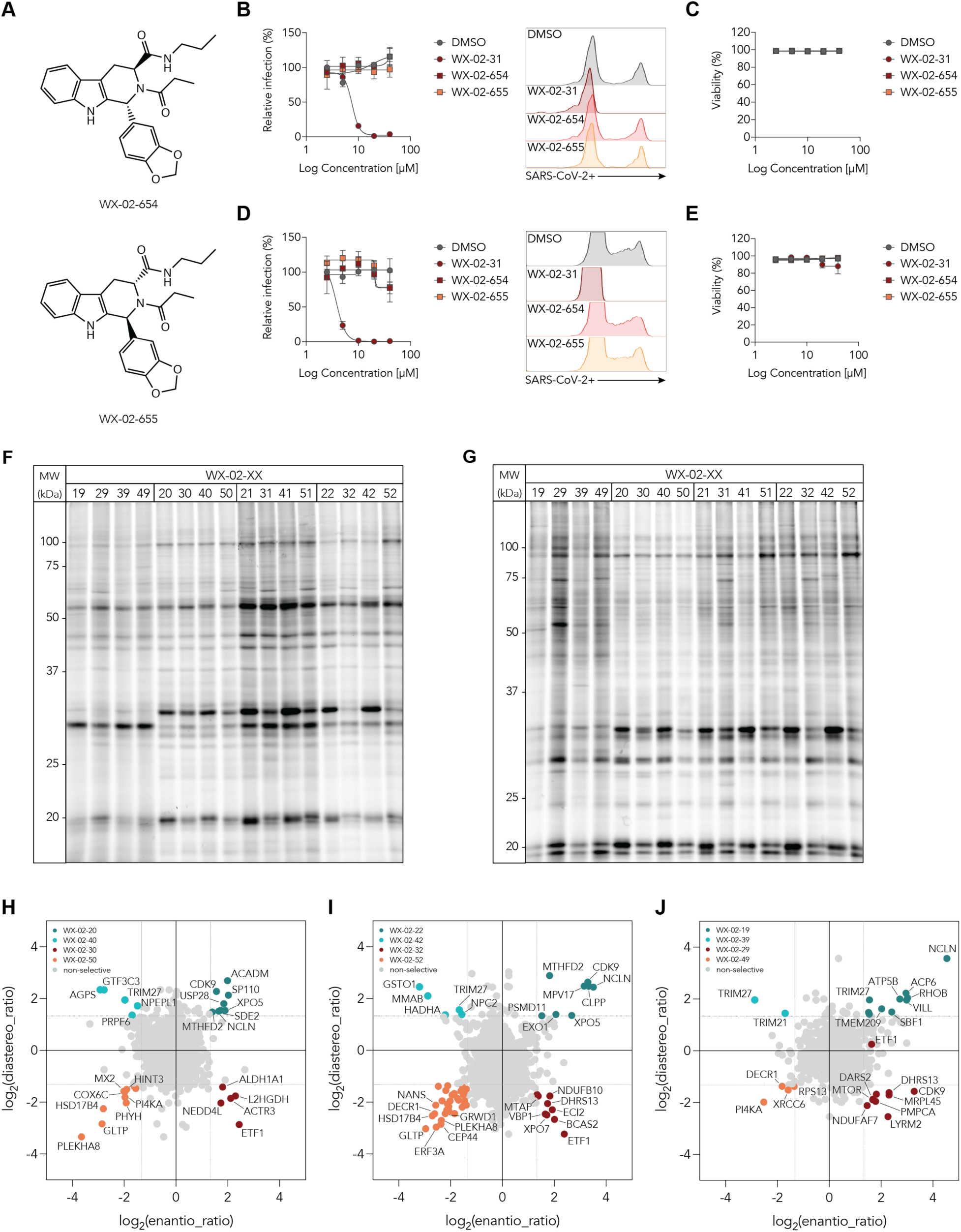
Further MS-based proteomic analysis of proteins that interact with photo-stereoprobes in Calu-3 cells. (A) Structures of non-alkyne analogs of WX-02-31 (WX-02-654) and WX-02-51 (WX-02-655). (B-E) Concentration-dependent effects of WX-02-654 and WX-02-655 on SARS-CoV-2 infection and cell growth in Calu-3 (B-C) and HeLa-ACE2 (D-E) cells. Cells were pre-treated with indicated concentrations of photo-stereoprobes for 3 h and then inoculated with SARS-CoV-2 and viral antigen (B and D) and cell viability (C and E) were assessed by flow cytometry after 24 h. Data are average values ± SD for two independent experiments. Representative flow cytometry histograms are shown from two independent experiments (B and D, right panels). (F-G) Fluorescent gel images showing photo-stereoprobe-protein interactions in Calu-3 (F) and HEK293T (G) cells. Cells were treated with photo-stereoprobes (20 µM) for 3 h and then analyzed as described in Fig. 3A-B. Data are from a single experiment representative of two independent experiments. (H-J) Quadrant plots displaying stereoselectively liganded proteins for WX-02-20 (H), WX-02-22 (I), and WX-02-19 (J) photo-stereoprobe sets in Calu-3 cells (20 µM, 3 h). The enantio-ratio (x-axis) is the enrichment ratio of the stereoisomer showing maximum engagement of a protein (probe^max^) to its enantiomer. The diastereo-ratio (y-axis) is ratio of probe^max^ to the diastereomer with higher relative engagement. Colored dots: proteins with an enantio-ratio ;= 2.5 and diastereo-ratio ;= 2.5; gray dots: non-selective proteins. Data are the average values from two independent experiments.

**Figure S3.**
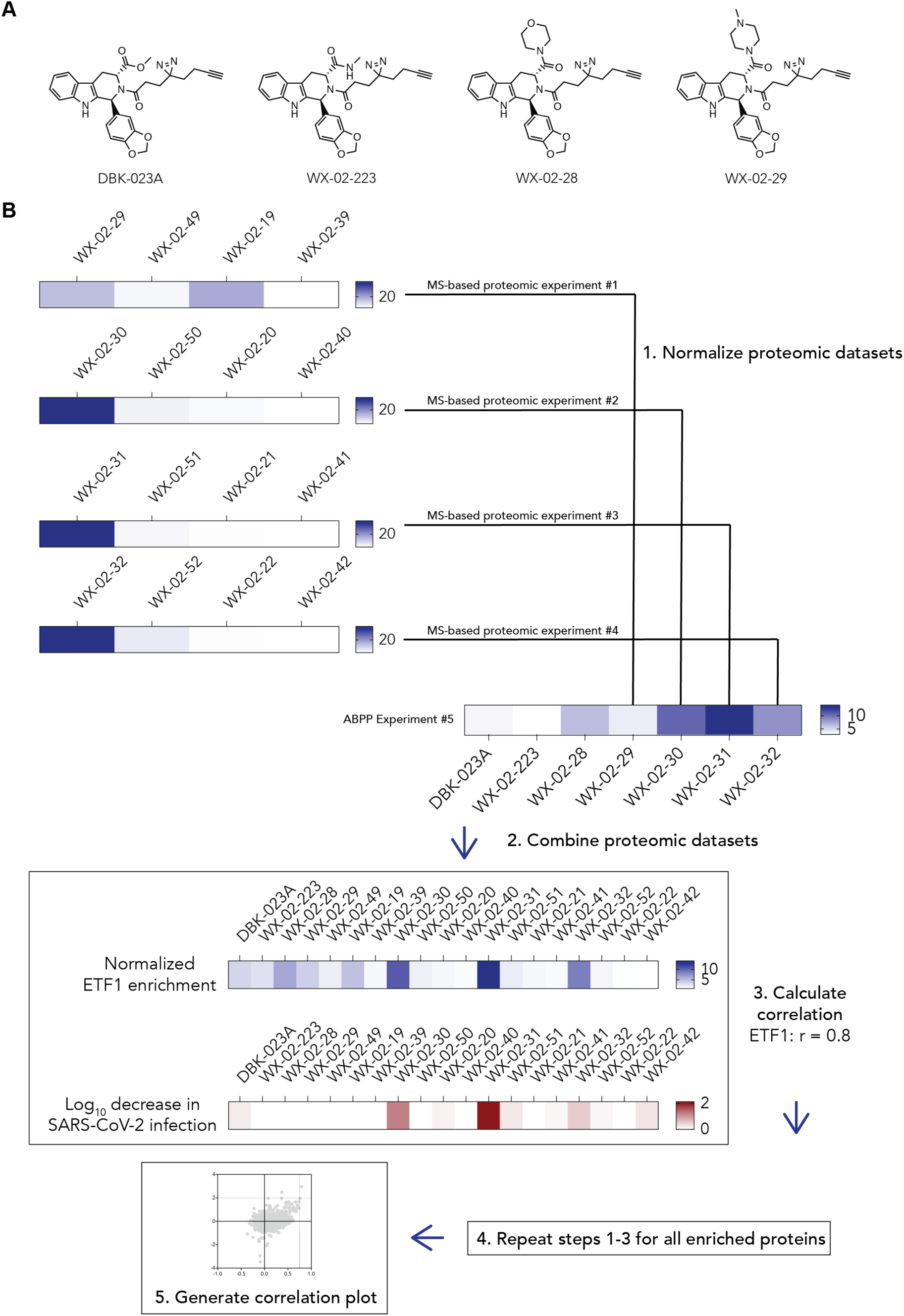
Workflow for the correlation analysis between photo-stereoprobe enrichment of proteins and inhibition of SARS-CoV-2 infection. (A) Structures of the inactive (1*R*, 3*S*) photo-stereoprobes evaluated for interacting proteins by MS-based proteomics. (B) Workflow for correlation of binding and antiviral activity profiles of the indicated photo-stereoprobes. For each protein quantified by MS-based proteomic experiments: 1) the data in Experiments #1-4 were normalized using as reference the condition in common with Experiment #5 (e.g., WX-02-29 was used as reference to normalize Experiment #1 data); 2) normalized proteomic data from Experiments #1-#4 were combined with data from Experiment #5; 3) the Pearson correlation between protein enrichment and decrease in SARS-CoV-2 infection (log_10_-fold) was calculated. A correlation plot (Fig. 2B) was generated considering data for proteins quantified in > 80% of treatment conditions.

**Figure S4.**
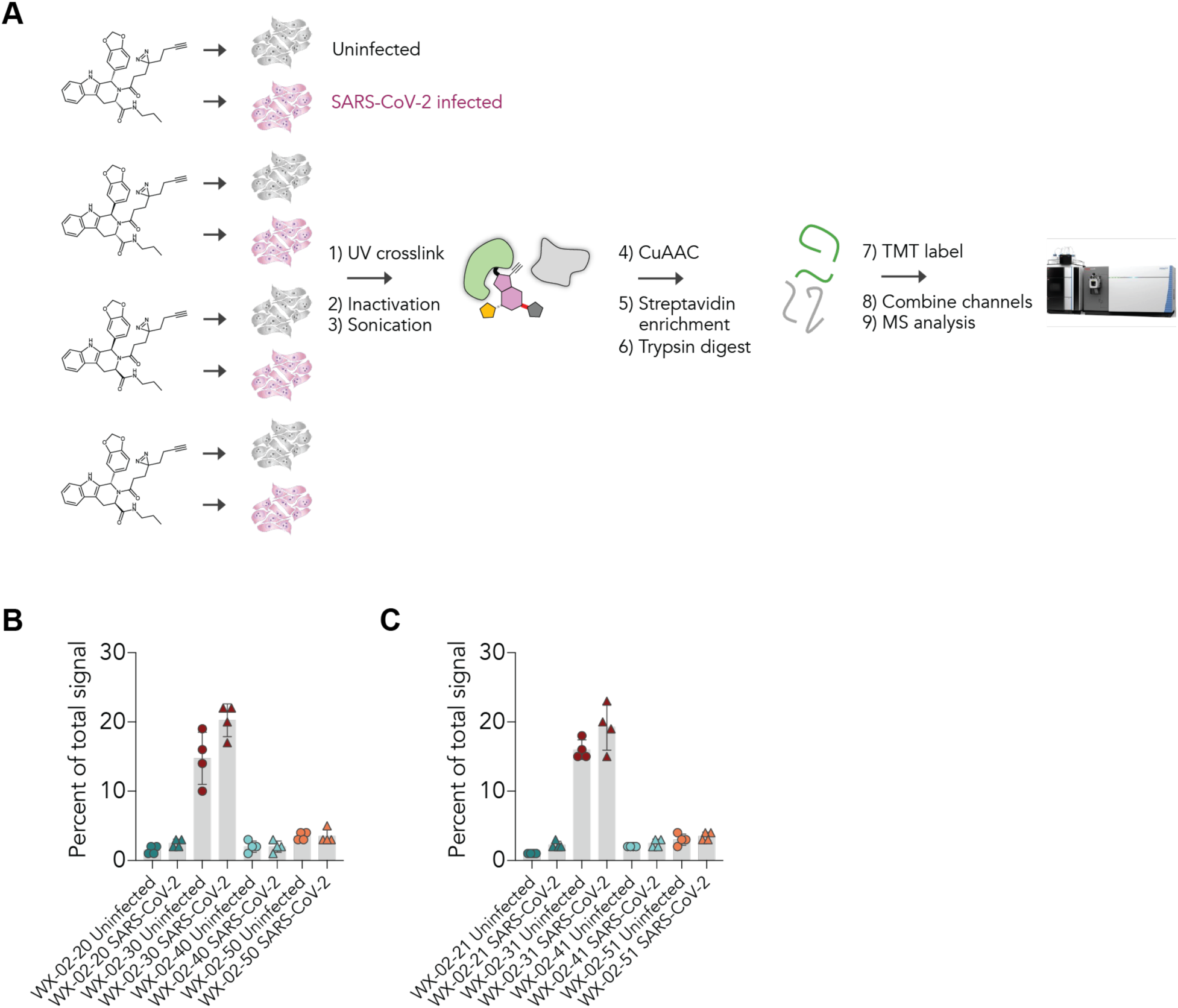
MS-based proteomic analysis of proteins that interact with photo-stereoprobe in SARS-CoV-2-infected Calu-3 cells. (A) Schematic of MS-based proteomic experiments performed on uninfected (gray) or SARS-CoV-2 infected (pink) Calu-3 cells. Uninfected and infected cells (24 h post infection) were treated with stereoprobes for 3 h and analyzed by MS-based proteomics following previously described protocols (24). For SARS-CoV-2 infected cells, cell pellets were subjected to 1% Triton X-100 treatment to inactivate virus prior to sonication. (B-C) Enrichment profile for ETF1 by the WX-02-20 (B) and WX-02-21 (C) photo-stereoprobe compared to their respective stereoisomeric controls (20 µM, 3 h) in uninfected and SARS-CoV-2 infected Calu-3 cells. For (B-C), data are average values ± SD for two independent experiments.

**Figure S5.**
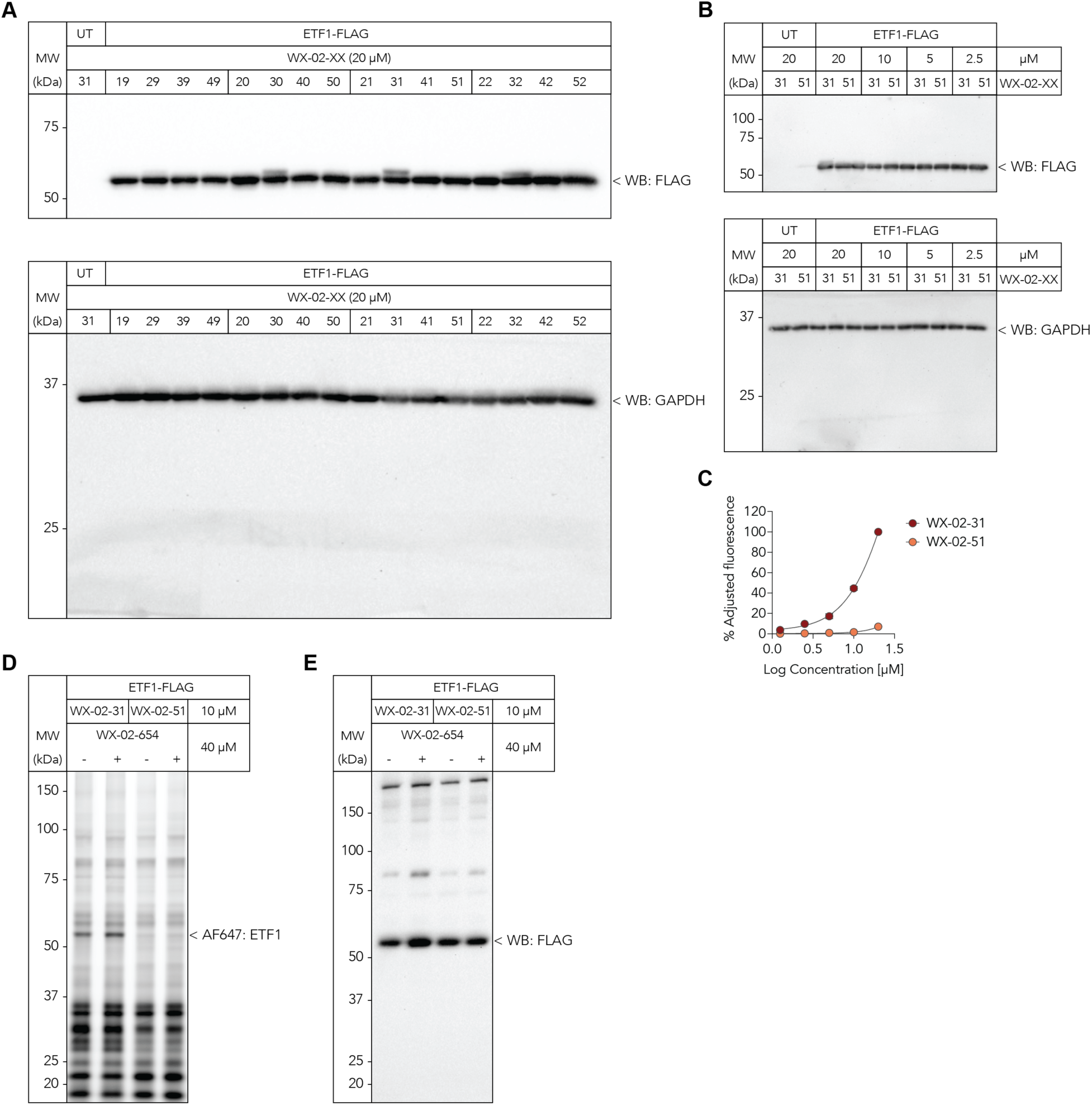
Stereoselective engagement of ETF1 by photo-stereoprobes WX-02-31 and lack of competition of this engagement by WX-02-654. (A-B) Anti-FLAG and anti-GAPDH western blot images confirm expression of recombinant FLAG epitope-tagged ETF1 (top) and protein loading (bottom) (related to Fig. 3A-B). (C) Quantification of fluorescent gel data in Fig. 3B showing concentration-dependent engagement of recombinant FLAG epitope-tagged ETF1 in transfected HEK293T cells treated with the WX-02-31 or WX-02-51 (20 µM, 3 h). Data are average values ± SD for two independent experiments. (D-E) Fluorescent gel image (D) showing lack of competition of WX-02-31 engagement of recombinant FLAG epitope-tagged ETF1 in transfected HEK293T cells by the non-alkyne photo-stereoprobe analog WX-02-654. Cells were pre-treated with WX-02-654 (40 µM, 2 h) followed by WX-02-31 (10 µM, 1 h). Anti-FLAG western blot image (E) confirming expression of recombinant FLAG epitope-tagged ETF1. Data are representative of two independent experiments.

**Figure S6.**
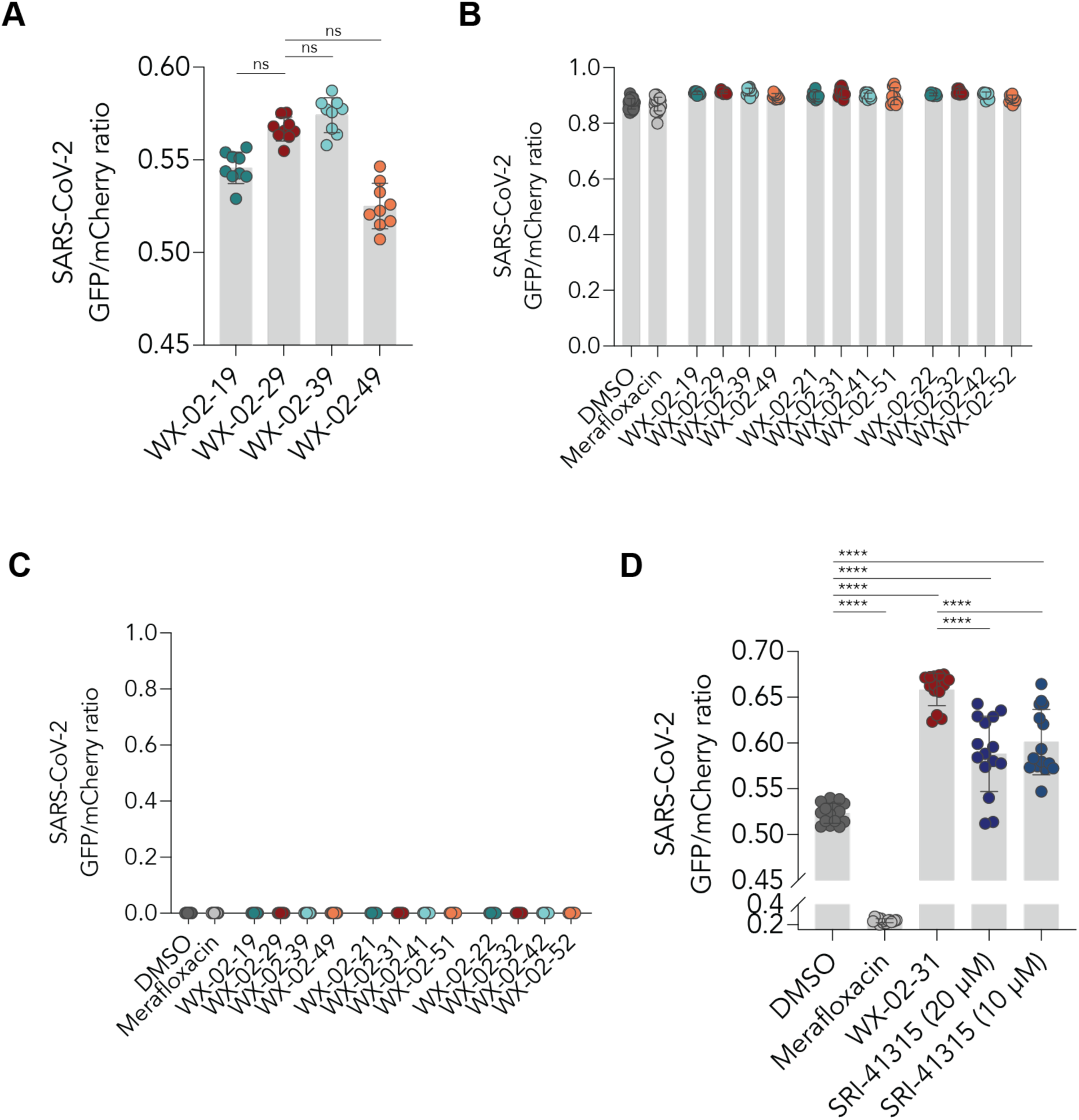
Further characterization of the effects of photo-stereoprobes on protein translation. (A-D) Effects of photo-stereoprobes (20 µM) or indicated compounds using the SARS-CoV-2 PRF reporter constructs from Fig. 4A. HeLa cells stably expressing the mCherry-FSE-GFP(−1) (A and D), mCherry-FSE-GFP(No-FSE) (B), or mCherry-FSE-GFP(0) (C) constructs were treated with stereoprobes and assessed by flow cytometry 48 h post treatment. DMSO and merafloxacin (20 µM) (34) were included as controls. Data are average values ± SD for three independent experiments.

**Figure S7.**
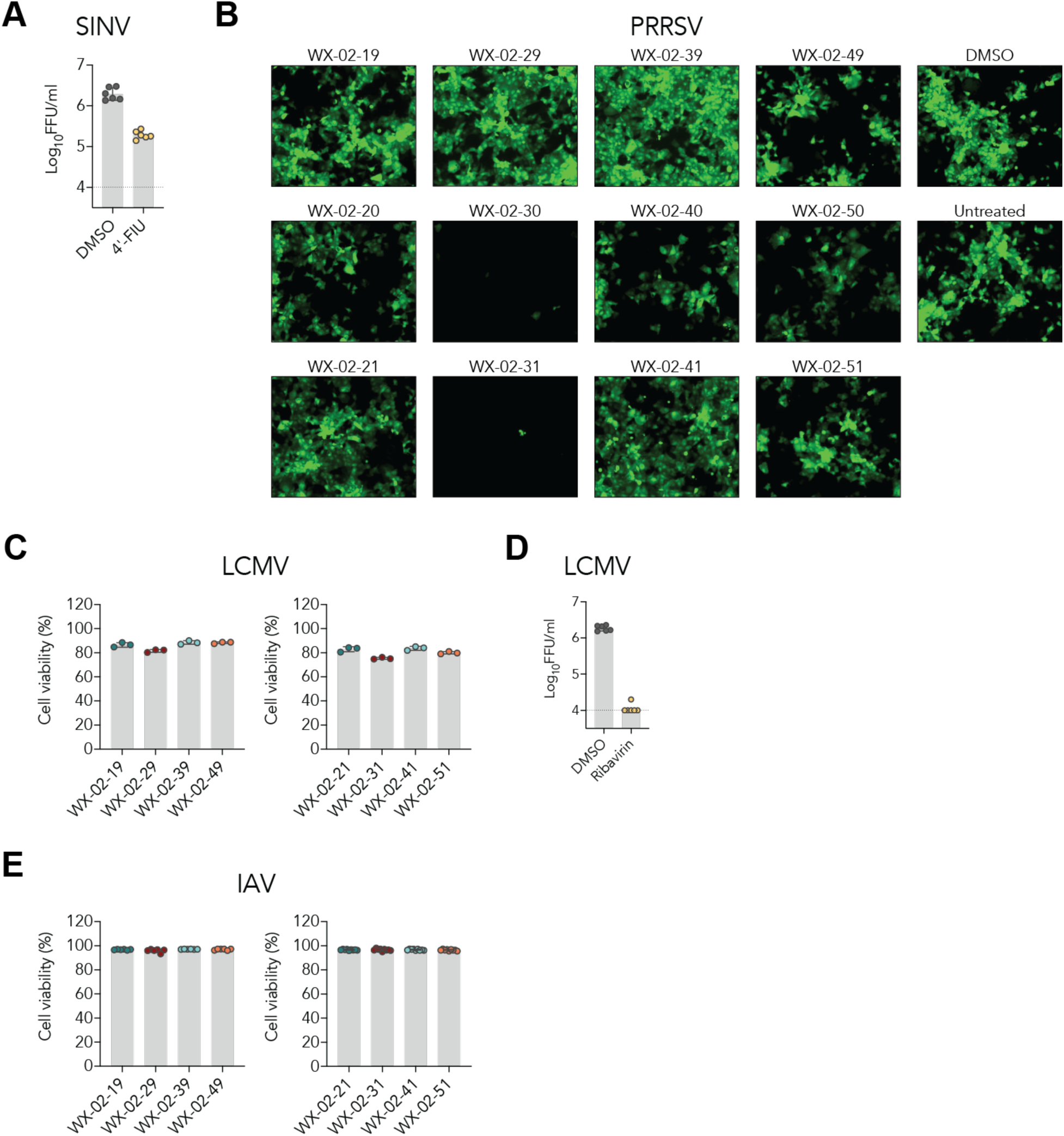
Characterization of photo-stereoprobes in additional viral infection models. (A) HeLa cells were pre-treated with either DMSO or 4’-FIU (10 µM) (56), inoculated with SINV (MOI 3) in the presence of compounds, and viral supernatant titers were assessed after 48 h. Data are average values ± SD for two independent experiments. (B) MA104 cells were pre-treated with the indicated photo-stereoprobes (10 µM), inoculated with PRRSV-GFP (MOI 0.1) in the presence of compounds, and viral antigen levels were assessed by GFP expression after 24 h. Data are from a single experiment representative of two independent experiments. (C-D) HeLa cells were pre-treated with DMSO, the indicated photo-stereoprobes (10 µM, 3 h) or ribavirin (100 µM, 3 h) (57) and inoculated with LCMV (MOI 0.01) in the presence of compounds. Cell viability (C) and viral supernatant titers (D) were assessed after 48 h. Data are average values ± SD for two independent experiments. (E) Calu-3 cells were pre-treated with the indicated photo-stereoprobes (20 µM, 3 h), inoculated with IAV (strain WSN, MOI 0.005) in the presence of compounds and cell viability was assessed by flow cytometry after 48 h. For (C and E), cell viability was assessed using the Zombie Violet Fixable Viability Kit. Data are average values ± SD for 2-3 independent experiments.

## ACKNOWLEDGEMENTS

We thank Susan Weiss (University of Pennsylvania) and Hani Zaher (Washington University in St. Louis) for invaluable discussions. We thank Xuedong Liu and Bing Chen (WuXi AppTec) for small-molecule synthesis and Brandon Orzolek, Jason Lee, Quynh Wong, Jillian Smith and Catherine Chiang (Scripps Automated Synthesis Facility) for support with high-resolution mass spectrometry). This work was supported by NIH grants U19 AI142784 (B.F.C.), T32 AI007354 (A.S.K.), R21 AI180295 (A.L.B.), and F32 CA265211 (C.J.R.). A.S.K. was supported by Open Philanthropy and the Life Sciences Research Foundation.

## AUTHOR CONTRIBUTIONS

A.S.K. and B.F.C. conceived the study. A.S.K. performed viral infection studies with help from K.M. A.S.K. performed recombinant protein expression. A.S.K. designed and performed proteomics experiments. A.S.K., D.O., and B.M. performed proteomics data analysis. A.S.K. performed target validation studies with help from K.M. and D.C.L. T.S. and A.L.B. performed PRRSV infection studies. B.M. supervised compound synthesis and characterization with C.J.R. J.C.T. contributed key reagents. A.S.K., A.L.B., J.C.T., J.R.T., and B.F.C. obtained funding. A.S.K. and B.F.C. wrote the initial draft, with the other authors providing comments.

## COMPETING FINANCIAL INTERESTS

The authors declare no competing financial interests.

## MATERIALS AND METHODS

### Cells

Calu-3 (HTB-55), Vero E6 (CRL-1586), HEK-293T (CRL-3216), HeLa (CCL-2), NIH-3T3 (CRL-1658), and MA-104 (CRL-2378.1) cells were obtained from the American Type Culture Collection (ATCC). All cell lines, except for Calu-3 cells, were maintained at 37°C in DMEM supplemented with 10% fetal bovine serum (Omega), 100 U/ml penicillin, 100 µg/ml streptomycin, and 10 mM HEPES. Calu-3 cells were maintained in EMEM supplemented with 20% fetal bovine serum (Omega), 100 U/ml penicillin, 100 µg/ml streptomycin, and 10 mM HEPES. HeLa-ACE2 cells were generated by lentiviral transduction of human ACE2 as previously described (58).

### Viruses

All virus work was approved by the Institutional Biosafety Committee at the Scripps Research Institute. SARS-CoV-2 strains USA-WA1/2020 (NR-52281) and icSARS-CoV-2-eGFP (NR-54002) were obtained from BEI and propagated in Vero-TMPRSS2 cells (gift from Sean Whelan, Washington University) and titered on Vero E6 cells (59, 60). All SARS-CoV-2 work up to December 2024 was performed in an approved biosafety level 3 facility at the Scripps Research Institute with powered air purifying respirators. Subsequent SARS-CoV-2 work was performed in an approved biosafety level 2 facility per the NIH and Institutional Biosafety Committee guidelines. HIV-1 luciferase reporter pseudovirus was generated by transfection of HEK293T cells with the pNL4-3.Luc.R-E-plasmid (41). Sindbis virus containing GFP (strain TE; gift from Matt Daugherty, UC San Diego) was propagated and titered in Vero E6 cells (61). Porcine reproductive and respiratory system virus with GFP (PRRSV-GFP) was kindly provided by Jay Calvert (Zoetis) via Bob Rowland (University of Illinois) and propagated on MARC-145 cells (62). Lymphocytic choriomeningitis virus (LCMV clone 13 variant) was propagated in BHK-21 cells and titered in Vero E6 cells (63).

### Monoclonal antibodies

The heavy and light variable chains of CR3022 (64), S309 (65), and DC2.112 (61) were codon-optimized, synthesized (Integrated DNA Technologies), and cloned into mammalian expression human IgG1 antibody vectors (GenBank FJ475055 and FJ475056) under the control of a CMV promoter. After sequence confirmation, Expi293 cells were transfected with antibody plasmids that were diluted at a 1:1 ratio in Opti-MEM and complexed with 293fectin (Thermo Fisher). Transfected cells were supplemented with Expi293 medium and 2% (w/v) Hyclone Cell Boost. Supernatant was harvested five days post transfection and purified using Protein A Sepharose 4B (Thermo Fisher) and dialyzed into PBS overnight at 4°C. Purity of antibodies was assessed by SDS-PAGE. The anti-LCMV nucleoprotein antibody VL4 (BE0106) was commercially obtained from Bio X Cell.

### Protein expression and purification

A codon-optimized DNA fragment encoding residues 1-437 of the human ETF1 protein (UniProt P62495) and a C-terminal His tag was synthesized (Integrated DNA Technologies) and cloned into the pET21a bacterial expression vector. After sequence confirmation, ETF1 was transformed into BL21(DE3) competent cells (New England Biolabs). Cells were grown in LB at 37°C to an OD_600_ of 0.8 and then induced with 1 mM isopropyl-β-D-thiogalactopyranoside overnight at 18°C. Cells were harvested and resuspended in 50 mM Tris-HCl, 150 mM NaCl pH 8.5 containing 1% Triton X-100, 100 μg/ml lysozyme, 100 μg/ml DNase I, 10 mM MgCl_2_, and 10 mM CaCl_2_, and Complete EDTA-free protease inhibitor cocktail (Roche). After stirring for 1 h, resuspended cells were sonicated and the lysate centrifuged at 6,000 x g for 30 min. The clarified supernatant was subjected to nickel affinity chromatography and the eluted protein was dialyzed overnight in 1X PBS buffer at 4°C using dialysis cassettes. Purity of the ETF1 protein was assessed by SDS-PAGE.

### SARS-CoV-2 phenotypic screen

Calu-3 cells were seeded a day prior in 96 well plates (Corning) at a density of 3 x 10^4^ cells per well. The following day, cells were treated with compounds (20 µM) for 3 h and then inoculated with SARS-CoV-2 (strain USA-WA1/2020) at an MOI of 0.1. Virus supernatant was harvested 48 hours post infection (hpi) and virus titers were determined on Vero E6 cells as previously described (59). Vero E6 cells were seeded a day prior in 96 wells at 3 x 10^4^ cells per well. Serially diluted virus was added to Vero E6 for 1 h at 37°C, and was followed by a 1% methylcellulose overlay in Minimal Essential Medium supplemented with 2% fetal bovine serum. After 24 hpi, viral supernatant was removed from Vero E6 cells and fixed with 4% paraformaldehyde for 20 mins. Plates were washed with PBS and then incubated with CR3022 antibody (1 µg/ml) diluted in a permeabilization wash buffer containing 1X PBS, 1% (w/v) saponin, and 1% (w/v) bovine serum albumin for 2 h at room temperature. Plates were washed with PBS containing 0.05% Tween-20 (PBST) and incubated with horseradish peroxidase conjugated goat anti-human IgG (H+L) (1:5,000 dilution; Jackson ImmunoResearch) for 1 h at room temperature. After washing with PBST, plates were developed with TrueBlue substrate (KPL) and foci were quantitated using a CTL ImmunoSpot plate reader. Compound hits were prioritized for secondary screening based off four criteria: 1) >1 log_10_-fold reduction of SARS-CoV-2 infectivity; 2) >1 log_10_-fold stereoselective inhibition; 3) >80% cell growth as determined by CellTiter-Glo luminescent cell viability assay.

### Inhibition assays

<u>SARS-CoV-2/Calu-3.</u> Dose-response curves of the hit compounds against SARS-CoV-2 were determined by treating Calu-3 cells with serially diluted compounds for 3 h at 37°C and then inoculated with SARS-CoV-2 (strain USA-WA1/2020) at an MOI of 0.1. Virus titers were determined on Vero E6 cells as described above in “SARS-CoV-2 phenotypic screen.” <u>SARS-CoV-2/HeLa-ACE2.</u> HeLa-ACE2 cells were treated with compounds (10 µM) for 3 h and then inoculated with icSARS-CoV-2-eGFP (MOI 0.01). Cells were harvested 24 hpi, fixed with 4% paraformaldehyde for 20 mins, and incubated with S309 antibody (1 µg/ml) diluted in Foxp3 Transcription Factor Permeabilization buffer (Thermo Fisher) for 30 mins at 4°C. Cells were washed then incubated with Alexa Fluor 647 conjugated goat anti-human IgG (1:2000 dilution; Thermo Fisher A21445) for 30 min at 4°C. After washing, cells were analyzed on a NovoCyte Quanteon flow cytometer (Agilent) and data analyzed with FlowJo 10. <u>HIV-1 pseudovirus.</u> HEK293T cells were seeded a day prior in 96 well plates at a density of 3 x 10^4^ cells/well. The following day, cells were treated with compounds (20 µM) for 3 h then transfected with pNL4-3.Luc.R-E-plasmid using Fugene HD (Promega). Cells were lysed and assayed 24 h post transfection for luciferase activity using ONE-Glo EX Luciferase Assay System per manufacturer’s instructions (Promega) on a CLARIOstar plate reader. <u>SINV and LCMV</u>. HeLa cells, which were seeded a day prior at 3 x 10^4^ cells per well, were treated with compounds (20 µM) for 3 h and inoculated with SINV-GFP or LCMV at an MOI of 1 or 0.1, respectively. Virus supernatant was harvested 48 hpi and virus titers were determined using Vero E6 cells as described above in “SARS-CoV-2 phenotypic screen” but with DC2.112 (SINV-GFP) or VL4 (LCMV) as the primary staining antibodies. <u>PRRSV-GFP</u>. MA-104 cells were seeded in 24 well plates at 2 x 10^5^ cells per well one day prior to inoculation. Cells were treated with compounds (10 µM) for 3 h, inoculated with PRRSV-GFP (MOI 0.1), and GFP levels and cell viability were assessed by flow cytometry 48 hpi.

### Cell viability assays

<u>Luminescent cell viability</u>. Calu-3, HeLa, and NIH/3T3 cells were treated with serially diluted compounds and cell viability was assayed 24 h and 48 h post treatment with CellTiter-Glo Luminescent Cell Viability Assay (Promega) per manufacturer’s instructions. Cell viability was normalized to a DMSO-treated cells. <u>Fixable viability dye</u>. Prior to paraformaldehyde fixation, cells were incubated with Zombie Violet Fixable Viability Kit (1:1,000; BioLegend) for 10 mins at room temperature. Cells were washed, fixed, and subjected to antibody staining and flow cytometry analysis as described above.

### Gel-based proteome analysis

Calu-3 (5 x 10^5^ cells/well) and HEK293T (2.5 x 10^5^ cells/well) cells were seeded in 12 well plates. The following day, cells were treated with compounds (20 µM) for 3 h and photo-crosslinked with a UV Stratalinker 1800 (Stratagene) for 10 mins at 4°C. Cells were washed with cold PBS, harvested, resuspended in cold PBS, and probe sonicated. Protein concentration was determined DC protein assay (Bio-Rad) and 50 µg of proteome was conjugated to Alexa Fluor 647 (AF647) azide through copper-catalyzed azide-alkyne cycloaddition (CuAAC). A reagent mixture containing 0.1 mM tris(benzyltriazolylmethyl)amine (TBTA) (3 µl/sample, 1.7 mM in 1:4 DMSO:t-ButOH), 1 mM CuSO_4_ (1 µl/sample, 50 mM in H_2_O), 12.5 mM Alexa Fluor 647 (AF647) azide (1 µl/sample, 0.625 mM in DMSO), and freshly prepared 1 mM tris(2-carboxyethyl)phosphine HCl (TCEP) (1 µl/sample, 50 mM in H_2_O) was added to conjugate AF647-azide to the probe-crosslinked proteome. CuAAC reactions were quenched after 1 h at room temperature with the addition of 4X SDS loading buffer. The AF647-labeled proteome was separated (15 µg/well) on a 4-12% Bis-Tris protein gel (Thermo Fisher) and visualized by in-gel fluorescence with a ChemiDoc MP (Bio-Rad). Images were processed and analyzed using Image Lab software (Bio-Rad). To determine probe labeling of recombinantly overexpressed ETF1, HEK293T cells were seeded in 12 well plates and transiently transfected with a pCDNA3.1 mammalian expression vector encoding full-length human ETF1 (UniProt P62495) and a C-terminal FLAG tag. Transfected cells were treated with compounds (20 µM) for 3 h and then UV-crosslinked at 4°C. Cells were processed and fluorescent gels were analyzed as previously described in this section.

### Western blots

Expression of ETF1 in Calu-3 cells and overexpression of recombinant ETF1 in HEK293T was confirmed by western blot analysis. After separation of the proteome by SDS-PAGE, the protein was transferred to a PVDF membrane using a Trans-Blot Turbo Transfer System (Bio-Rad). The membrane was blocked with 4% (w/v) nonfat milk in 1X PBS and 0.05% Tween-20 for 1 h at room temperature and then incubated overnight at 4°C with anti-ETF1 antibody (Abcam ab31799), anti-FLAG antibody (Sigma F3165), or anti-GAPDH (Thermo Fisher AM4300) antibody diluted (1:1,000 dilution) in the blocking buffer. After washing, the membrane was incubated with horseradish peroxidase conjugated goat anti-rabbit or goat anti-mouse IgG (1:10,000 dilution, Jackson ImmunoResearch) for 1 h at room temperature, washed, developed with Pierce ECL Western Blotting Substrate (Thermo Fisher), and scanned with a ChemiDoc MP Imaging System (Bio-Rad).

### Mass spectrometry-based proteome analysis

Calu-3 cells were grown to 80% confluency in 10 cm dishes. Cells were treated with photo-stereoprobes diluted in supplemented EMEM media for 3 h and then UV crosslinked at 365 nm (Agilent Stratalinker 1800) for 10 min. Cells were washed 3x with cold DPBS and detached using a cell scraper. Cell pellets were snap frozen in liquid nitrogen and kept at −80°C until use. For MS-based experiments with SARS-CoV-2 infected cells, Calu-3 cells were inoculated with SARS-CoV-2 (MOI 0.1) and 24 hpi, cells were treated with photo-stereoprobes and harvested as described above with the appropriate biosafety protocols. Photo-stereoprobe treated cell pellets were resuspended in DPBS and probe-sonicated (Branson). For SARS-CoV-2 infected cells, cell pellets were resuspended in DPBS with 1% Triton X-100 to inactivate virus prior to probe sonication. Cell lysates were normalized to 2 mg/ml (500 µl) and conjugated to biotin-PEG_4_-azide by adding a copper-catalyzed alkyne-azide cycloaddition (CuAAC) master mix (5 µl of 10 mM biotin-PEG_4_-azide in DMSO, 10 µl of freshly prepared 50 mM TCEP in H_2_O, 30 µl of 1.7 mM TBTA in t-BuOH:DMSO (4:1, v/v), and 10 µl of 50 mM CuSO_4_ in H_2_O) for 1 h at room temperature. Biotin-conjugated cell lysates were precipitated out of solution through the addition of cold methanol (600 µl) and chloroform (200 µl) followed by vortexing and centrifugation at 16,000 x *g* for 10 mins to create a protein disk. The protein disks were resuspended in cold methanol, sonicated, and centrifuged at 16,000 x *g* for 10 mins to generate a pellet which was stored at −80°C until use. For SARS-CoV-2 infected cells, protein pellets were subjected to an additional wash with cold methanol. The protein pellets were resuspended in 8 M urea in DPBS (500 µl) and 10 µl of 10% SDS followed by probe sonication. Sonicated samples were reduced by the addition of 10 mM dithiothreitol and incubation at 65°C for 15 mins, alkylated with the addition of 20 mM iodoacetamide and incubation at 37°C for 30 mins, and quenched with the addition of 130 µl of 10% SDS. Samples were then added to streptavidin beads and incubated rotating for 1.5 h at room temperature. After incubation, streptavidin beads were washed with 0.2% SDS in DPBS (2 x 10 ml washes), DPBS (1 x 10 ml wash), and then transferred to Protein LoBind tubes (Eppendorf). Streptavidin beads were additionally washed with HPLC-grade water (2 x 1 ml washes) and 200 mM EPPS (1 x 1 ml wash). Streptavidin-enriched proteins were digested on beads shaking overnight at 37°C in 200 µl of buffer containing 2M urea, 200 mM EPPS, pH 8.0, 1 mM CaCl2 and 10 µg/ml sequencing-grade trypsin (Promega). After trypsin digestion, the supernatant containing digested peptides were transferred to a new Protein LoBind tube and acetonitrile was added to a final volume of 30% (v/v). Digested peptides were tandem mass tag (TMT)-labeled with the addition of 6 µl (20 µg/µl stock concentration in dry acetonitrile) of TMT 10-plex or TMT 16-plex (Thermo Fisher), incubated for 1.5 h at room temperature, and quenched with the addition of 6 µl of a 5% hydroxylamine solution for 15 mins followed by 20 µl of 100% formic acid. Samples were combined, dried overnight in a SpeedVac, and then desalted using a Sep-Pak C18 cartridge (Waters).

### HPLC fractionation

Desalted samples were resuspended in 500 µl of Buffer A (95% HPLC grade water, 5% acetonitrile, 0.1% formic acid), sonicated, and subjected to HPLC fractionation (Agilent) into 96 deep-well plate containing 20 µl of 20% formic acid using a ZORBAX 300Extend-C18, 3.5 µm column (Agilent). Samples were separated at a flow rate of 0.5 ml/min using the following gradient: 100% Buffer A from 0-2 min, 0%-13% Buffer B from 2-3 min, 13%-42% Buffer B from 3-60 min, 42%-100% Buffer B from 60-61 min, 100% Buffer B from 61-65 min, 100%-0% Buffer B from 65-66 min, 100% Buffer A from 66-75 min, 0%-13% Buffer B from 75-78 min, 13%-80% Buffer B from 78-80 min, 80% Buffer B from 80-85 min, 100% Buffer A from 86-91 min, 0%-13% Buffer B from 91-94 min, 13%-80% Buffer B from 94-96 min, 80% Buffer B from 96-101 min, and 80%-0% Buffer B from 101-102 min (Buffer A: 10 mM aqueous NH_4_HCO_3_; Buffer B: Acetonitrile). Fractionated samples were evaporated to dryness with a SpeedVac and peptides were resuspended in 80% acetonitrile, 20% HPLC-grade water, 0.1% formic acid (3 x 300 µl/well) and combined to a total of 12 fractions (e.g., Fraction 1: wells 1A + 1B…+ 1H; Fraction 2: wells 2A + 2B…+ 2H; etc.). Samples were dried using a SpeedVac, resuspended in Buffer A (95% HPLC grade water, 5% acetonitrile, 0.1% formic acid), and subjected to mass spectrometry analysis.

### Tandem mass tag liquid chromatography mass spectrometry

Samples were analyzed by liquid chromatography tandem mass-spectrometry using an Orbitrap Fusion Mass Spectrometer (Thermo Fisher) coupled to an UltiMate 3000 Series Rapid Separation LC system (Thermo Fisher) and Dionex Autosampler (Thermo Fisher), as previously described (24). The peptides were eluted onto a capillary column (75 μm inner diameter fused silica, packed with C18 (Waters, Acquity BEH C18, 1.7 μm, 25 cm) or an EASY-Spray HPLC column (Thermo Fisher, ES902, ES903) using an Acclaim PepMap 100 (Thermo Fisher, 164535) loading column, and separated at a flow rate of 0.25 μl/min. Data was acquired using either an MS3-based TMT method on Orbitrap Fusion or Orbitrap Eclipse Tribrid Mass Spectrometer. Briefly, the scan sequence began with an MS1 master scan (Orbitrap analysis, resolution 120,000, 400−1700 m/z, RF lens 60%, automatic gain control [AGC] target 2E5, maximum injection time 50 ms, centroid mode) with dynamic exclusion enabled (repeat count 1, duration 15 s). The top ten precursors were then selected for MS2/MS3 analysis. MS2 analysis consisted of: quadrupole isolation (isolation window 0.7) of precursor ion followed by collision-induced dissociation (CID) in the ion trap (AGC 1.8E4, normalized collision energy 35%, maximum injection time 120 ms). Following the acquisition of each MS2 spectrum, synchronous precursor selection (SPS) enabled the selection of up to 10 MS2 fragment ions for MS3 analysis. MS3 precursors were fragmented by HCD and analyzed using the Orbitrap (collision energy 55%, AGC 1.5e5, maximum injection time 120 ms, resolution 50,000). For MS3 analysis, we used charge state-dependent isolation windows. For charge state z = 2, the MS isolation window was set at 1.2; for z = 3-6, the MS isolation window was set at 0.7. Raw files were uploaded to Integrated Proteomics Pipeline (IP2) available at (http://ip2.scripps.edu/ip2/mainMenu.html) and MS2 and MS3 files extracted from the raw files using RAW Converter and searched using the ProLuCID algorithm using a reverse concatenated, non-redundant variant of the human (released July 2016) and the Severe acute respiratory syndrome coronavirus 2 (2019-nCoV) (released March 2020) UniProt databases. Cysteine residues were searched with a static modification for carboxyamidomethylation (+57.02146 Da). N-termini and lysine residues were also searched with a static modification corresponding to the TMT tag (+229.1629 Da for 10-plex and +304.2071 Da for 16-plex). Peptides were required to be at least 6 amino acids long. ProLuCID data was filtered through DTASelect (version 2.0) to achieve a peptide false-positive rate below 1%. The MS3-based peptide quantification was performed with reporter ion mass tolerance set to 20 ppm with Integrated Proteomics Pipeline (IP2).

### Proteomics data analysis

Enrichment ratios (probe versus probe) were calculated for each peptide-spectra match by dividing each TMT reporter ion intensity by the sum intensity of all TMT reporter ion channels as previously described (24). Peptide-spectra matches with summed reporter ion intensities ≥ 10,000, coefficient of variation of ≥ 0.5, and ≥ 2 distinct peptides were grouped based on protein ID. Replicate channels were grouped across each experiment, and average values were calculated for each protein. A variability metric was also computed across replicate channels, which equaled the ratio of median absolute deviation to average and was expressed in percentage. A protein was considered stereoselectively liganded if the average enrichment (of at least 2 biological replicates) by a photo-stereoprobe was > 2-fold that of its enantiomer and the variability corresponding to the photo-stereoprobe leading to the highest enrichment did not exceed 50%. The following additional filtering criteria were applied for removing low-quality enrichment events: 1) enrichment events of an enantiomer pair showing signals > 3-fold lower than the larger signal of their diastereomers were omitted; 2) enrichment events of an enantiomer pair with their lower signal < 1 were omitted. 3) enrichment events with diastereo-ratio (min) :: 2 were omitted unless the liganded protein was enantioselectively engaged by both enantiomer pairs of the photo-stereoprobes. “Diastereo-ratio (min)” refers to the highest signal (across four stereoisomers) divided by the lower signal of its diastereomers.

### Frameshift reporter assay

Photo-stereoprobes were tested for programmed ribosomal frameshifting using stable HeLa frameshifting reporter cell lines. The SARS-CoV-2 frameshifting reporter lentiviral plasmids mCherry-SARS2-FSE-GFP(−1) (Addgene #177168), mCherry-SARS2-FSE-GFP(No-FSE) (Addgene #177619), and mCherry-SARS2-FSE-GFP(0) (Addgene #177620) were obtained from Addgene (34). The HIV-1 frameshifting reporter lentiviral plasmid mCherry-HIV-FSE-GFP(−1) was generated by replacing the SARS-CoV-2 frameshift sequence with the HIV-1 frameshift sequence in mCherry-SARS2-FSE-GFP(−1) (Addgene #177168). The HIV-1 frameshift sequence was adapted from pAGH10 (Addgene #198224). Lentivirus was packaged by transfection of HEK293T cells with the frameshifting reporter lentiviral plasmid,psPAX2 (Addgene #12260), and pMD2.G (Addgene #12259) and harvested 48 h post transfection. HeLa cells were transduced with the frameshifting reporter lentiviruses for two days and then sorted for cells that were both mCherry and GFP positive. Sorted cells were expanded in the presence of puromycin and transduction of the lentiviral plasmids were confirmed by flow cytometry. Stable HeLa frameshifting reporter cell lines were treated photo-stereoprobes (20 µM), DMSO, or merafloxacin (20 µM) for 48 h. Cells were trypsinized, resuspended in 1X PBS supplemented with 2% FBS and 2 mM EDTA, analyzed on a NovoCyte Quanteon flow cytometer (Agilent), and data analyzed with FlowJo 10. The frameshifting ratio was determined by the ratio of GFP-positive cells to mCherry-positive cells.

### Translation readthrough assay

HEK293T (2.5 x 10^5^ cells/well) cells were seeded in 12 well plates and the following day, cells were transfected with the GFP reporter plasmid pMHG-W57X (37) using Fugene HD. Transfected cells were treated with either DMSO or compounds for 24 h and then GFP levels were assessed NovoCyte Quanteon flow cytometer (Agilent). GFP-positive cells were analyzed by gating on untransfected HEK293T cells using FlowJo 10.

## Notes

### Competing Interest Statement

The authors have declared no competing interest.

