## Supplementary material for "Integrated phenotypic screening and chemical proteomics identifies ETF1 ligands that modulate viral translation and replication": Synthetic Chemistry Procedures and Analytical Data

**General considerations**

All chemical reagents were purchased from commercial suppliers and used without further purification unless otherwise noted.

Preparative high-pressure liquid chromatography (prep-HPLC) was performed on a Gilson GX-281 instrument eluting with a mixture of acetonitrile and a buffered aqueous phase.

NMR spectra were recorded on Bruker Avance III 400, Avance III HD 400, or Avance Neo 400 spectrometers. Spectra are reported as follows: chemical shift (δ, ppm relative to residual solvent), apparent multiplicity (s = singlet, br. s = broad singlet, d = doublet, t = triplet, q = quartet, h = hextet, m = multiplet, or a combination thereof), and coupling constant (*J*, Hz).

Mass measurements for high-resolution mass spectrometry (HRMS) were performed on a Waters Xevo G2-XS TOF calibrated against sodium formate clusters and using a LeuEnk lockmass. Expected monoisotopic masses were calculated using MassLynx 4.1 and the m/z values for calibrant and lockmass were MassLynx-default values.

Analytical supercritical fluid chromatography (SFC) was performed on a Shimadzu LC system (flow rate: 3 mL/min, back pressure: 100 Bar, column temperature: 35 ºC, phase A: supercritical $CO_2$, phase B as indicated) equipped with a polydiode array detector.

**Synthetic procedure**

The compounds used in this study were synthesized by following the general procedure below starting from **S1**, which was previously reported or synthesized in an analogous fashion (1-3). Representative protocols and purifications are provided for each compound.

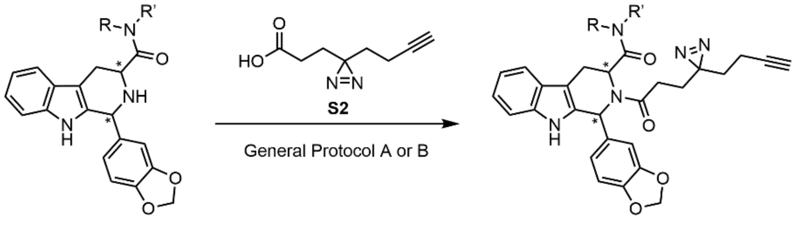


**S1** **S3**

**General Protocol A:** To a solution of **S1** (1 equiv) in dimethylformamide (DMF, 0.1 M) were added *O*-(7-azabenzotriazol-1-yl)-*N*,*N*,*N'*,*N'*-tetramethyluronium hexafluorophosphate (HATU, 1.2 equiv), **S2** (1 equiv), and *N*,*N*-diisopropylethylamine (DIPEA, 1.5 equiv). The mixture was stirred at 25 °C for 2–16 hours. Upon completion, the reaction mixture was diluted with water and extracted with ethyl acetate (3×). The combined organic layers were dried over anhydrous sodium sulfate, filtered and concentrated under reduced pressure to give a residue, which was purified by flash column chromatography ($SiO_2$), preparative thin layer chromatography (prep-TLC), and/or prep-HPLC to afford **S3**.

**General Protocol B:** To a solution of **S1** (1 equiv) in methylene chloride ($CH_2Cl_2$, 0.1 M) were added triethylamine (TEA, 3 equiv), bis(2-oxo-3-oxazolidinyl)phosphinic chloride (BOP-Cl,1.2 equiv), and **S2** (1 equiv). The mixture was stirred at 25 °C for 2–16 hours. Upon completion, the reaction mixture was diluted with water and extracted with ethyl acetate (3×). The combined organic layers were dried over anhydrous sodium sulfate, filtered and concentrated under reduced pressure to give a residue, which was purified by flash column chromatography ($SiO_2$), prep-TLC, and/or prep-HPLC to afford **S3**.

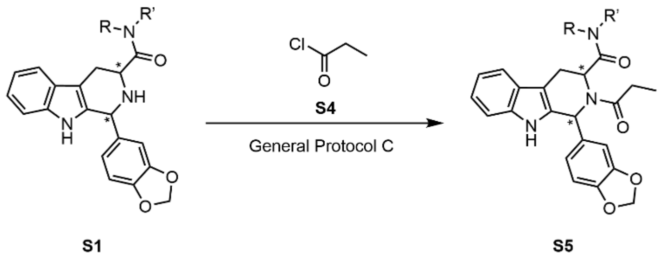

**General Protocol C:** To a solution of **S1** (1 equiv) and DIPEA (3 equiv) in $CH_2Cl_2$ (0.1 M) at 0 ºC was added **S4** (1.2 equiv). The mixture was stirred at the same temperature for 0.5 hours. Upon completion, the reaction mixture was filtered and concentrated under reduced pressure to give a residue, which was purified by flash column chromatography ($SiO_2$), preparative thin layer chromatography (prep-TLC), and/or prep-HPLC to afford **S5**.

### Analytical characterization

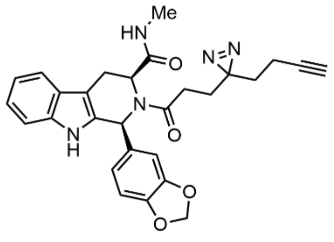

(1*S*,3*S*)-1-(benzo[*d*][1,3]dioxol-5-yl)-2-(3-(3-(but-3-yn-1-yl)-3*H*-diazirin-3-yl)propanoyl)-*N*-methyl-2,3,4,9-tetrahydro-1*H*-pyrido[3,4-*b*]indole-3-carboxamide (**WX-02-221**) – see ref (1).

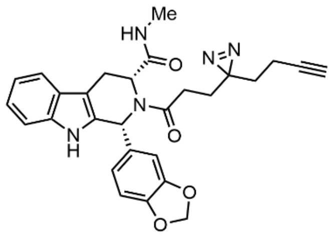

(1*R*,3*R*)-1-(benzo[*d*][1,3]dioxol-5-yl)-2-(3-(3-(but-3-yn-1-yl)-3*H*-diazirin-3-yl)propanoyl)-*N*-methyl-2,3,4,9-tetrahydro-1*H*-pyrido[3,4-*b*]indole-3-carboxamide (**WX-02-222**)

General Protocol A with prep-HPLC purification on a Phenomenex Unisil 3-100 C18 Ultra column (150 mm × 50 mm × 3 µm; mobile phase: [A: water containing 0.225% formic acid–MeCN]; B%: 45–65%, 10 min) and then further purification on a Phenomenex Luna C18 column (150 mm × 25 mm × 10 µm; mobile phase: [A: water containing 0.225% formic acid–MeCN]; B%: 52–82%, 10 min) to obtain WX-02-222 (22.3 mg, 31% yield) as an off-white solid.

**$^1$H NMR** (400 MHz, $CD_3OD$, 297 K) δ 7.53 (d, $J$ = 7.8 Hz, 1H), 7.27 (d, $J$ = 8.0 Hz, 1H), 7.09 (t, $J$ = 7.4 Hz, 1H), 7.06 – 6.91 (m, 2H), 6.81 (br. s, 1H), 6.68 (br. s, 2H), 5.89 (s, 2H), 5.01 (br. s, 0.5H), 4.60 (br. s, 0.5H), 3.81 – 3.39 (m, 1H), 2.98 (dd, $J$ = 16.0, 6.7 Hz, 1H), 2.63 – 2.32 (m, 3H), 2.33 – 2.15 (m, 3H), 2.10 – 1.95 (m, 2H), 1.94 – 1.76 (m, 2H), 1.74 – 1.52 (m, 2H). 2 exchangeable protons not observed.

**HRMS** ESI-TOF m/z calculated for $C_{28}H_{28}N_5O_4$ $[M+H]^+$ 498.2141. Found 498.2151.

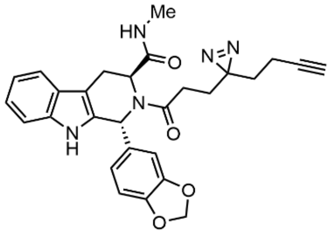

(1*R*,3*S*)-1-(benzo[*d*][1,3]dioxol-5-yl)-2-(3-(3-(but-3-yn-1-yl)-3*H*-diazirin-3-yl)propanoyl)-*N*-methyl-2,3,4,9-tetrahydro-1*H*-pyrido[3,4-*b*]indole-3-carboxamide (**WX-02-223**)

General Protocol A with prep-HPLC purification on a Phenomenex Synergi C18 column (150 mm × 25 mm × 10 µm; mobile phase: [A: water containing 0.225% formic acid–MeCN]; B%: 41–71%, 10 min) to obtain WX-02-223 (19.3 mg, 16% yield) as a yellow solid.

**$^1$H NMR** (400 MHz, $CD_3OD$, 323 K): δ 7.40 (d, *J* = 7.8 Hz, 1H), 7.25 (d, *J* = 8.0 Hz, 1H), 7.04 (t, *J* = 7.5 Hz, 1H), 6.97 (t, *J* = 7.3 Hz, 1H), 6.94 – 6.85 (m, 2H), 6.85 – 6.65 (m, 1H), 6.29 – 6.08 (m, 1H), 5.98 – 5.79 (m, 2H), 5.36 – 5.00 (m, 1H), 3.53 – 3.36 (m, 1H), 2.60 (s, 3H), 2.36 (ddd, *J* = 15.5, 8.8, 6.2 Hz, 1H). 2.27 – 2.07 (m, 2H), 1.98 – 1.86 (m, 2H), 1.83 – 1.40 (m, 4H). 2 exchangeable protons not observed, and 1 proton overlaps residual water.

**HRMS** ESI-TOF m/z calculated for $C_{28}H_{28}N_5O_4$ $[M+H]^+$ 498.2141. Found 498.2153.

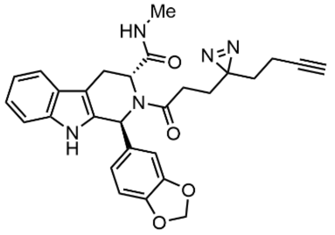

(1*S*,3*R*)-1-(benzo[*d*][1,3]dioxol-5-yl)-2-(3-(3-(but-3-yn-1-yl)-3*H*-diazirin-3-yl)propanoyl)-*N*-methyl-2,3,4,9-tetrahydro-1*H*-pyrido[3,4-*b*]indole-3-carboxamide (**WX-02-224**)

General Protocol A with prep-HPLC purification on a Phenomenex Unisil 3-100 C18 Ultra column (150 mm × 50 mm × 3 µm; mobile phase: [A: water containing 0.225% formic acid–MeCN]; B%: 45–65%, 10 min) and then further purification on a Phenomenex Luna C18 column (150 mm × 25 mm × 10 µm; mobile phase: [A: water containing 0.225% formic acid–MeCN]; B%: 52–82%, 10 min) to obtain WX-02-224 (27.0 mg, 38% yield) as an off-white solid.

**$^1$H NMR** (400 MHz, $CD_3OD$, 323 K): δ 7.40 (d, *J* = 8.0 Hz, 1H), 7.25 (d, *J* = 8.0 Hz, 1H), 7.04 (ddd, *J* = 8.2, 7.1, 1.3 Hz, 1H), 6.97 (ddd, *J* = 8.1, 7.0, 1.1 Hz, 1H), 6.94 – 6.85 (m, 2H), 6.85 – 6.65 (m, 1H), 6.29 – 6.08 (m, 1H), 5.98 – 5.73 (m, 2H), 5.36 – 5.00 (m, 1H), 4.40 – 4.08 (m, 1H), 3.53 – 3.36 (m, 1H), 2.60 (s, 3H), 2.36 (ddd, *J* = 16.3, 8.8, 6.2 Hz, 1H), 2.19 (t, *J* = 2.7 Hz, 1H), 2.12 (ddd, *J* = 15.9, 8.7, 6.5 Hz, 1H), 1.98 – 1.86 (m, 2H), 1.78 – 1.36 (m, 4H). 2 exchangeable protons not observed.

**HRMS** ESI-TOF m/z calculated for $C_{28}H_{28}N_5O_4$ $[M+H]^+$ 498.2141. Found 498.2150.

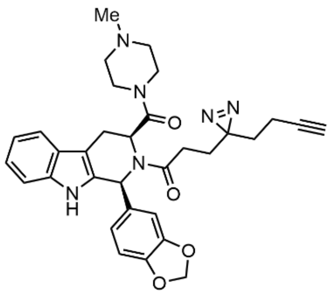

1-((1*S*,3*S*)-1-(benzo[*d*][1,3]dioxol-5-yl)-3-(4-methylpiperazine-1-carbonyl)-1,3,4,9-tetrahydro-2*H*-pyrido[3,4-*b*]indol-2-yl)-3-(3-(but-3-yn-1-yl)-3*H*-diazirin-3-yl)propan-1-one (**WX-02-19**) – see ref (1).

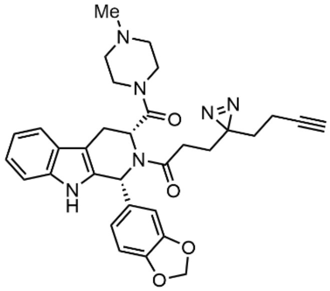

1-((1*R*,3*R*)-1-(benzo[*d*][1,3]dioxol-5-yl)-3-(4-methylpiperazine-1-carbonyl)-1,3,4,9-tetrahydro-2*H*-pyrido[3,4-*b*]indol-2-yl)-3-(3-(but-3-yn-1-yl)-3*H*-diazirin-3-yl)propan-1-one (**WX-02-39**, formic acid salt)

General Protocol A with prep-HPLC purification on a Phenomenex Gemini-NX C18 column (75 mm × 30 mm × 3 µm; mobile phase: [A: water containing 0.05% ammonium hydroxide–MeCN]; B%: 31–61%, 11.5 min) and then further purification on the same column with mobile phase: [A: water containing 0.225% formic acid–MeCN]; B%: 18–48%, 7 min) to obtain WX-02-39 (4.6 mg, 6% yield) as a white solid.

**$^1$H NMR** (400 MHz, $CD_3OD$, 298 K): δ 8.38 (br. s, 0.3H), 7.51 (d, *J* = 7.8 Hz, 1H), 7.29 (d, *J* = 8.1 Hz, 1H), 7.10 (t, *J* = 7.6 Hz, 1H), 7.03 (t, *J* = 7.4 Hz, 1H), 6.87 – 6.70 (m, 3H), 6.30 (br. s, 1H), 5.95 (d, *J* = 8.6 Hz, 2H), 5.83 (br s, 1H), 4.65 – 4.48 (m, 1H, overlaps residual water), 3.66 – 3.40 (m, 2H), 2.95 (dd, *J* = 15.5, 6.2 Hz, 1H), 2.76 – 2.17 (m, 11H), 2.11 – 1.78 (m, 4H), 1.73 – 1.54 (m, 2H). 1 exchangeable proton not observed.

**HRMS** ESI-TOF m/z calculated for $C_{32}H_{35}N_6O_4$ $[M+H]^+$ 567.2720. Found 567.2742.

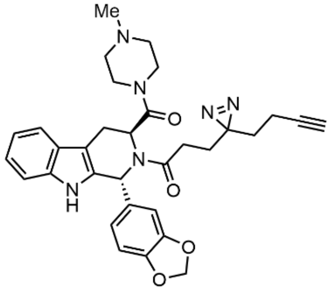

1-((1*R*,3*S*)-1-(benzo[*d*][1,3]dioxol-5-yl)-3-(4-methylpiperazine-1-carbonyl)-1,3,4,9-tetrahydro-2*H*-pyrido[3,4-*b*]indol-2-yl)-3-(3-(but-3-yn-1-yl)-3*H*-diazirin-3-yl)propan-1-one (**WX-02-29**)

General Protocol A except with 3 equiv. of DIPEA. prep-HPLC purification on a Phenomenex Synergi C18 column (150 mm × 25 mm × 10 µm; mobile phase: [A: water containing 0.225% formic acid–MeCN]; B%:

27–57%, 10 min) and then further purification on a Waters Xbridge BEH C18 column (150 mm × 25 mm × 5 µm; mobile phase: [A: water containing 10 mM ammonium bicarbonate–MeCN]; B%: 30–63%, 9 min) to obtain WX-02-29 (7.4 mg, 8% yield) as a yellow solid.

**$^1$H NMR** (400 MHz, $CD_3OD$, 323 K): δ 7.44 (d, *J* = 7.7 Hz, 1H), 7.26 (d, *J* = 8.0 Hz, 1H), 7.06 (ddd, *J* = 8.2, 7.0, 1.3 Hz, 1H), 6.99 (td, *J* = 7.5, 1.2 Hz, 1H), 6.93 – 6.85 (m, 2H), 6.84 – 6.75 (m, 1H), 6.39 – 6.10 (m, 1H), 5.91 (d, *J* = 1.7 Hz, 2H), 5.19 (t, *J* = 5.5 Hz, 1H), 3.61 – 3.33 (m, 4H), 3.29 – 3.16 (m, 2H), 2.49 – 2.20 (m, 9H), 2.19 (t, *J* = 2.6 Hz, 1H), 1.95 (td, *J* = 7.5, 2.7 Hz, 2H), 1.78 – 1.59 (m, 2H), 1.52 (t, *J* = 7.0 Hz, 2H). 1 exchangeable proton not observed.

**HRMS** ESI-TOF m/z calculated for $C_{32}H_{35}N_6O_4$ $[M+H]^+$ 567.2720. Found 567.2745.

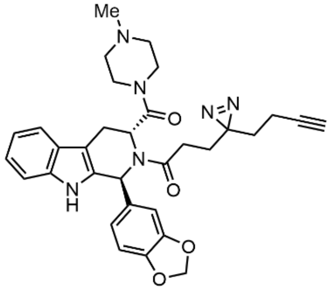


1-((1*S*,3*R*)-1-(benzo[*d*][1,3]dioxol-5-yl)-3-(4-methylpiperazine-1-carbonyl)-1,3,4,9-tetrahydro-2*H*-pyrido[3,4-*b*]indol-2-yl)-3-(3-(but-3-yn-1-yl)-3*H*-diazirin-3-yl)propan-1-one (**WX-02-49**)

General protocol A with prep-HPLC purification on a Phenomenex Gemini-NX C18 column (75 mm × 30 mm × 3 µm; mobile phase: [A: water containing 0.05% ammonium hydroxide– MeCN]; B%: 28–58%, 11.5 min) and then further purification on a Waters Xbridge BEH C18 column (150 mm × 25 mm × 5 µm; mobile phase: [A: water containing 0.05% ammonium hydoxide–MeCN]; B%: 42–47%, 10 min) to obtain WX-02-49 (8.3 mg, 12% yield) as an off-white solid.

**$^1$H NMR** (400 MHz, $CD_3OD$, 323 K): δ 7.44 (d, *J* = 7.8 Hz, 1H), 7.27 (d, *J* = 8.0 Hz, 1H), 7.06 (t, *J* = 7.5 Hz, 1H), 7.00 (t, *J* = 7.4 Hz, 1H), 6.93 – 6.85 (m, 2H), 6.85 – 6.76 (m, 1H), 6.39 – 6.10 (m, 1H), 5.93 (s, 2H), 5.21 (t, *J* = 5.5 Hz, 1H), 4.29 (m, 1H, overlaps residual water), 3.62 – 3.36 (m, 4H), 3.29 – 3.23 (m, 2H), 2.65 – 2.46 (m, 4H), 2.46 – 2.32 (m, 4H), 2.32 – 2.15 (m, 2H), 1.95 (t, *J* = 6.5 Hz, 2H), 1.78 – 1.59 (m, 2H), 1.52 (t, *J* = 7.5 Hz, 2H).

**HRMS** ESI-TOF m/z calculated for $C_{32}H_{35}N_6O_4$ $[M+H]^+$ 567.2720. Found 567.2728.

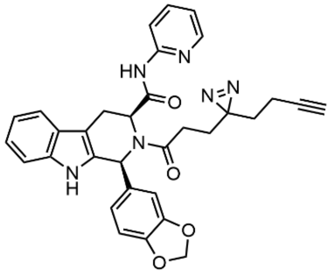


(1*S*,3*S*)-1-(benzo[*d*][1,3]dioxol-5-yl)-2-(3-(3-(but-3-yn-1-yl)-3*H*-diazirin-3-yl)propanoyl)-*N*-(pyridin-2-yl)-2,3,4,9-tetrahydro-1*H*-pyrido[3,4-*b*]indole-3-carboxamide (**WX-02-20**) – see ref (1).

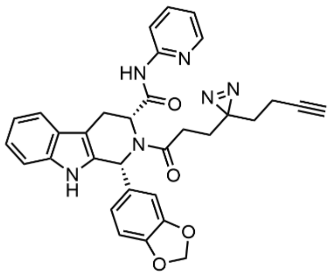

(1*R*,3*R*)-1-(benzo[*d*][1,3]dioxol-5-yl)-2-(3-(3-(but-3-yn-1-yl)-3*H*-diazirin-3-yl)propanoyl)-*N*-(pyridin-2-yl)-2,3,4,9-tetrahydro-1*H*-pyrido[3,4-*b*]indole-3-carboxamide (**WX-02-40**, formic acid salt)

General Protocol B with prep-HPLC purification on a Phenomenex Luna C18 column (150 mm × 25 mm × 10 µm; mobile phase: [A: water containing 0.225% formic acid – MeCN]; B%: 45–75%, 11 min) to obtain WX-02-40 (23.8 mg, 22% yield) as an off-white solid.

**$^1$H NMR** (400 MHz, $CD_3OD$, 323 K): δ 8.10 (d, *J* = 3.9 Hz, 1H), 7.80 (d, *J* = 8.5 Hz, 1H), 7.62 (td, *J* = 7.7, 1.4 Hz, 1H), 7.56 (d, *J* = 7.7 Hz, 1H), 7.29 (dd, *J* = 8.0, 1.2 Hz, 1H), 7.12 (td, *J* = 7.6, 1.4 Hz, 1H), 7.06 (td, *J* = 7.3, 1.1 Hz, 1H), 7.00 (dd, *J* = 7.4, 5.1 Hz, 1H), 6.76 – 6.63 (m, 2H), 6.52 – 6.22 (m, 1H), 5.64 (s, 1H), 5.61 – 5.51 (m, 1H), 5.50 – 5.16 (m, 1H), 4.29 (s, 1H), 3.81 – 3.56 (m, 1H), 3.07 (dd, *J* = 15.9, 6.8 Hz, 1H), 2.75 – 2.63 (m, 1H), 2.63 – 2.46 (m, 1H), 2.20 (t, *J* = 2.7 Hz, 1H), 2.05 (t, *J* = 7.5 Hz, 2H), 1.94 (t, *J* = 7.6 Hz, 2H), 1.67 (t, *J* = 7.4 Hz, 2H). 2 exchangeable protons not observed.

**HRMS** ESI-TOF m/z calculated for $C_{32}H_{29}N_6O_4$ $[M+H]^+$ 561.2250. Found 561.2264.

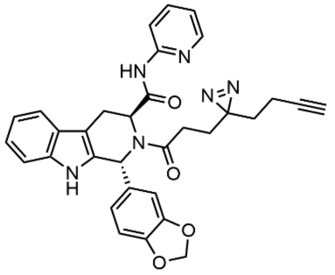

(1*R*,3*S*)-1-(benzo[*d*][1,3]dioxol-5-yl)-2-(3-(3-(but-3-yn-1-yl)-3*H*-diazirin-3-yl)propanoyl)-*N*-(pyridin-2-yl)-2,3,4,9-tetrahydro-1*H*-pyrido[3,4-*b*]indole-3-carboxamide (**WX-02-30**)

General Protocol A with 1.5 equiv HATU and 3 equiv DIPEA. prep-HPLC purification on a Phenomenex Synergi C18 column (150 mm × 25 mm × 10 µm; mobile phase: [A: water containing 0.225% formic acid – MeCN]; B%: 42–72%, 10 min) and then further purification via prep-TLC (50% EtOAc/petroleum ether) to obtain WX-02-30 (16.1 mg, 17% yield) as an off-white solid.

**$^1$H NMR** (400 MHz, $CD_3OD$, 323 K): δ 8.26 – 8.18 (m, 1H), 7.85 (d, *J* = 8.4 Hz, 1H), 7.63 (ddd, *J* = 8.5, 7.3, 1.9 Hz, 1H), 7.41 (dt, *J* = 7.7, 0.9 Hz, 1H), 7.26 (d, *J* = 8.1 Hz, 1H), 7.06 – 6.99 (m, 2H), 6.98 – 6.92 (m, 3H), 6.86 – 6.73 (m, 1H), 6.25 (s, 1H), 5.90 (s, 2H), 5.37 (dd, *J* = 5.9, 3.7 Hz, 1H), 4.27 (br. s, 1H), 3.68 – 3.34 (m, 2H), 2.47 – 2.35 (m, 1H), 2.27 – 2.12 (m, 2H), 1.89 (td, *J* = 7.5, 2.7 Hz, 2H), 1.76 – 1.55 (m, 2H), 1.47 (t, *J* = 7.5 Hz, 2H). 1 exchangeable proton not observed.

**HRMS** ESI-TOF m/z calculated for $C_{32}H_{29}N_6O_4$ $[M+H]^+$ 561.2250. Found 561.2272.

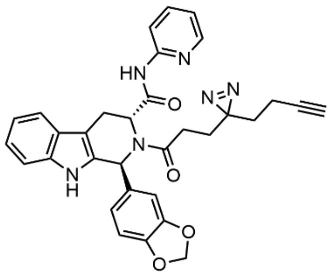

(1*S*,3*R*)-1-(benzo[*d*][1,3]dioxol-5-yl)-2-(3-(3-(but-3-yn-1-yl)-3*H*-diazirin-3-yl)propanoyl)-*N*-(pyridin-2-yl)-2,3,4,9-tetrahydro-1*H*-pyrido[3,4-*b*]indole-3-carboxamide (**WX-02-50**)

General Protocol A with prep-HPLC purification on a Phenomenex Unisil 3-100 C18 Ultra column (150 mm × 50 mm × 3 µm; mobile phase: [A: water containing 0.225% formic acid MeCN]; B%: 45–65%, 10 min) and then further purification on a Phenomenex Luna C18 column (150 mm × 25 mm × 10 µm; mobile phase: [A: water containing 0.225% formic acid–MeCN]; B%: 52–82%, 10 min) to obtain WX-02-50 (9.6 mg, 14% yield) as an off-white solid.

**$^1$H NMR** (400 MHz, $CD_3OD$, 323 K): δ 8.26 – 8.18 (m, 1H), 7.85 (d, *J* = 8.4 Hz, 1H), 7.64 (ddd, *J* = 8.4, 7.3, 1.9 Hz, 1H), 7.42 (dt, *J* = 7.8, 0.8 Hz, 1H), 7.26 (d, *J* = 8.1 Hz, 1H), 7.06 – 7.00 (m, 2H), 6.99 – 6.91 (m, 3H), 6.86 – 6.73 (m, 1H), 6.25 (s, 1H), 5.91 (s, 2H), 5.38 (dd, *J* = 5.9, 3.6 Hz, 1H), 4.30 (br. s, 1H, overlaps residual water), 3.62 – 3.34 (m, 2H), 2.48– 2.35 (m, 1H), 2.27 – 2.11 (m, 2H), 1.90 (td, *J* = 7.5, 2.6 Hz, 2H), 1.76 – 1.55 (m, 2H), 1.47 (t, *J* = 7.5 Hz, 2H). 1 exchangeable proton not observed.

**HRMS** ESI-TOF m/z calculated for $C_{32}H_{29}N_6O_4$ $[M+H]^+$ 561.2250. Found 561.2258.

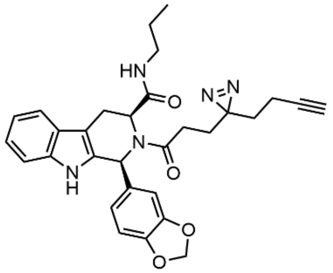

(1*S*,3*S*)-1-(benzo[*d*][1,3]dioxol-5-yl)-2-(3-(3-(but-3-yn-1-yl)-3*H*-diazirin-3-yl)propanoyl)-*N*-propyl-2,3,4,9-tetrahydro-1*H*-pyrido[3,4-*b*]indole-3-carboxamide (**WX-02-21**) – see ref (1).

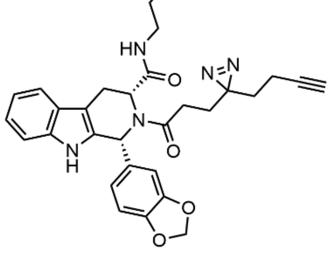

(1*R*,3*R*)-1-(benzo[*d*][1,3]dioxol-5-yl)-2-(3-(3-(but-3-yn-1-yl)-3*H*-diazirin-3-yl)propanoyl)-*N*-propyl-2,3,4,9-tetrahydro-1*H*-pyrido[3,4-*b*]indole-3-carboxamide (**WX-02-41**)

General Protocol A with prep-HPLC purification on a Phenomenex Gemini-NX C18 column (75 mm × 30 mm × 3 µm; mobile phase: [A: water containing 0.05% ammonium hydroxide–MeCN]; B%: 40–70%, 11.5 min) to obtain WX-02-41 (15.1 mg, 13% yield) as a yellow solid.

**$^1$H NMR** (400 MHz, $CD_3OD$, 297 K): δ 7.52 (d, *J* = 7.8 Hz, 1H), 7.27 (d, *J* = 8.0 Hz, 1H), 7.09 (t, *J* = 7.5 Hz, 1H), 7.03 (t, *J* = 7.4 Hz, 1H), 7.00 – 6.57 (m, 3H), 5.90 (s, 2H), 5.45 – 4.95 (m, 1H), 4.59 (br. s, 1H), 3.75 – 3.37 (m, 1H), 3.09 – 2.72 (m, 2H), 2.64 – 2.31 (m, 3H), 2.27 – 2.19 (m, 1H), 2.10 – 1.95 (m, 2H), 1.95 – 1.75 (m, 2H), 1.73 – 1.54 (m, 2H), 1.44 – 1.06 (m, 2H), 0.77 (br. s, 3H). 2 exchangeable protons not observed.

**HRMS** ESI-TOF m/z calculated for $C_{30}H_{32}N_5O_4$ $[M+H]^+$ 526.2454. Found 526.2462.

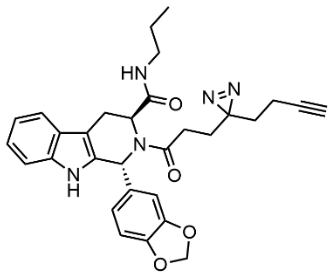


(1*R*,3*S*)-1-(benzo[*d*][1,3]dioxol-5-yl)-2-(3-(3-(but-3-yn-1-yl)-3*H*-diazirin-3-yl)propanoyl)-*N*-propyl-2,3,4,9-tetrahydro-1*H*-pyrido[3,4-*b*]indole-3-carboxamide (**WX-02-31**)

General Protocol A with $SiO_2$ flash chromatography purification (10–50% EtOAc/petroleum ether) and then prep-HPLC purification on a Phenomenex Luna C18 column (150 mm × 25 mm × 10 µm; mobile phase: [A: water containing 0.225% formic acid–MeCN]; B%: 43–73%, 10 min) to obtain WX-02-31 (22.3 mg, 32% yield) as an off-white solid.

**$^1$H NMR** (400 MHz, $CD_3OD$, 323 K): δ 7.39 (d, *J* = 7.7 Hz, 1H), 7.24 (d, *J* = 8.0 Hz, 1H), 7.02 (td, *J* = 7.0, 1.2 Hz, 1H), 6.96 (td, *J* = 7.3, 1.1 Hz, 1H), 6.94 – 6.87 (m, 2H), 6.84 – 6.66 (m, 1H), 6.17 (br. s, 1H), 5.87 (br. s, 2H), 5.52 – 4.88 (m, 1H), 3.56 – 3.32 (m, 2H), 3.00 (t, *J* = 6.9 Hz, 2H), 2.36 (ddd, *J* = 16.1, 8.9, 6.2 Hz, 1H), 2.19 (t, *J* = 2.6 Hz, 1H), 2.11 (ddd, *J* = 15.9, 8.8, 6.5 Hz, 1H), 1.92 (td, *J* = 7.4, 2.6 Hz, 2H), 1.83 – 1.38 (m, 4H), 1.29 (h, *J* = 7.2 Hz, 2H), 0.65 (t, *J* = 7.4 Hz, 3H). 2 exchangeable protons not observed.

**HRMS** ESI-TOF m/z calculated for $C_{30}H_{32}N_5O_4$ $[M+H]^+$ 526.2454. Found 526.2468.

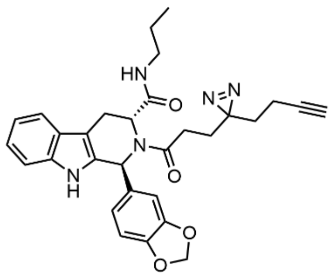


(1*S*,3*R*)-1-(benzo[*d*][1,3]dioxol-5-yl)-2-(3-(3-(but-3-yn-1-yl)-3*H*-diazirin-3-yl)propanoyl)-*N*-propyl-2,3,4,9-tetrahydro-1*H*-pyrido[3,4-*b*]indole-3-carboxamide (**WX-02-51**)

General Protocol A with prep-HPLC purification on a Phenomenex Luna C18 column (150 mm × 25 mm × 10 µm; mobile phase: [A: water containing 0.225% formic acid–MeCN]; B%: 53–83%, 10 min) to obtain WX-02-51 (24.7 mg, 49% yield) as an off-white solid.

**$^1$H NMR** (400 MHz, $CD_3OD$, 323 K): δ 7.39 (d, *J* = 7.7 Hz, 1H), 7.25 (d, *J* = 8.0 Hz, 1H), 7.03 (td, *J* = 6.9, 0.9 Hz, 1H), 6.96 (td, *J* = 6.9, 1.1 Hz, 1H), 6.94 – 6.61 (m, 3H), 6.17 (br. s, 1H), 5.88 (br. s, 2H), 5.51 – 4.88 (m, 1H), 3.56 – 3.33 (m, 2H), 3.00 (t, *J* = 6.9 Hz, 2H), 2.36 (ddd, *J* = 15.6, 8.9, 6.2 Hz, 1H), 2.19 (t, *J* = 2.6

Hz, 1H), 2.11 (ddd, *J* = 16.0, 8.8, 6.4 Hz, 1H), 1.98 – 1.87 (m, 2H), 1.82 – 1.37 (m, 4H), 1.30 (h, *J* = 7.2 Hz, 2H), 0.65 (t, *J* = 7.4 Hz, 3H). 2 exchangeable protons not observed.

**HRMS** ESI-TOF m/z calculated for $C_{30}H_{32}N_5O_4$ $[M+H]^+$ 526.2454. Found 526.2471.

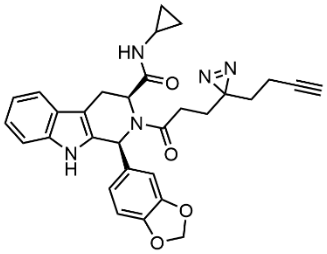

(1*S*,3*S*)-1-(benzo[*d*][1,3]dioxol-5-yl)-2-(3-(3-(but-3-yn-1-yl)-3*H*-diazirin-3-yl)propanoyl)-*N*-cyclopropyl-2,3,4,9-tetrahydro-1*H*-pyrido[3,4-*b*]indole-3-carboxamide (**WX-02-22**) – see ref (1).

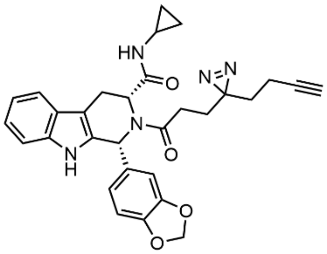

(1*R*,3*R*)-1-(benzo[*d*][1,3]dioxol-5-yl)-2-(3-(3-(but-3-yn-1-yl)-3*H*-diazirin-3-yl)propanoyl)-*N*-cyclopropyl-2,3,4,9-tetrahydro-1*H*-pyrido[3,4-*b*]indole-3-carboxamide (**WX-02-42**)

prep-HPLC purification on a Phenomenex Gemini-NX C18 column (75 mm × 30 mm × 3 μm; mobile phase: [A: water containing 0.05% ammonium hydroxide–MeCN]; B%: 35–65%, 7 min) to obtain WX-02-42 (17.1 mg, 24% yield) as a yellow solid.

**$^1$H NMR** (400 MHz, $CD_3OD$, 323 K): δ 7.51 (d, *J* = 7.8 Hz, 1H), 7.28 (d, *J* = 8.0 Hz, 1H), 7.09 (t, *J* = 7.6 Hz, 1H), 7.03 (t, *J* = 7.4 Hz, 1H), 6.94 – 6.77 (m, 2H), 6.74 (d, *J* = 8.0 Hz, 1H), 5.91 (s, 2H), 5.18 – 4.88 (m, 1H), 4.45 – 4.16 (m, 1H, overlaps residual water), 3.73 – 3.40 (m, 1H), 3.02 (dd, *J* = 15.9, 6.9 Hz, 1H), 2.52 – 2.32 (m, 2H), 2.32 – 2.07 (m, 2H), 2.07 – 1.97 (m, 2H), 1.97 – 1.72 (m, 2H), 1.63 (t, *J* = 7.2 Hz, 2H), 0.61 – 0.38 (m, 2H), 0.31 – 0.11 (m, 2H). 2 exchangeable protons not observed.

**HRMS** ESI-TOF m/z calculated for $C_{30}H_{30}N_5O_4$ $[M+H]^+$ 524.2298. Found 524.2307.

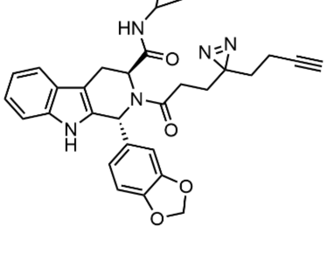

(1*R*,3*S*)-1-(benzo[*d*][1,3]dioxol-5-yl)-2-(3-(3-(but-3-yn-1-yl)-3*H*-diazirin-3-yl)propanoyl)-*N*-cyclopropyl-2,3,4,9-tetrahydro-1*H*-pyrido[3,4-*b*]indole-3-carboxamide (**WX-02-32**)

General Protocol A with $SiO_2$ flash chromatography purification (10–50% EtOAc/petroleum ether) and then prep-HPLC purification on a Phenomenex Luna C18 column (150 mm × 25 mm × 10 μm; mobile phase: [A:

water containing 0.225% formic acid–MeCN]; B%: 47–77%, 10 min) to obtain WX-02-32 (22.2 mg, 32% yield) as an off-white solid.

**$^1$H NMR** (400 MHz, $CD_3OD$, 323 K): δ 7.39 (d, *J* = 7.8 Hz, 1H), 7.25 (d, *J* = 8.0 Hz, 1H), 7.03 (t, *J* = 7.5 Hz, 1H), 6.97 (t, *J* = 7.6 Hz, 1H), 6.94 – 6.84 (m, 2H), 6.82 – 6.61 (m, 1H), 6.26 – 6.06 (m, 1H), 5.87 (br. s, 2H), 5.50 – 4.94 (m, 1H), 3.48 – 3.33 (m, 2H, overlaps trace solvent), 2.50 – 2.41 (m, 1H), 2.40 – 2.27 (m, 1H), 2.19 (q, *J* = 2.4 Hz, 1H), 2.10 (dt, *J* = 16.0, 7.7 Hz, 1H), 1.93 (td, *J* = 7.4, 1.8 Hz, 2H), 1.81 – 1.29 (m, 4H), 0.65 – 0.47 (m, 2H), 0.42 – 0.29 (m, 1H), 0.26 – 0.10 (m, 1H). 2 exchangeable protons not observed.

**HRMS** ESI-TOF m/z calculated for $C_{30}H_{30}N_5O_4$ $[M+H]^+$ 524.2298. Found 524.2314.

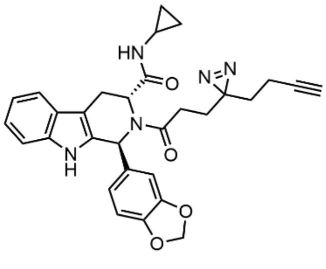

(1*S*,3*R*)-1-(benzo[*d*][1,3]dioxol-5-yl)-2-(3-(3-(but-3-yn-1-yl)-3*H*-diazirin-3-yl)propanoyl)-*N*-cyclopropyl-2,3,4,9-tetrahydro-1*H*-pyrido[3,4-*b*]indole-3-carboxamide (**WX-02-52**)

General Protocol A with prep-HPLC purification on a Phenomenex Luna C18 column (150 mm × 25 mm × 10 µm; mobile phase: [A: water containing 0.225% formic acid–MeCN]; B%: 50–80%, 10 min) to obtain WX-02-52 (24.9 mg, 50% yield) as an off-white solid.

**$^1$H NMR** (400 MHz, $CD_3OD$, 323 K): δ 7.39 (d, *J* = 7.8 Hz, 1H), 7.25 (d, *J* = 8.0 Hz, 1H), 7.03 (t, *J* = 7.2 Hz, 1H), 6.98 (t, *J* = 7.5 Hz, 1H), 6.94 – 6.86 (m, 2H), 6.85 – 6.67 (br. s, 1H), 6.26 – 6.06 (m, 1H), 5.88 (s, 2H), 5.36 – 5.02 (m, 1H), 4.28 (br. s, 1H), 3.48 – 3.33 (m, 2H, overlaps trace solvent), 2.45 (tt, *J* = 7.2, 3.8 Hz, 1H), 2.34 (ddd, *J* = 15.5, 8.8, 6.2 Hz, 1H), 2.19 (t, *J* = 2.6 Hz, 1H), 2.11 (ddt, *J* = 15.8, 7.9 Hz, 1H), 1.93 (td, *J* = 7.5, 2.5 Hz, 2H), 1.81 – 1.29 (m, 4H), 0.65 – 0.47 (m, 2H), 0.42 – 0.29 (m, 1H), 0.26 – 0.10 (m, 1H). 1 exchangeable proton not observed.

**HRMS** ESI-TOF m/z calculated for $C_{30}H_{30}N_5O_4$ $[M+H]^+$ 524.2298. Found 524.2318.

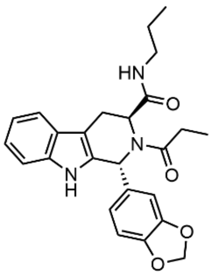

(1*R*,3*S*)-1-(benzo[*d*][1,3]dioxol-5-yl)-2-propionyl-*N*-propyl-2,3,4,9-tetrahydro-1*H*-pyrido[3,4-*b*]indole-3-carboxamide (**WX-02-654**)

General Protocol C with prep-HPLC purification on a Waters Xbridge BEH C18 column (150 mm × 25 mm × 5 µm; mobile phase: [A: water containing 10 mM ammonium bicarbonate–MeCN]; B%: 34–64%, 9 min) to obtain WX-02-654 (51.0 mg, 30% yield) as a white solid.

**$^1$H NMR** (400 MHz, $CD_3OD$, 298 K): δ 7.39 (d, *J* = 8.0 Hz, 1H), 7.29 – 7.16 (m, 1H), 7.07 – 7.00 (m, 1H), 7.00 – 6.85 (m, 3H), 6.82 – 6.62 (m, 1H), 6.30 – 6.14 (m, 1H), 5.96 – 5.77 (m, 2H), 5.32 (br. s, 1H), 5.13 (br. s, 0.5H), 4.59 (br. s, 0.5H), 3.56 – 3.37 (m, 1H), 3.00 (t, *J* = 6.9 Hz, 2H), 2.64 – 2.49 (m, 1H), 2.39 –

2.13 (m, 1H), 1.38 – 1.17 (m, 2H), 1.10 – 0.91 (m, 3H), 0.79 – 0.55 (m, 3H). 2 exchangeable protons not observed.

**HRMS** ESI-TOF m/z calculated for $C_{25}H_{28}N_3O_4$ $[M+H]^+$ 434.2080. Found 434.2092.

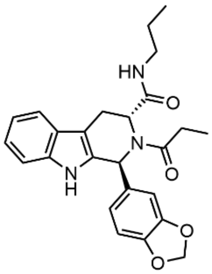

(1*S*,3*R*)-1-(benzo[*d*][1,3]dioxol-5-yl)-2-propionyl-*N*-propyl-2,3,4,9-tetrahydro-1*H*-pyrido[3,4-*b*]indole-3-carboxamide (**WX-02-655**)

General Protocol C with prep-HPLC purification on a Waters Xbridge BEH C18 column (150 mm × 25 mm × 5 μm; mobile phase: [A: water containing 10 mM ammonium bicarbonate–MeCN]; B%: 35–65%, 9 min) to obtain WX-02-654 (91.0 mg, 79% yield) as a white solid.

**$^1$H NMR** (400 MHz, $CD_3OD$, 299 K): δ 7.39 (d, *J* = 8.0 Hz, 1H), 7.29 – 7.17 (m, 1H), 7.07 – 7.00 (m, 1H), 7.00 – 6.84 (m, 3H), 6.82 – 6.63 (m, 1H), 6.30 – 6.12 (m, 1H), 5.96 – 5.76 (m, 2H), 5.32 (br. s, 1H), 5.13 (br. s, 0.5H), 4.58 (br. s, 0.5H), 3.60 – 3.35 (m, 1H), 3.07 – 2.91 (m, 2H), 2.64 – 2.49 (m, 1H), 2.39 – 2.13 (m, 1H), 1.41 – 1.18 (m, 2H), 1.14 – 0.89 (m, 3H), 0.76 – 0.50 (m, 3H). 2 exchangeable protons not observed.

**HRMS** ESI-TOF m/z calculated for $C_{25}H_{28}N_3O_4$ $[M+H]^+$ 434.2080. Found 434.2093.

**NMR spectra**

[1]H NMR spectrum of WX-02-222 (400 MHz, $CD_3OD$, 297 K)

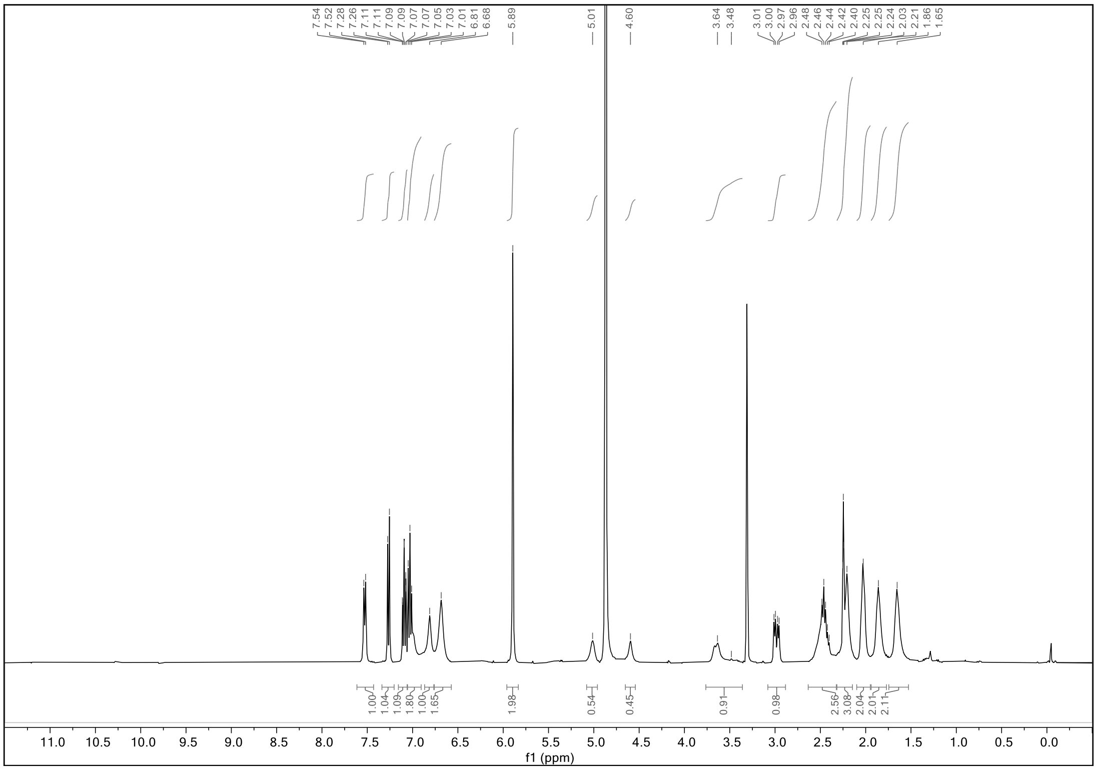

[1]H NMR spectrum of WX-02-223 (400 MHz, $CD_3OD$, 323 K)

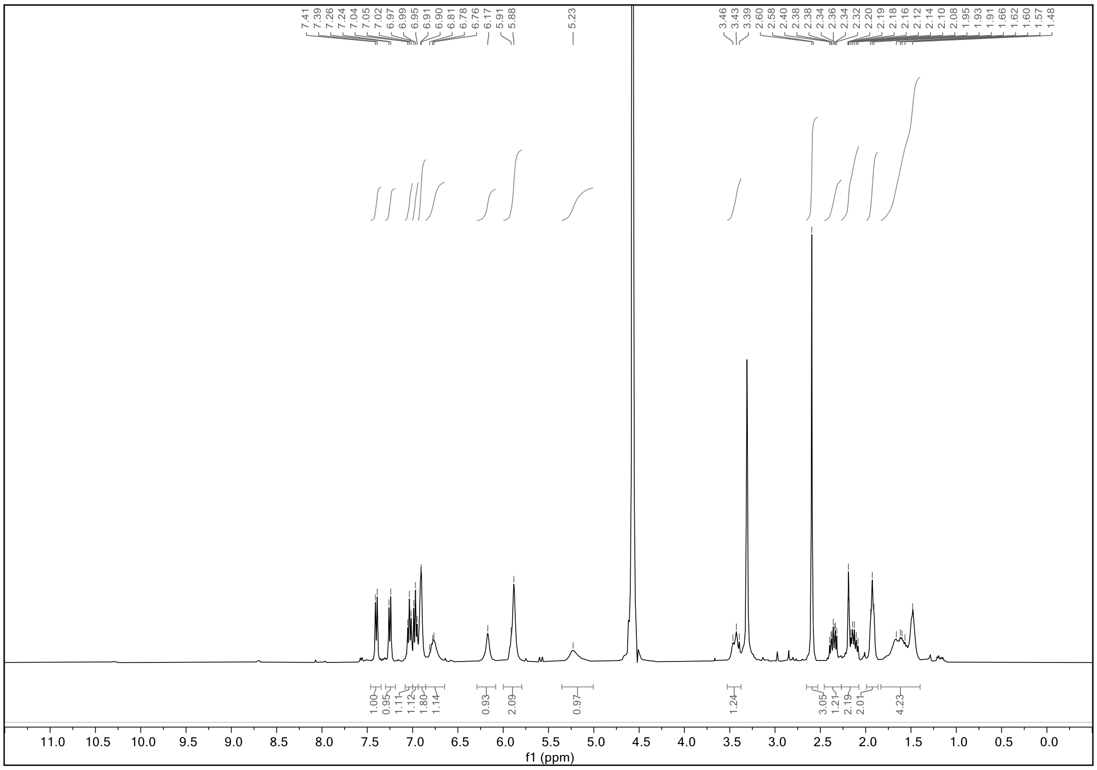

[1]H NMR spectrum of WX-02-224 (400 MHz, $CD_3OD$, 323 K)

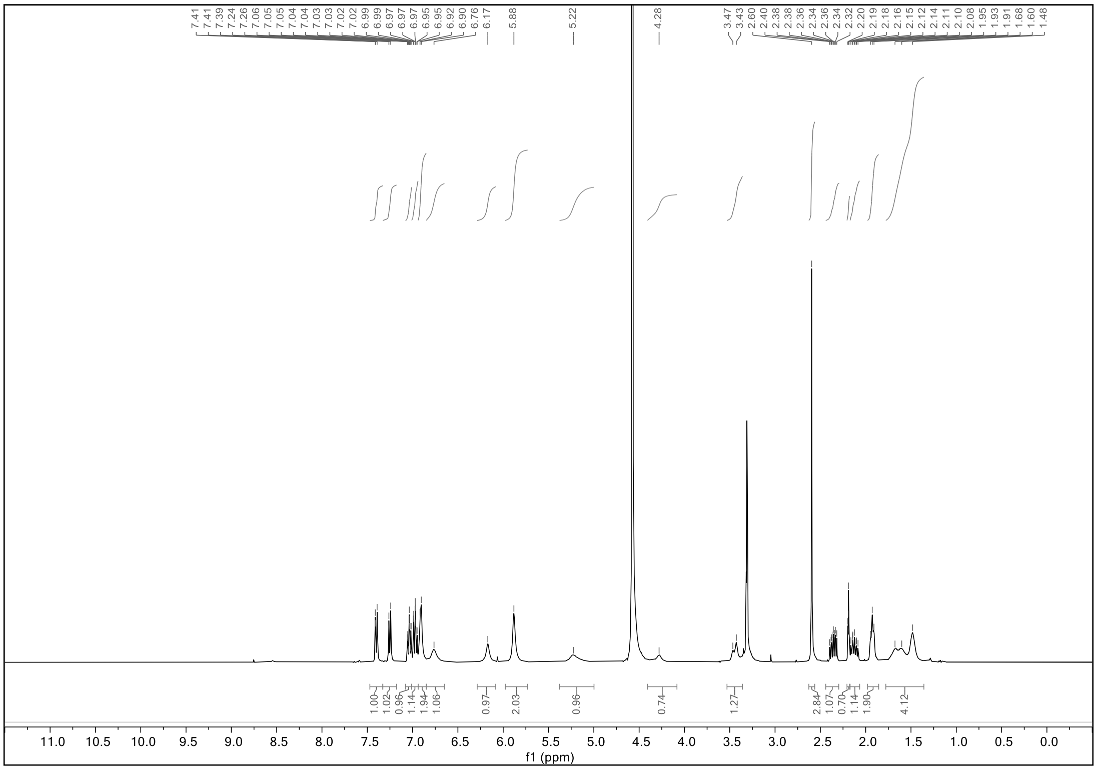

[1]H NMR spectrum of WX-02-39 (400 MHz, $CD_3OD$, 298 K)

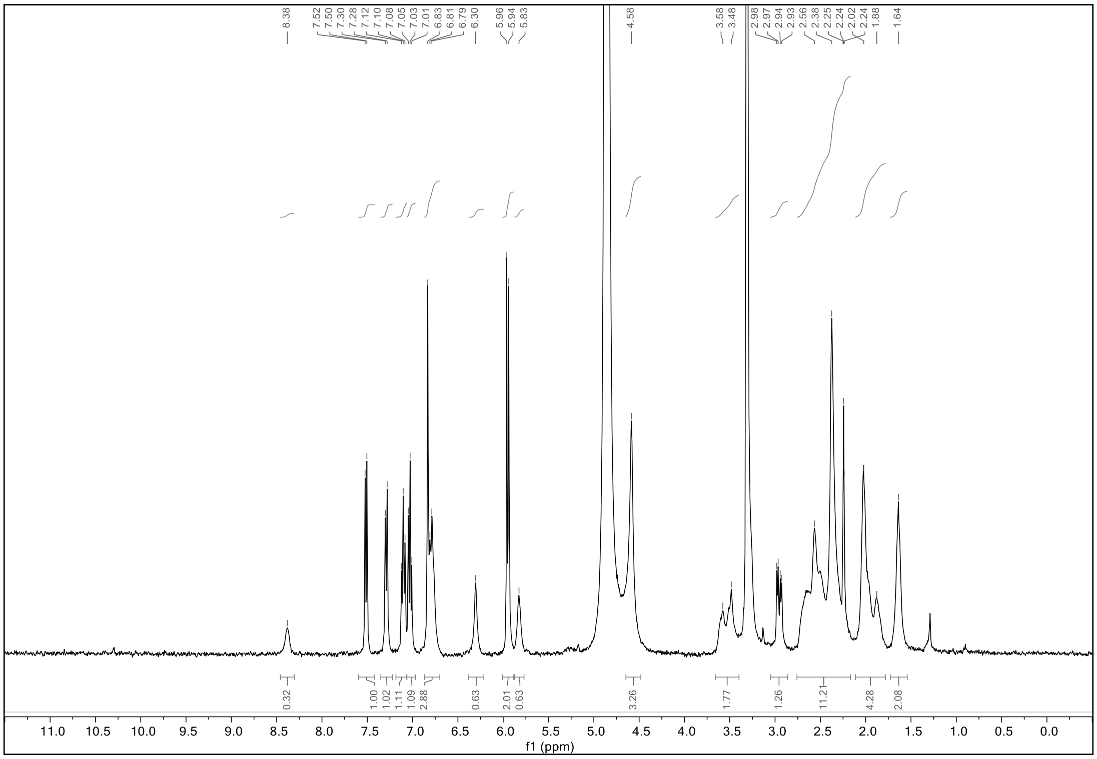

$^{1}H$ NMR of WX-02-29 (400 MHz, $CD_3OD$, 323 K)

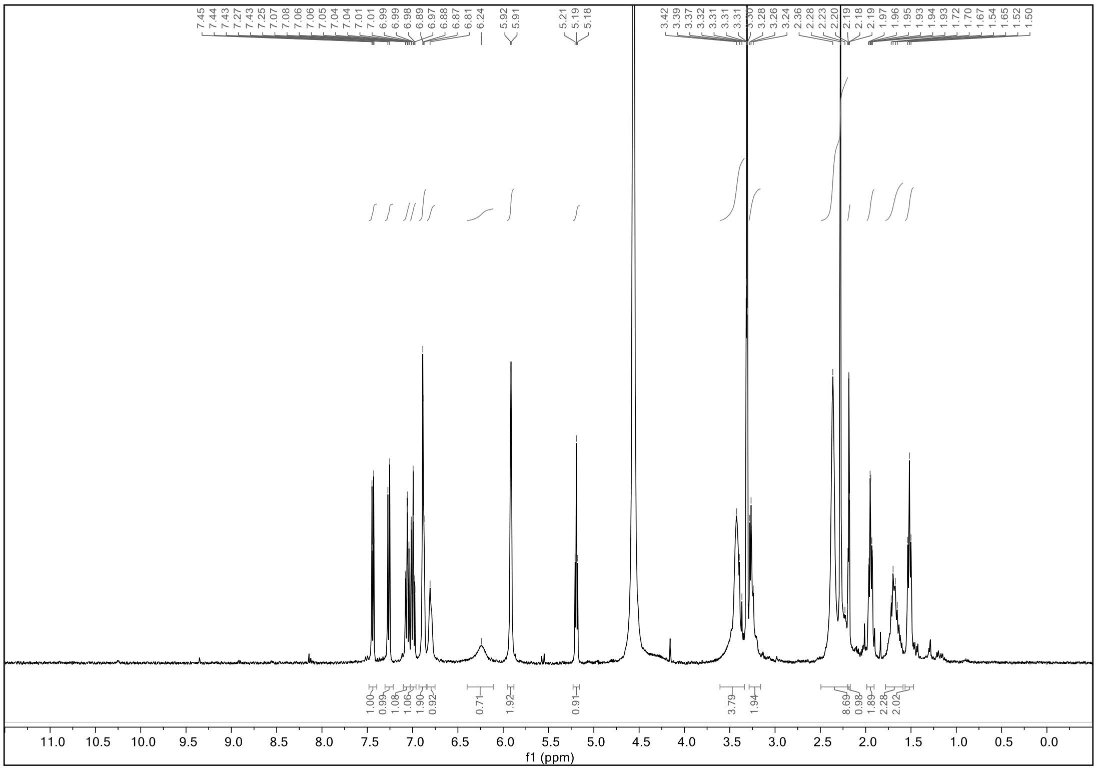

$^{1}H$ NMR spectrum of WX-02-49 (400 MHz, $CD_3OD$, 323 K)

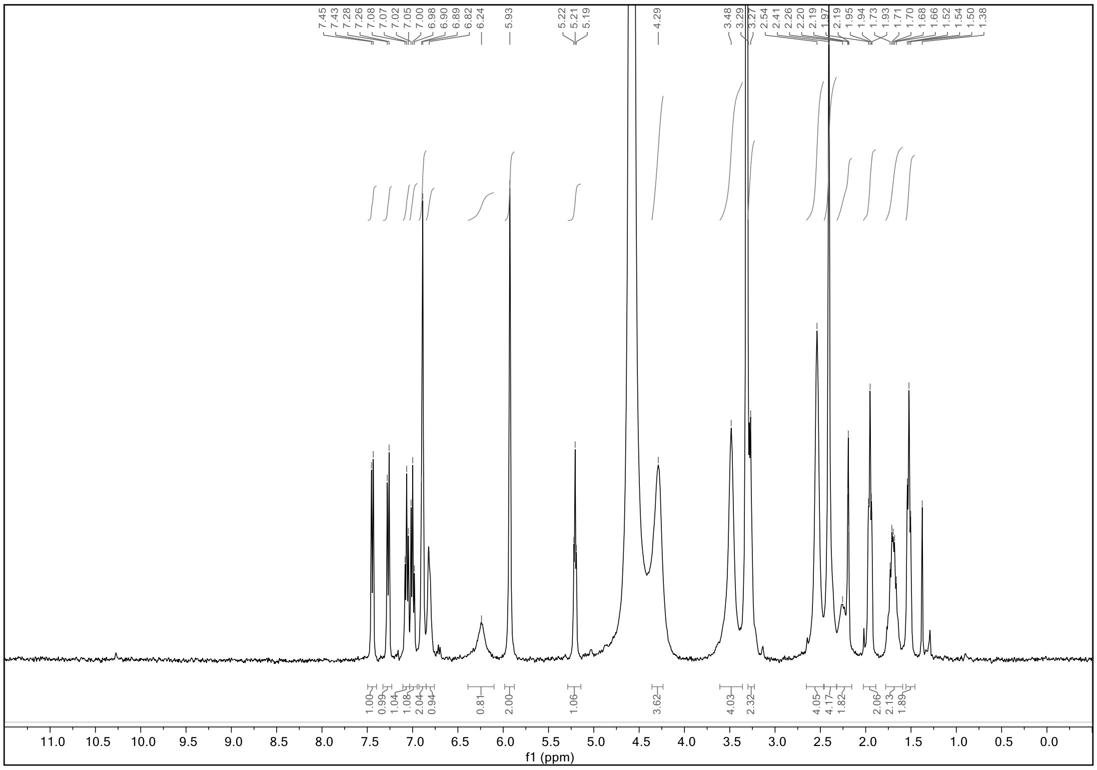

[1]H NMR spectrum of WX-02-40 (400 MHz, $CD_3OD$, 323 K)

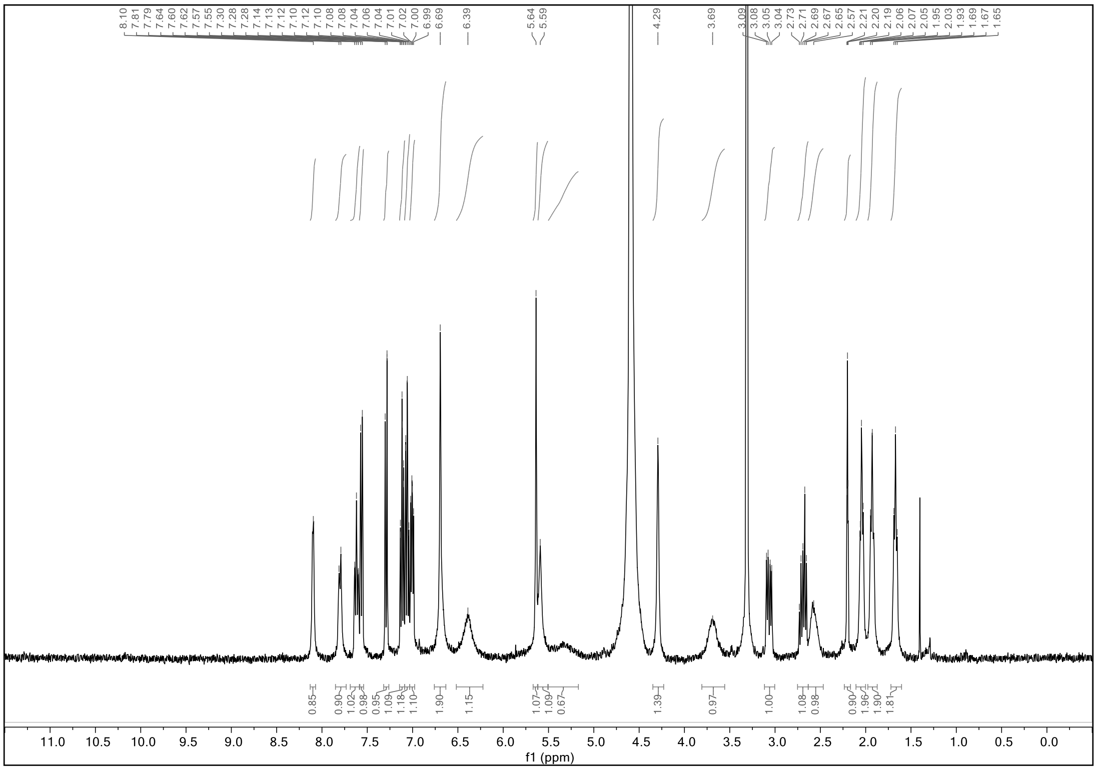

[1]H NMR spectrum of WX-02-30 (400 MHz, $CD_3OD$, 323 K)

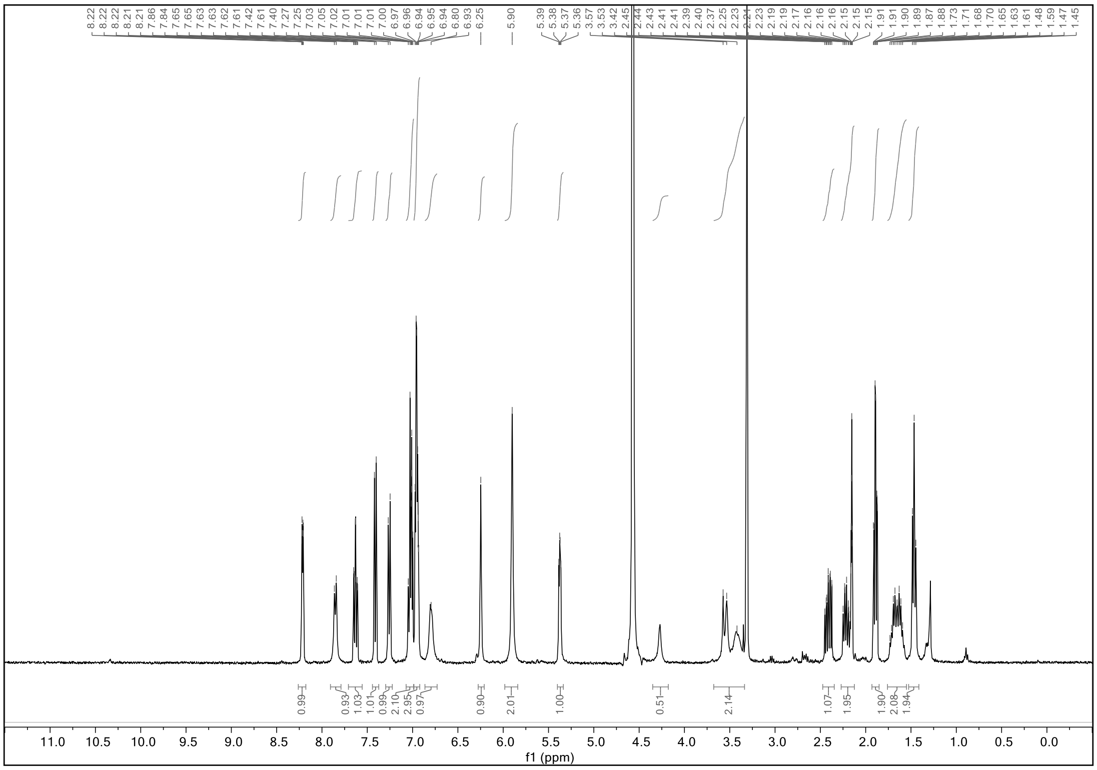

[1]H NMR spectrum of WX-02-50 (400 MHz, $CD_3OD$, 323 K)

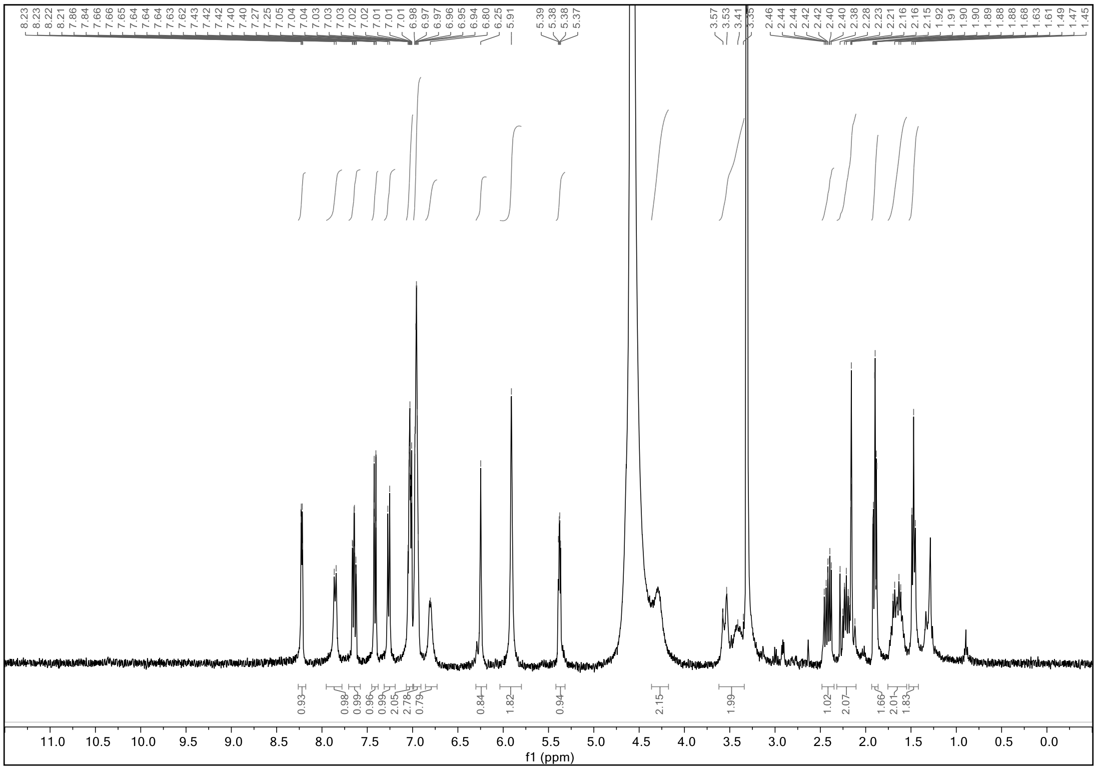

[1]H NMR spectrum of WX-02-41 (400 MHz, $CD_3OD$, 297 K)

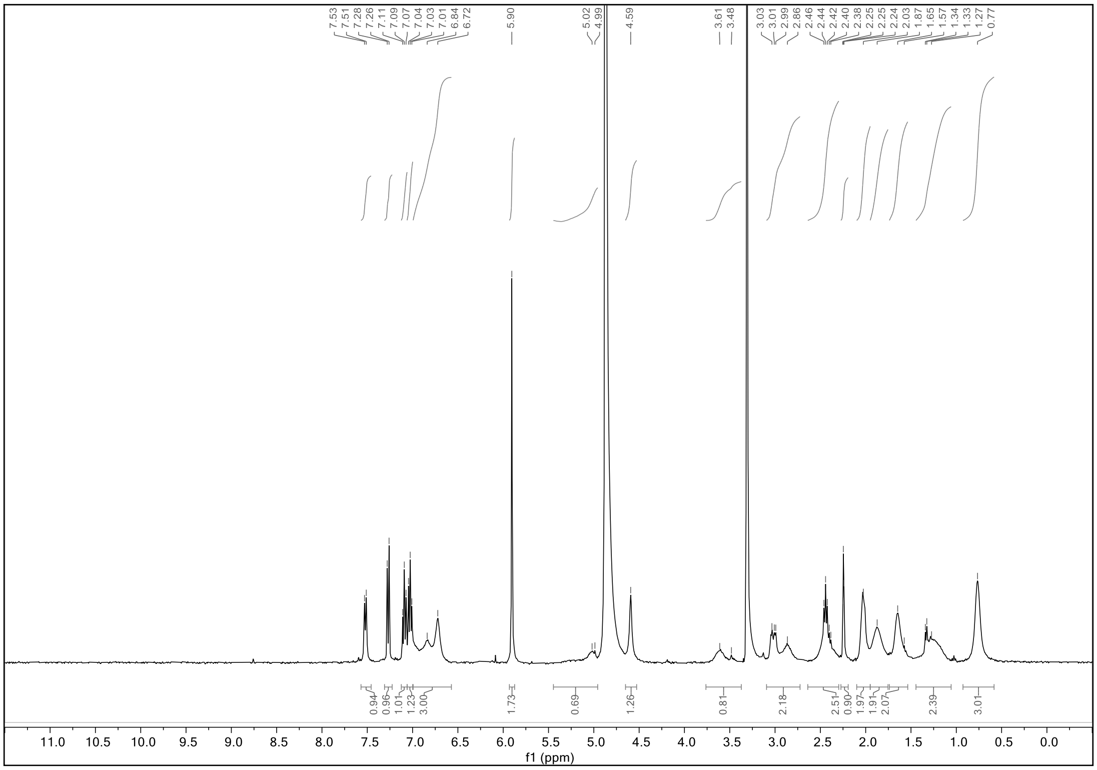

[1]H NMR spectrum of WX-02-31 (400 MHz, $CD_3OD$, 323 K)

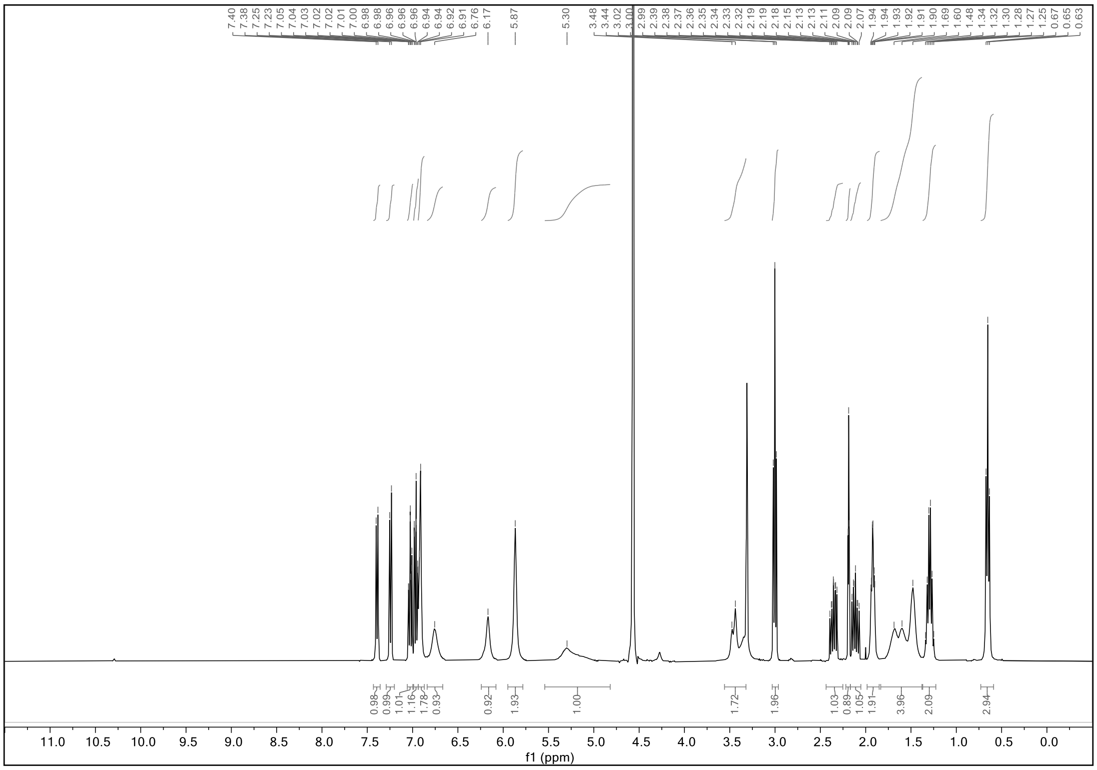

[1]H NMR spectrum of WX-02-51 (400 MHz, $CD_3OD$, 323 K)

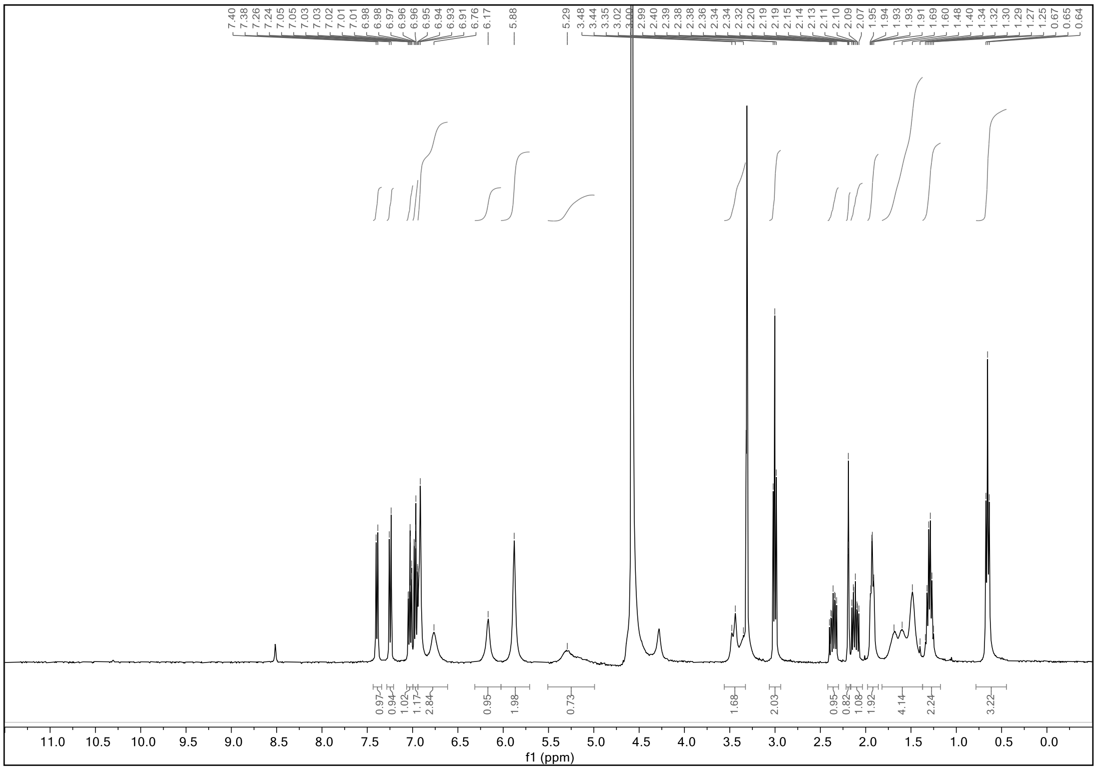

[1]H NMR spectrum of WX-02-42 (400 MHz, $CD_3OD$, 323 K)

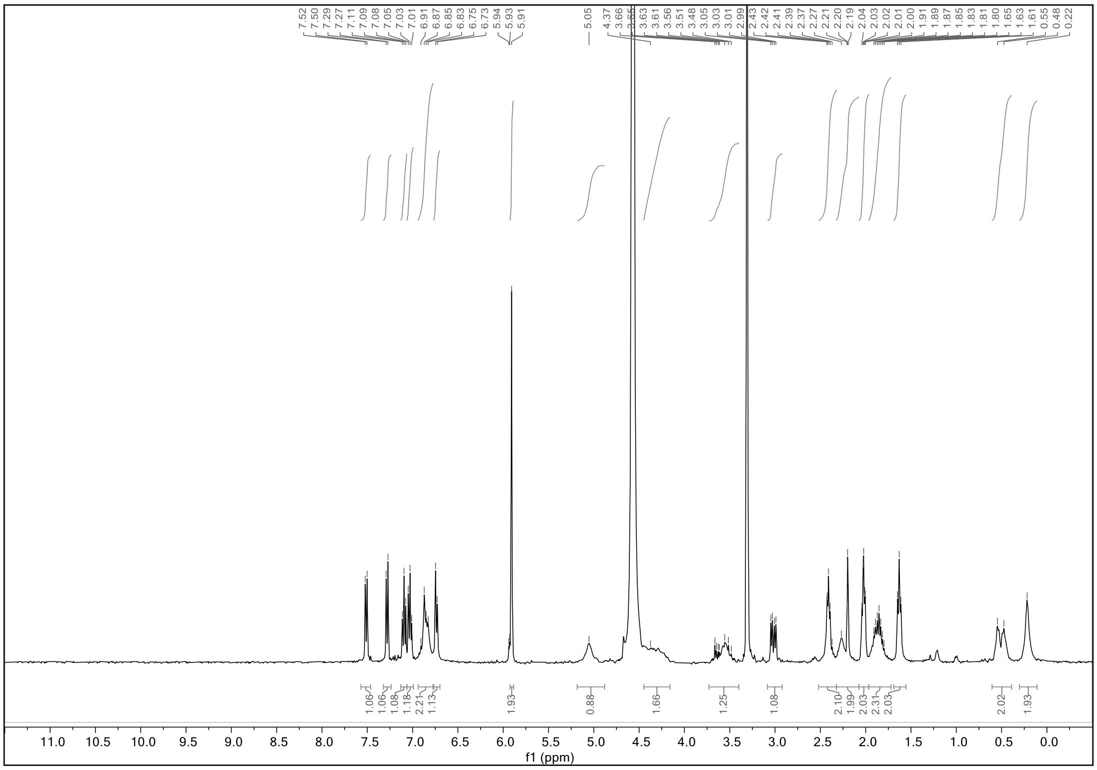

$^1$H NMR spectrum of WX-02-32 (400 MHz, $CD_3OD$, 323 K)

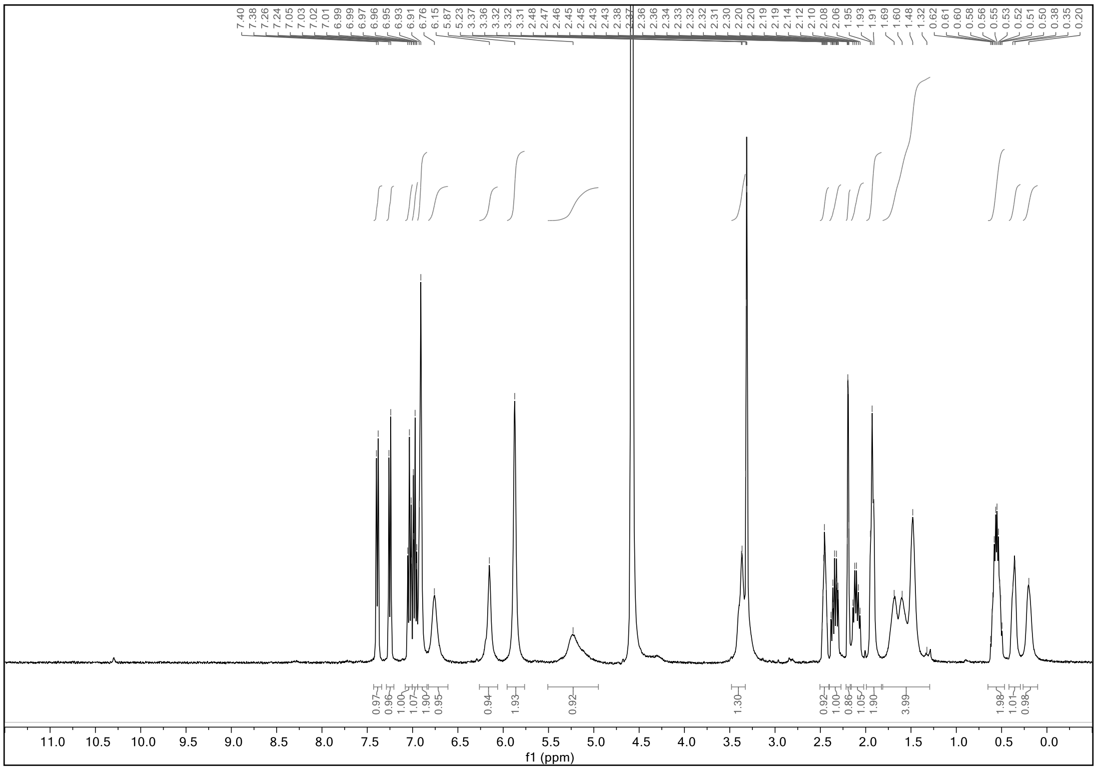

$^1$H NMR spectrum of WX-02-52 (400 MHz, $CD_3OD$, 323 K)

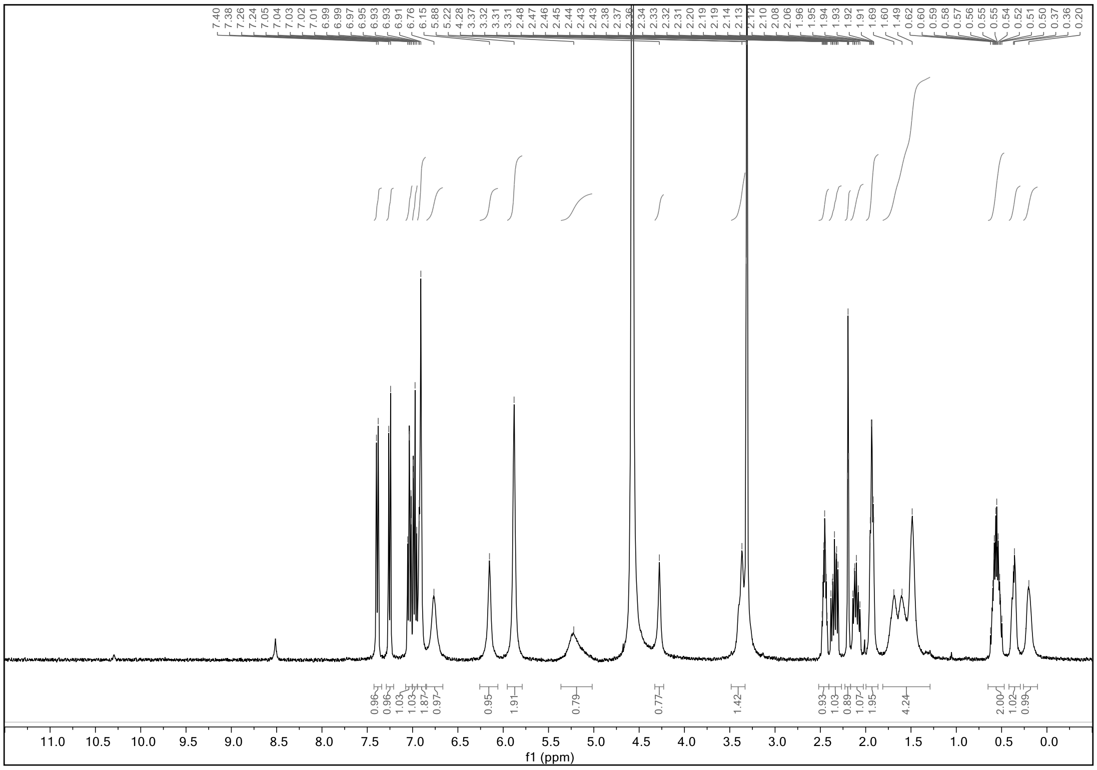

$^{1}H$ NMR spectrum of WX-02-654 (400 MHz, $CD_3OD$, 298 K)

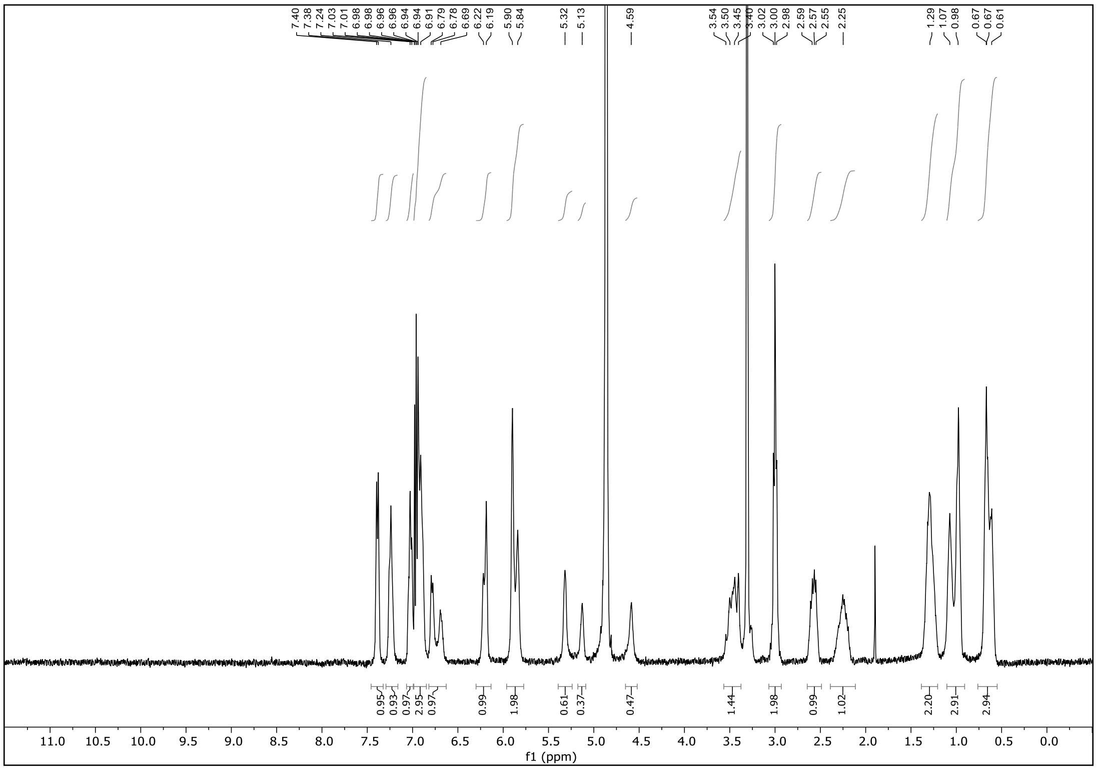

$^{1}H$ NMR spectrum of WX-02-655 (400 MHz, $CD_3OD$, 299 K)

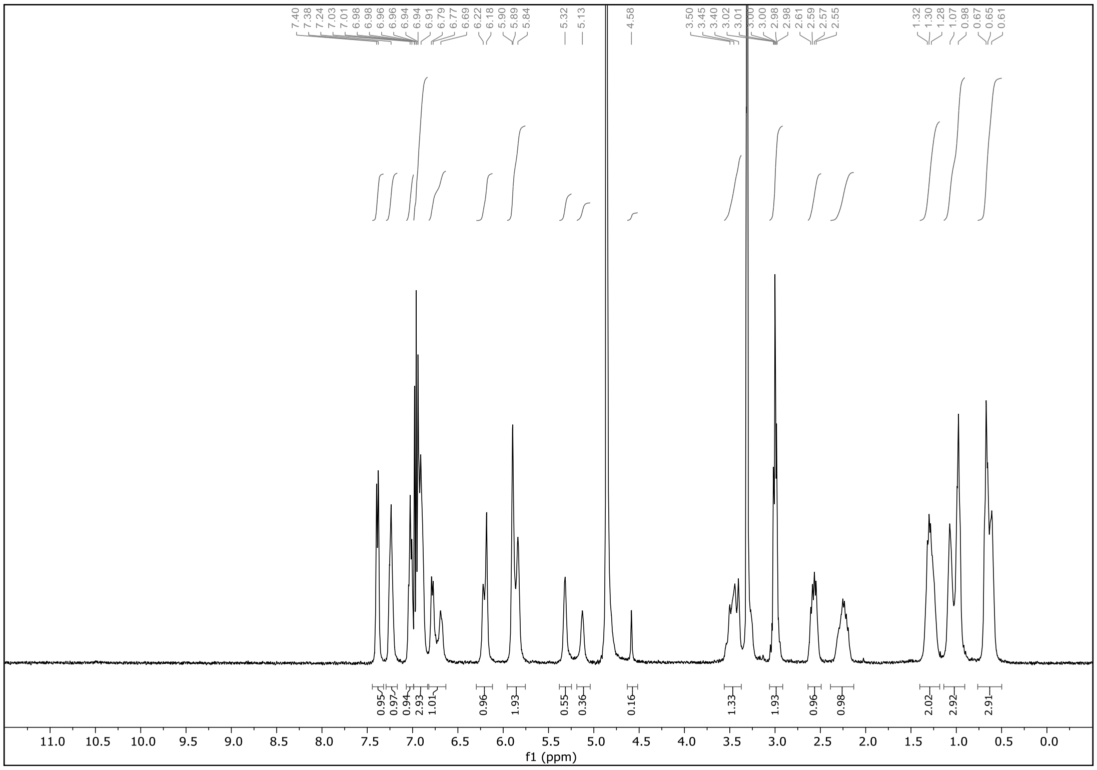

**Chiral stationary phase SFC**

WX-02-221

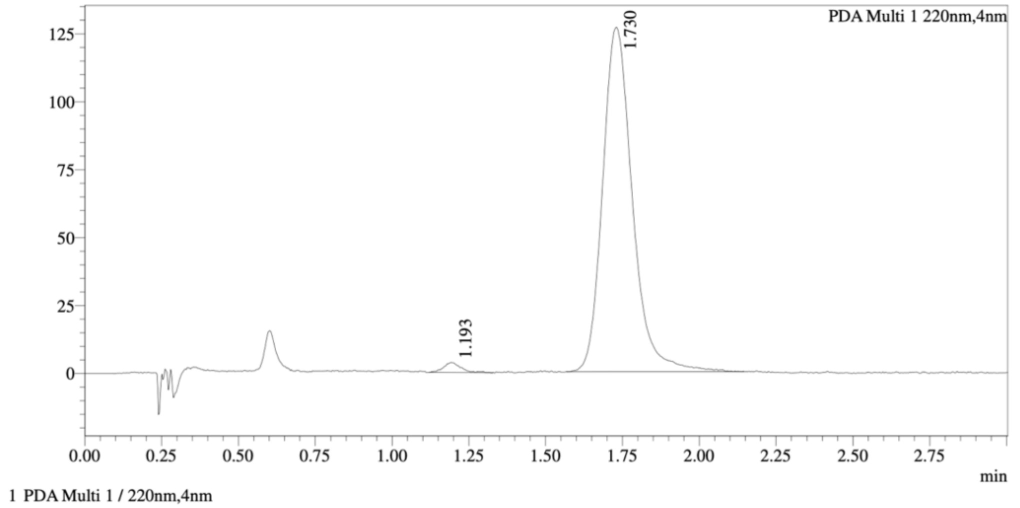


Integration Result

Peak Table

PDA Ch1 220nm

| Peak# | Ret. Time | Height | Height% | Resolution(USP) | Area | Area% |
|---|---|---|---|---|---|---|
| 1 | 1.193 | 3570 | 2.794 | -- | 14204 | 1.633 |
| 2 | 1.730 | 124173 | 97.206 | 3.939 | 855705 | 98.367 |

WX-02-222

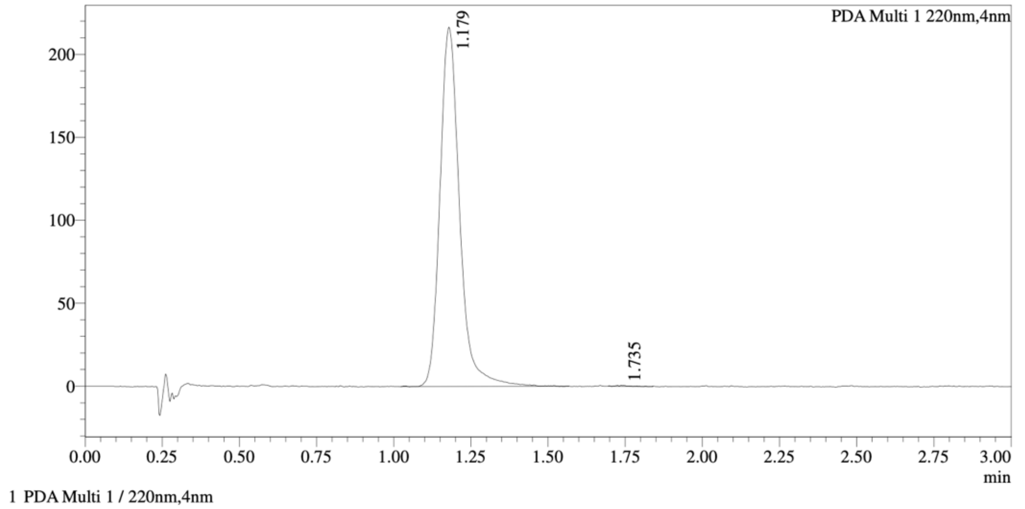


Integration Result

Peak Table

PDA Ch1 220nm

| Peak# | Ret. Time | Height | Height% | Resolution(USP) | Area | Area% |
|---|---|---|---|---|---|---|
| 1 | 1.179 | 213197 | 99.833 | -- | 960230 | 99.877 |
| 2 | 1.735 | 357 | 0.167 | 7.324 | 1187 | 0.123 |

Mixture of WX-02-221 & WX-02-222

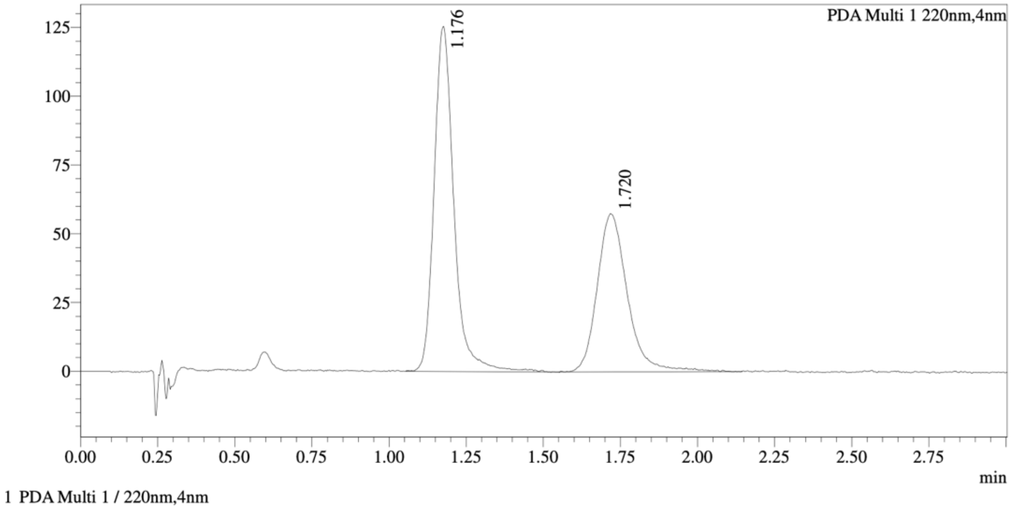


==========================================================================================
Integration Result
==========================================================================================

Peak Table

PDA Ch1 220nm

| Peak# | Ret. Time | Height | Height% | Resolution(USP) | Area | Area% |
|---|---|---|---|---|---|---|
| 1 | 1.176 | 123365 | 68.384 | -- | 559365 | 58.854 |
| 2 | 1.720 | 57035 | 31.616 | 3.808 | 391066 | 41.146 |

Method details

Column: Chiralpak AD-3, 50 mm length × 4.6 mm internal diameter, 3 μm particle size.
Mobile phase: A: supercritical $CO_2$; B: *i*-PrOH (0.05% diethylamine).
Isocratic elution: 40% B.

WX-02-223

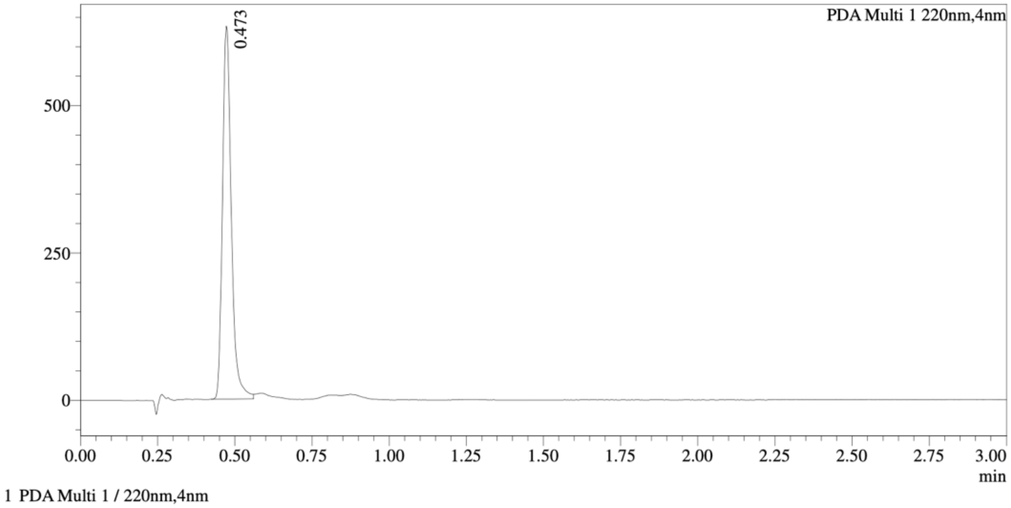


Integration Result

Peak Table

PDA Ch1 220nm

| Peak# | Ret. Time | Height | Height% | Resolution(USP) | Area | Area% |
|---|---|---|---|---|---|---|
| 1 | 0.473 | 599695 | 100.000 | -- | 1259867 | 100.000 |

WX-02-224

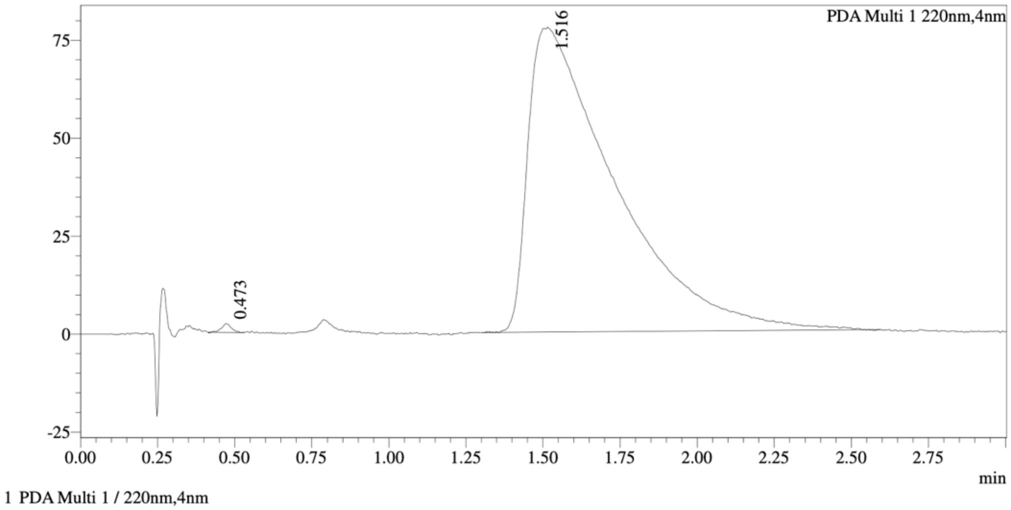


Integration Result

Peak Table

PDA Ch1 220nm

| Peak# | Ret. Time | Height | Height% | Resolution(USP) | Area | Area% |
|---|---|---|---|---|---|---|
| 1 | 0.473 | 2100 | 2.634 | -- | 4944 | 0.323 |
| 2 | 1.516 | 77629 | 97.366 | 3.581 | 1527995 | 99.677 |

Mixture of WX-02-223 & WX-02-224

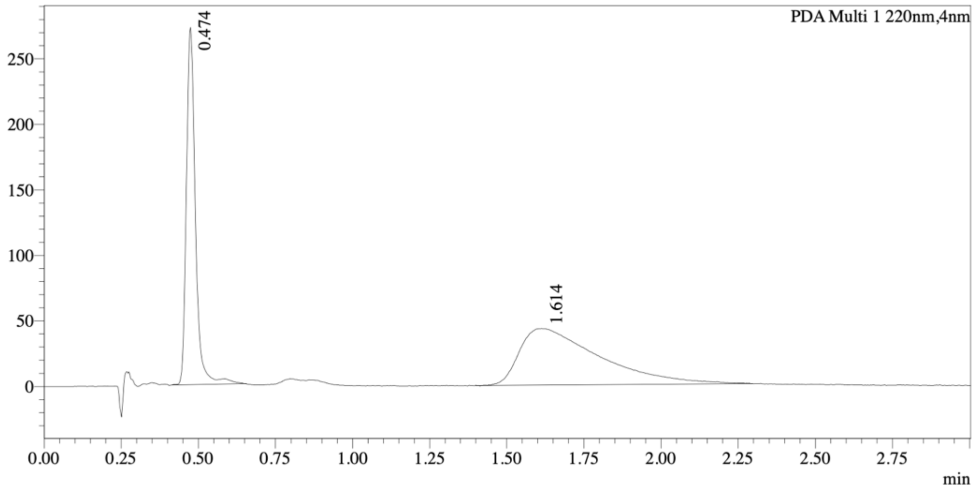


1 PDA Multi 1 / 220nm,4nm

Integration Result

Peak Table

PDA Ch1 220nm

| Peak# | Ret. Time | Height | Height% | Resolution(USP) | Area | Area% |
|---|---|---|---|---|---|---|
| 1 | 0.474 | 260385 | 85.816 | -- | 555514 | 41.695 |
| 2 | 1.614 | 43039 | 14.184 | 4.473 | 776799 | 58.305 |

Method details

Column: Chiralcel OD-3, 50 mm length × 4.6 mm internal diameter, 3 μm particle size.
Mobile phase: A: supercritical $CO_2$; B: MeOH (0.05% diethylamine).
Isocratic elution: 40% B.

WX-02-19

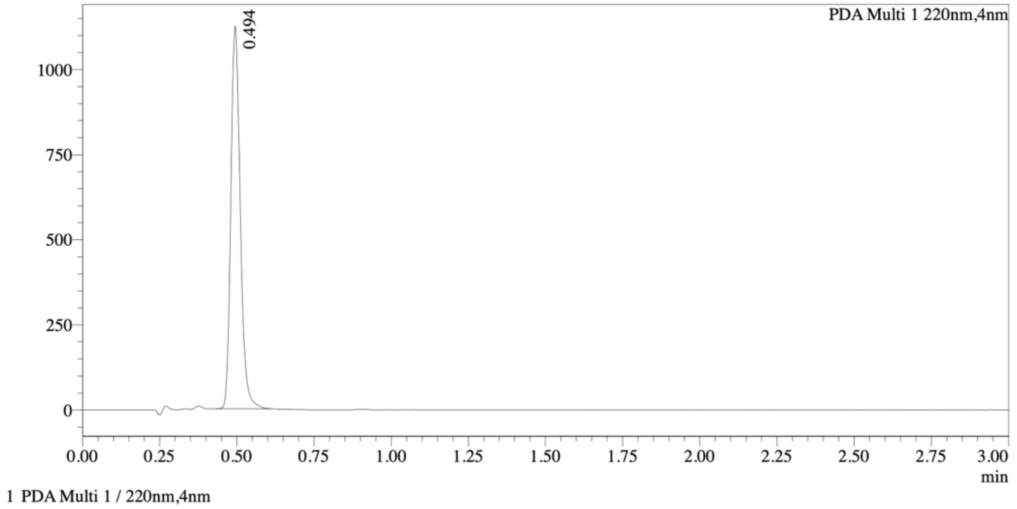


==================================================================================

Integration Result

==================================================================================

Peak Table

PDA Ch1 220nm

| Peak# | Ret. Time | Height | Height% | Resolution(USP) | Area | Area% |
|---|---|---|---|---|---|---|
| 1 | 0.494 | 1109274 | 100.000 | -- | 2387206 | 100.000 |

WX-02-39

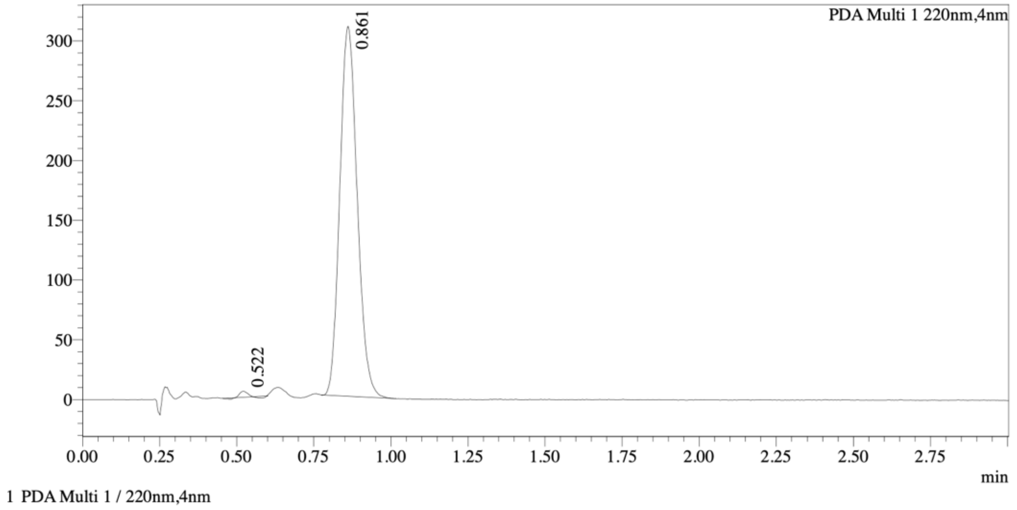


==================================================================================

Integration Result

==================================================================================

Peak Table

PDA Ch1 220nm

| Peak# | Ret. Time | Height | Height% | Resolution(USP) | Area | Area% |
|---|---|---|---|---|---|---|
| 1 | 0.522 | 4691 | 1.498 | -- | 5408 | 0.443 |
| 2 | 0.861 | 308375 | 98.502 | 4.254 | 1216246 | 99.557 |

Mixture of WX-02-19 & WX-02-39

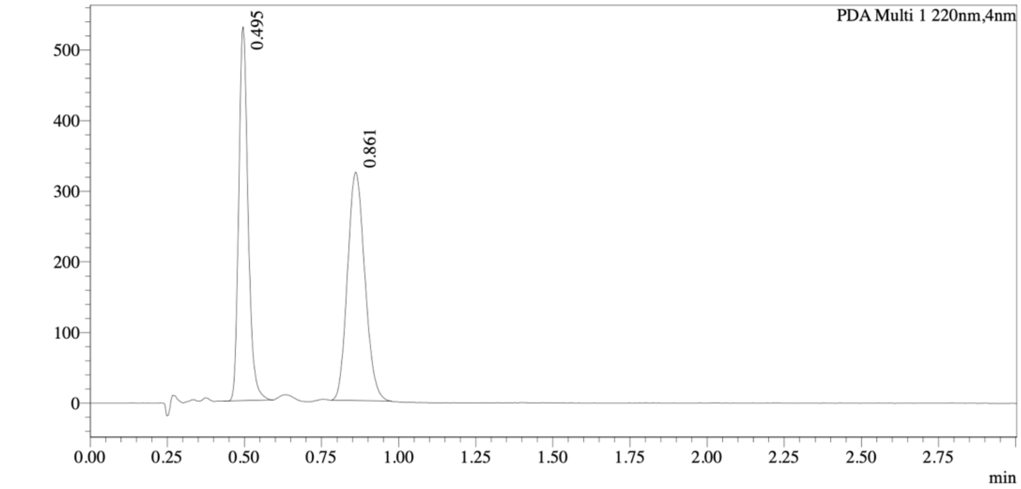


Integration Result

Peak Table

PDA Ch1 220nm

| Peak# | Ret. Time | Height | Height% | Resolution(USP) | Area | Area% |
|---|---|---|---|---|---|---|
| 1 | 0.495 | 517033 | 61.583 | -- | 1114513 | 46.806 |
| 2 | 0.861 | 322541 | 38.417 | 4.459 | 1266627 | 53.194 |

Method details

Column: Chiralcel OD-3, 50 mm length × 4.6 mm internal diameter, 3 μm particle size.
Mobile phase: A: supercritical $CO_2$; B: 2:1 MeOH/MeCN (0.05% diethylamine).
Isocratic elution: 40% B.

WX-02-20

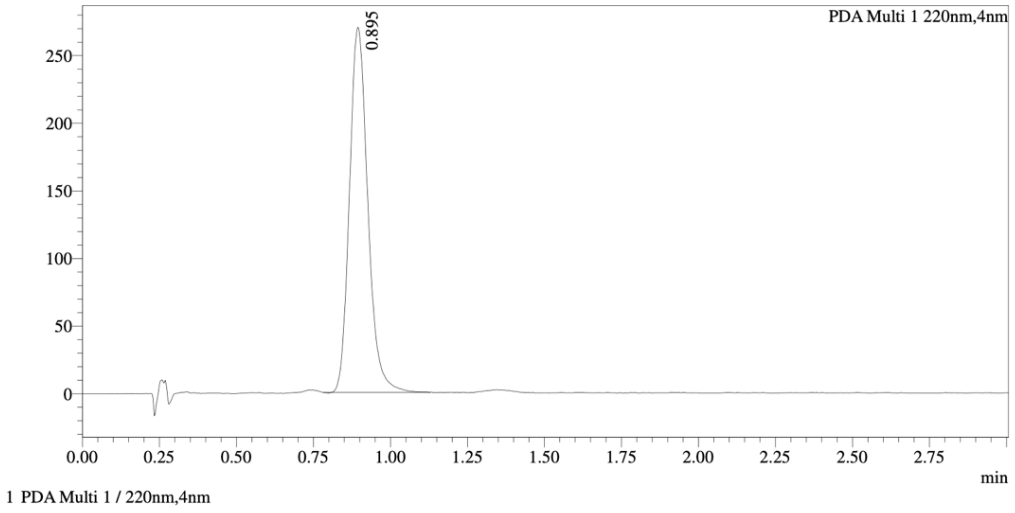

| | |
|---|---|
| | Integration Result |

Peak Table

PDA Ch1 220nm

| Peak# | Ret. Time | Height | Height% | Resolution(USP) | | Area | Area% |
|---|---|---|---|---|---|---|---|
| 1 | 0.895 | 269009 | 100.000 | | -- | 1120398 | 100.000 |

WX-02-40

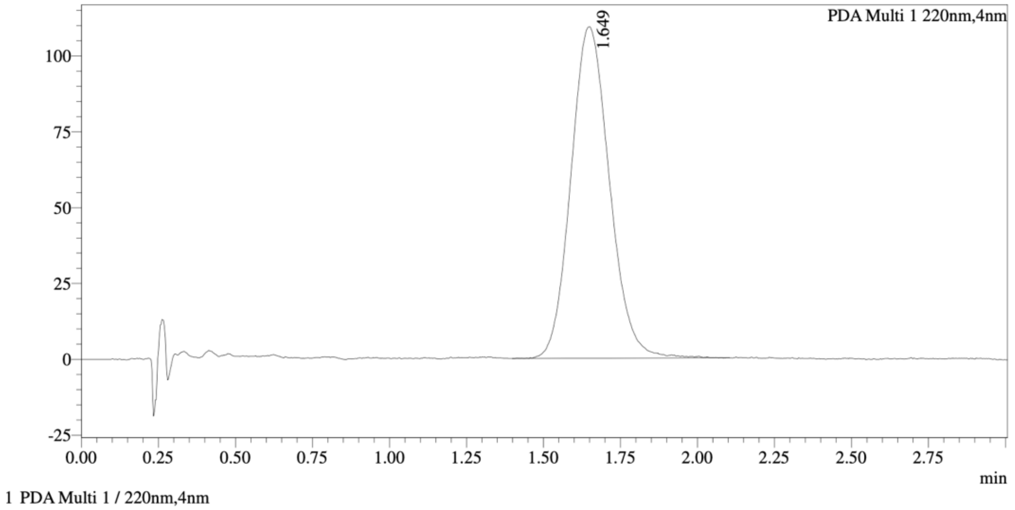

Integration Result

Peak Table

PDA Ch1 220nm

| Peak# | Ret. Time | Height | Height% | Resolution(USP) | | Area | Area% |
|---|---|---|---|---|---|---|---|
| 1 | 1.649 | 109016 | 100.000 | | -- | 952325 | 100.000 |

Mixture of WX-02-20 & WX-02-40

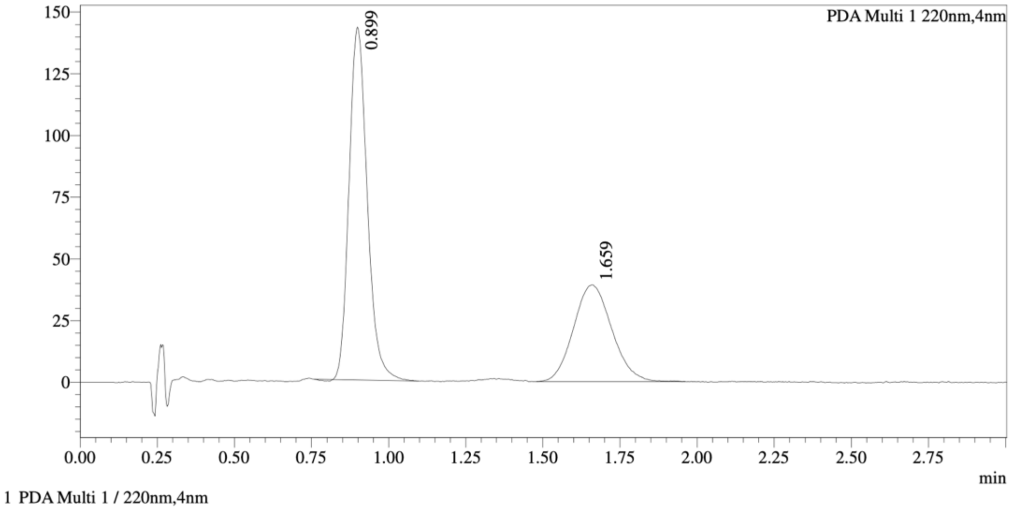


Integration Result

Peak Table

PDA Ch1 220nm

| Peak# | Ret. Time | Height | Height% | Resolution(USP) | Area | Area% |
|---|---|---|---|---|---|---|
| 1 | 0.899 | 141799 | 78.349 | -- | 586232 | 63.392 |
| 2 | 1.659 | 39185 | 21.651 | 4.503 | 338546 | 36.608 |

Method details

Column: Chiralcel OD-3, 50 mm length × 4.6 mm internal diameter, 3 μm particle size.
Mobile phase: A: supercritical $CO_2$; B: EtOH (0.05% diethylamine).
Isocratic elution: 40% B.

WX-02-30

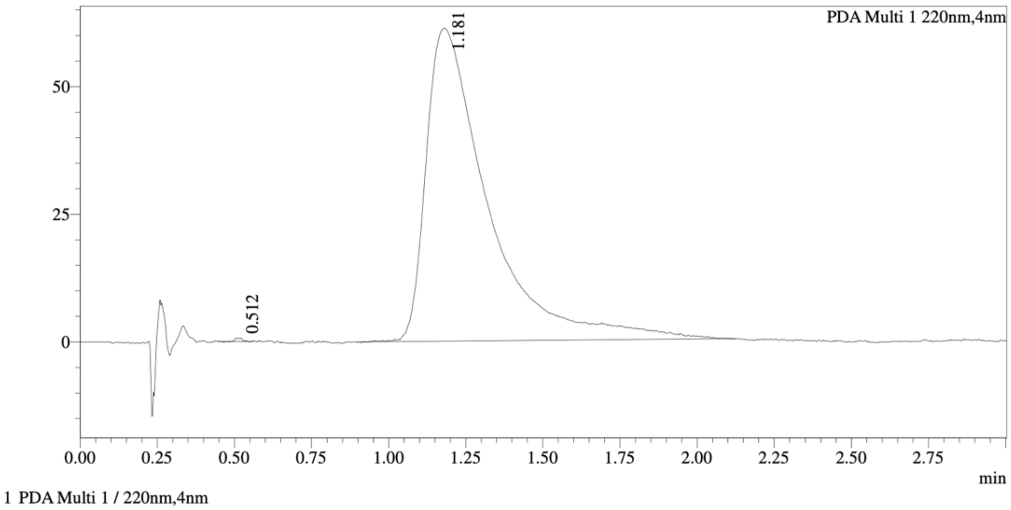


=====================================================================
Integration Result
=====================================================================

Peak Table

PDA Ch1 220nm

| Peak# | Ret. Time | Height | Height% | Resolution(USP) | Area | Area% |
|---|---|---|---|---|---|---|
| 1 | 0.512 | 622 | 1.005 | -- | 1016 | 0.118 |
| 2 | 1.181 | 61285 | 98.995 | 3.626 | 858014 | 99.882 |

WX-02-50

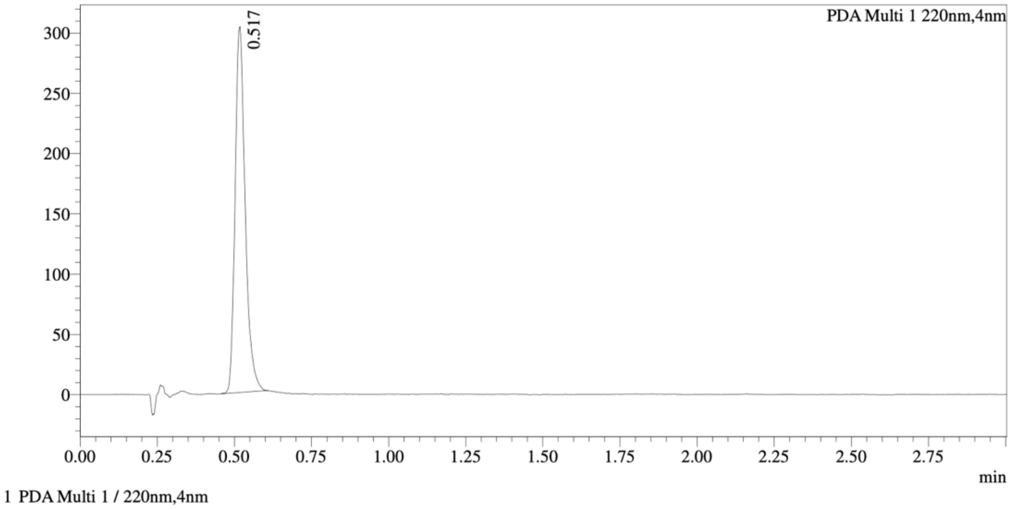


=====================================================================
Integration Result
=====================================================================

Peak Table

PDA Ch1 220nm

| Peak# | Ret. Time | Height | Height% | Resolution(USP) | Area | Area% |
|---|---|---|---|---|---|---|
| 1 | 0.517 | 300146 | 100.000 | -- | 691761 | 100.000 |

Mixture of WX-02-30 & WX-02-50

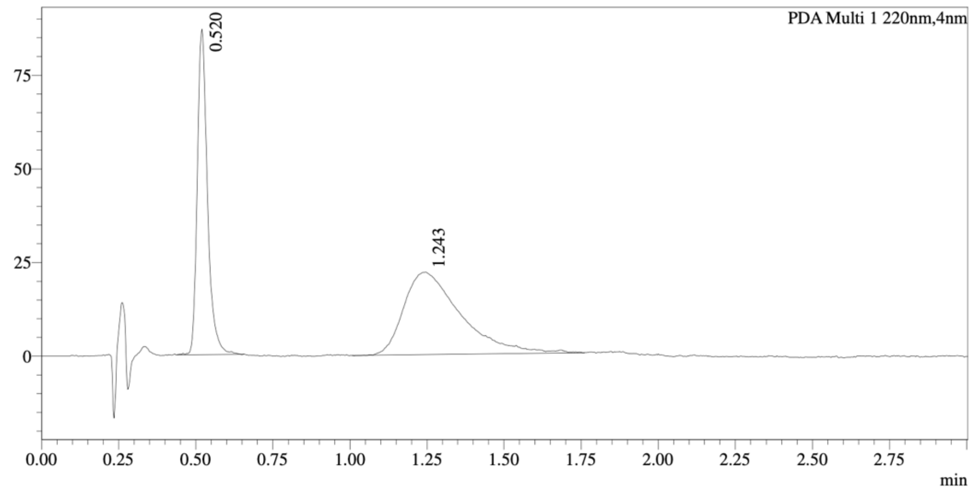

1 PDA Multi 1 / 220nm,4nm

====================================================================================
Integration Result
====================================================================================

Peak Table

PDA Ch1 220nm

| Peak# | Ret. Time | Height | Height% | Resolution(USP) | Area | Area% |
|---|---|---|---|---|---|---|
| 1 | 0.520 | 84890 | 79.414 | -- | 199609 | 41.351 |
| 2 | 1.243 | 22005 | 20.586 | 3.782 | 283105 | 58.649 |

Method details

Column: Chiralcel OD-3, 50 mm length × 4.6 mm internal diameter, 3 μm particle size.
Mobile phase: A: supercritical $CO_2$; B: EtOH (0.05% diethylamine).
Isocratic elution: 40% B.

WX-02-21

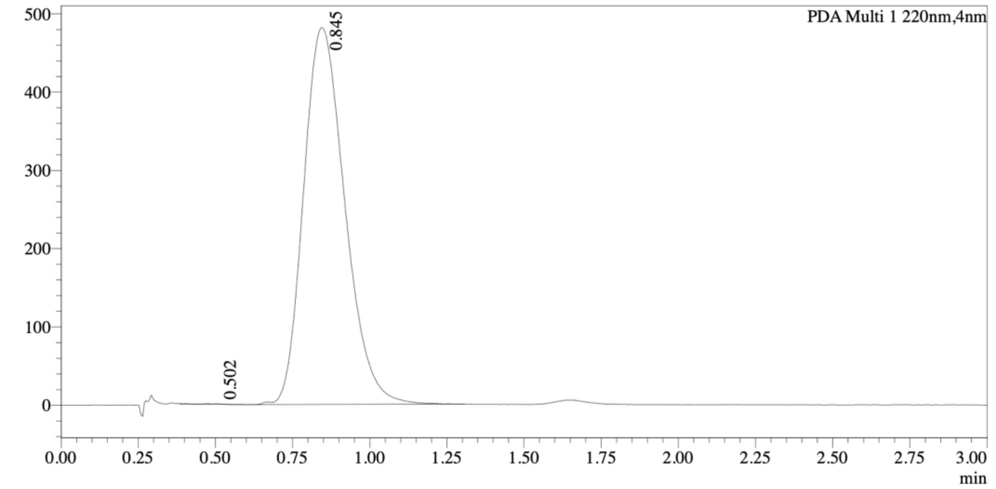


1 PDA Multi 1 / 220nm,4nm

Integration Result

Peak Table

PDA Ch1 220nm

| Peak# | Ret. Time | Height | Height% | Resolution(USP) | Area | Area% |
|---|---|---|---|---|---|---|
| 1 | 0.502 | 405 | 0.084 | -- | -1612 | -0.036 |
| 2 | 0.845 | 481081 | 99.916 | 2.406 | 4487118 | 100.036 |

WX-02-41

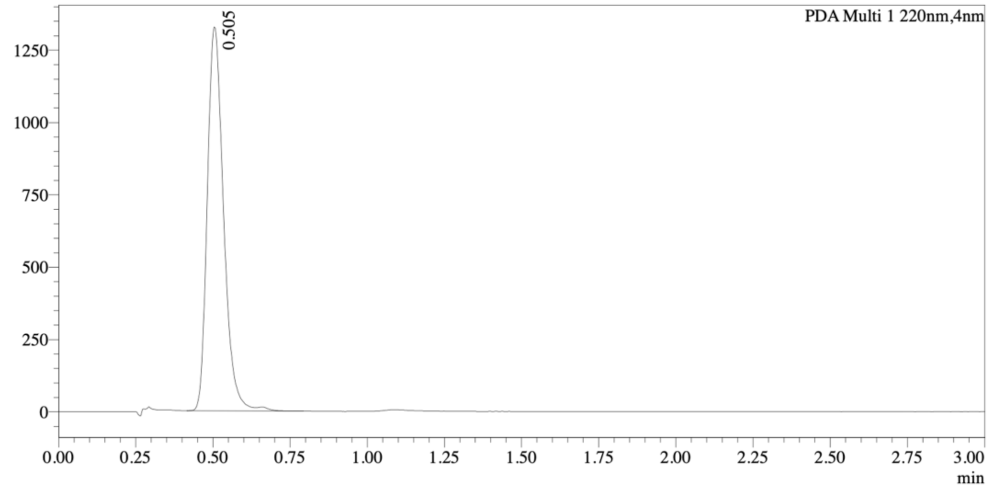


1 PDA Multi 1 / 220nm,4nm

Integration Result

Peak Table

PDA Ch1 220nm

| Peak# | Ret. Time | Height | Height% | Resolution(USP) | Area | Area% |
|---|---|---|---|---|---|---|
| 1 | 0.505 | 1306232 | 100.000 | -- | 4868953 | 100.000 |

Mixture of WX-02-21 & WX-02-41

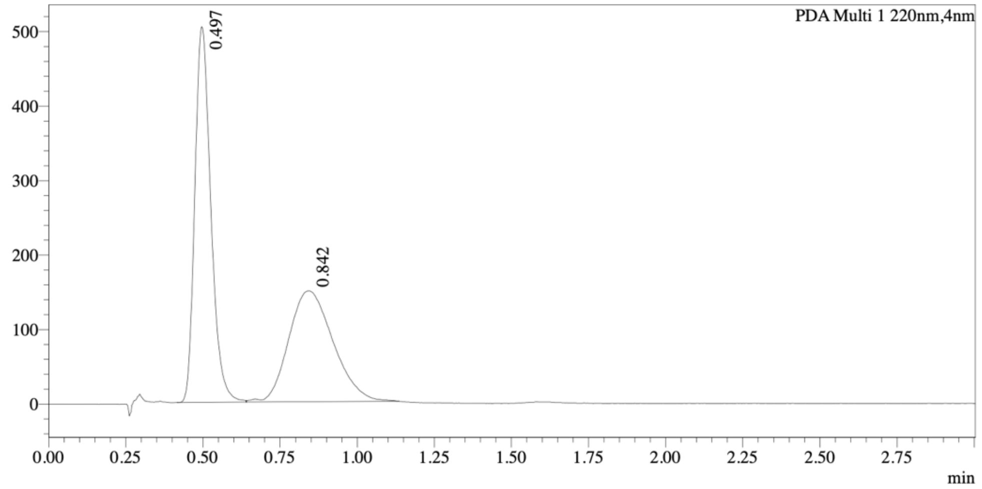


1 PDA Multi 1 / 220nm,4nm

Integration Result

Peak Table

PDA Ch1 220nm

| Peak# | Ret. Time | Height | Height% | Resolution(USP) | Area | Area% |
|---|---|---|---|---|---|---|
| 1 | 0.497 | 496954 | 76.963 | -- | 1823540 | 55.021 |
| 2 | 0.842 | 148750 | 23.037 | 1.925 | 1490717 | 44.979 |

Method details

Column: Chiralpak AD-3, 50 mm length × 4.6 mm internal diameter, 3 μm particle size.
Mobile phase: A: supercritical $CO_2$; B: 2:1 MeOH/MeCN (0.05% diethylamine).
Isocratic elution: 40% B.

WX-02-31

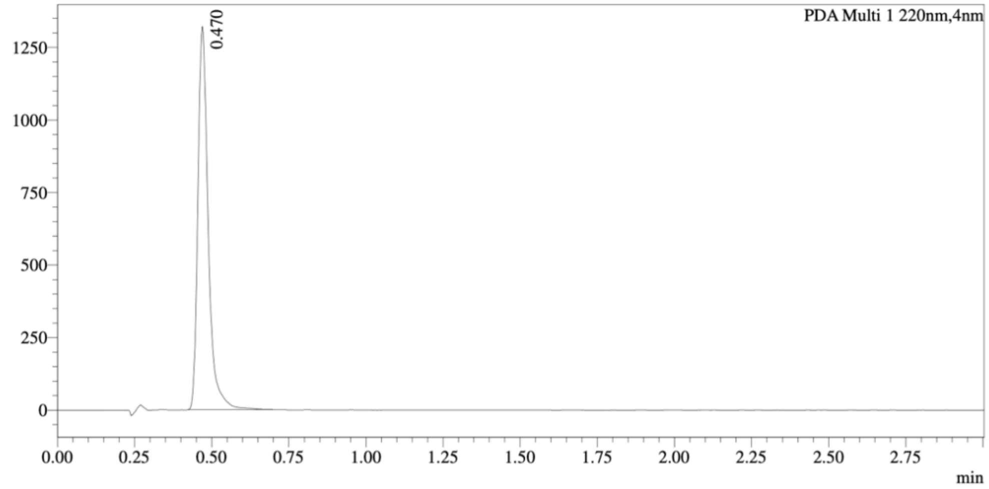


1 PDA Multi 1 / 220nm,4nm

Integration Result

Peak Table

PDA Ch1 220nm

| Peak# | Ret. Time | Height | Height% | Resolution(USP) | Area | Area% |
|---|---|---|---|---|---|---|
| 1 | 0.470 | 1305089 | 100.000 | -- | 3080588 | 100.000 |

WX-02-51

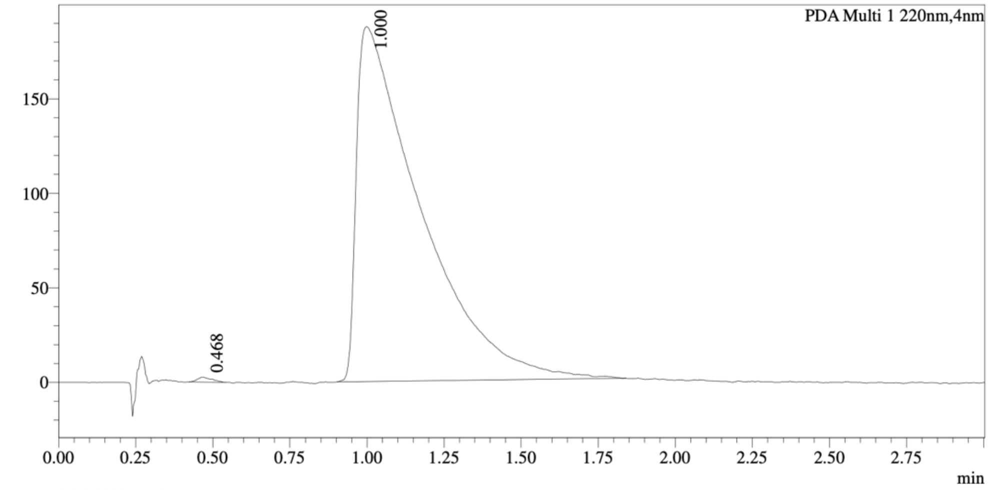


1 PDA Multi 1 / 220nm,4nm

Integration Result

Peak Table

PDA Ch1 220nm

| Peak# | Ret. Time | Height | Height% | Resolution(USP) | Area | Area% |
|---|---|---|---|---|---|---|
| 1 | 0.468 | 2575 | 1.354 | -- | 8501 | 0.312 |
| 2 | 1.000 | 187552 | 98.646 | 2.272 | 2718424 | 99.688 |

Mixture of WX-02-31 & WX-02-51

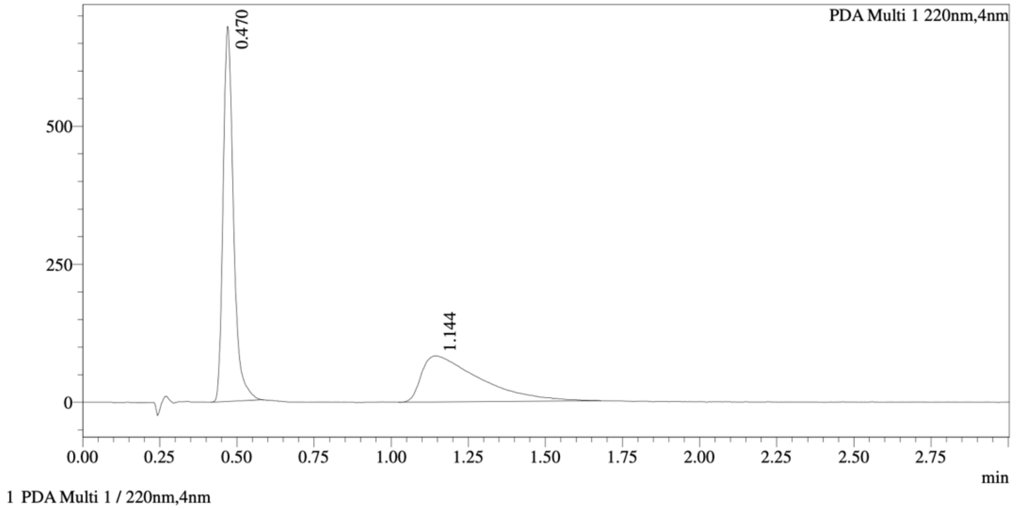


Integration Result

Peak Table

PDA Ch1 220nm

| Peak# | Ret. Time | Height | Height% | Resolution(USP) | Area | Area% |
|---|---|---|---|---|---|---|
| 1 | 0.470 | 671269 | 88.974 | -- | 1587162 | 59.187 |
| 2 | 1.144 | 83190 | 11.026 | 3.248 | 1094423 | 40.813 |

Method details

Column: Chiralcel OD-3, 50 mm length × 4.6 mm internal diameter, 3 μm particle size.
Mobile phase: A: supercritical $CO_2$; B: MeOH (0.05% diethylamine).
Isocratic elution: 40% B.

WX-02-22

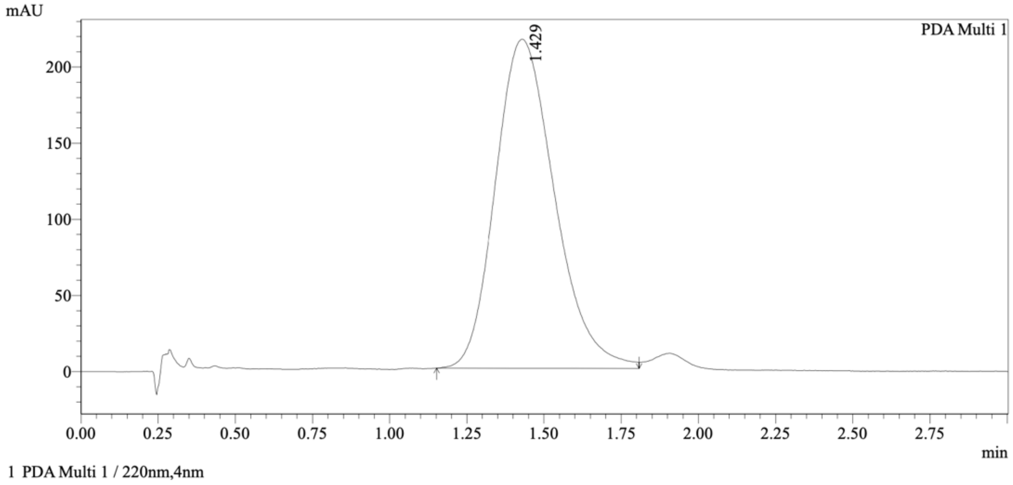


Integration Results

PeakTable

PDA Ch1 220nm

| Peak# | Ret. Time | USP Width | Resolution | Height | Area | Area % |
|---|---|---|---|---|---|---|
| 1 | 1.429 | 0.349 | 0.000 | 215984 | 2886386 | 100.000 |
| Total | | | | 215984 | 2886386 | 100.000 |

WX-02-42

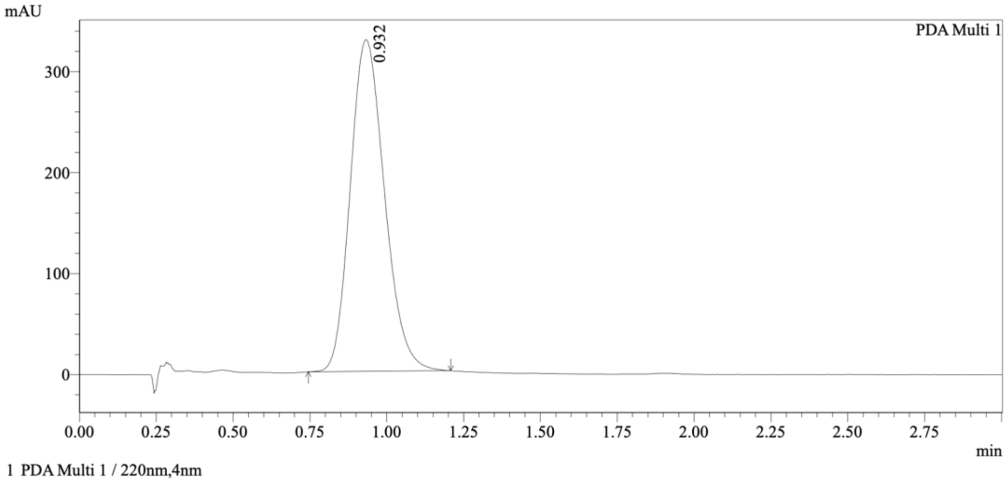


Integration Results

PeakTable

PDA Ch1 220nm

| Peak# | Ret. Time | USP Width | Resolution | Height | Area | Area % |
|---|---|---|---|---|---|---|
| 1 | 0.932 | 0.205 | 0.000 | 328045 | 2550933 | 100.000 |
| Total | | | | 328045 | 2550933 | 100.000 |

Mixture of WX-02-22 & WX-02-42

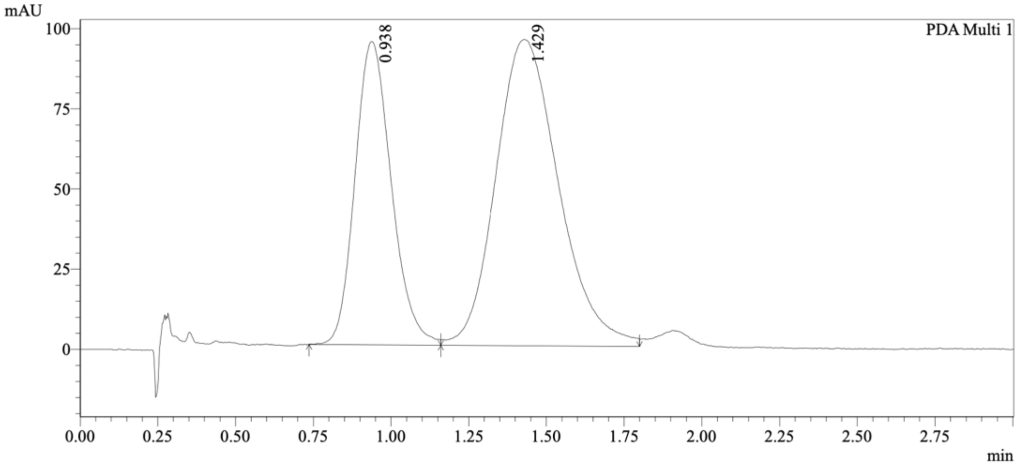


1 PDA Multi 1 / 220nm,4nm

Integration Results

PeakTable

PDA Ch1 220nm

| Peak# | Ret. Time | USP Width | Resolution | Height | Area | Area % |
|---|---|---|---|---|---|---|
| 1 | 0.938 | 0.215 | 0.000 | 94303 | 770307 | 36.395 |
| 2 | 1.429 | 0.366 | 1.694 | 95443 | 1346240 | 63.605 |
| Total | | | | 189746 | 2116548 | 100.000 |

Method details

Column: Chiralpak AD-3, 50 mm length × 4.6 mm internal diameter, 3 μm particle size.
Mobile phase: A: supercritical $CO_2$; B: MeOH (0.05% diethylamine).
Isocratic elution: 40% B.

WX-02-32

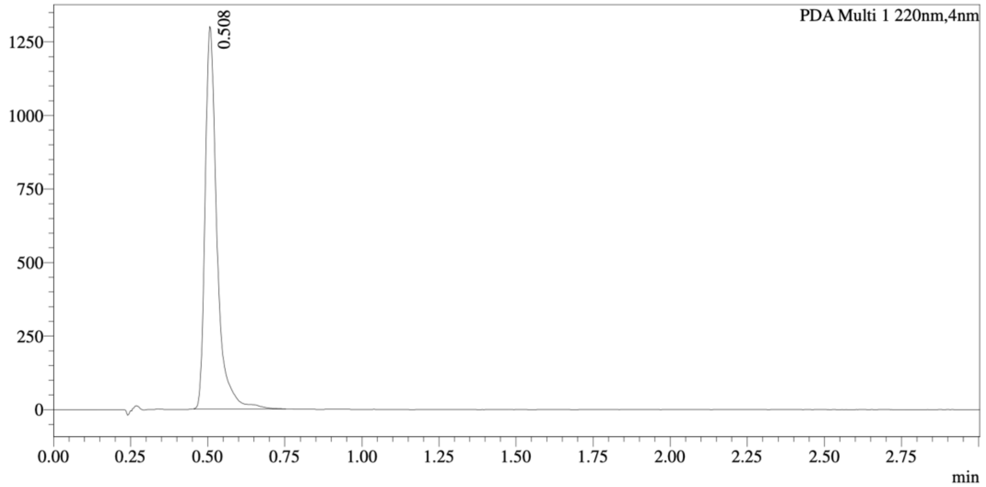


1 PDA Multi 1 / 220nm,4nm

Integration Result

Peak Table

PDA Ch1 220nm

| Peak# | Ret. Time | Height | Height% | Resolution(USP) | Area | Area% |
|---|---|---|---|---|---|---|
| 1 | 0.508 | 1279461 | 100.000 | -- | 3468410 | 100.000 |

WX-02-52

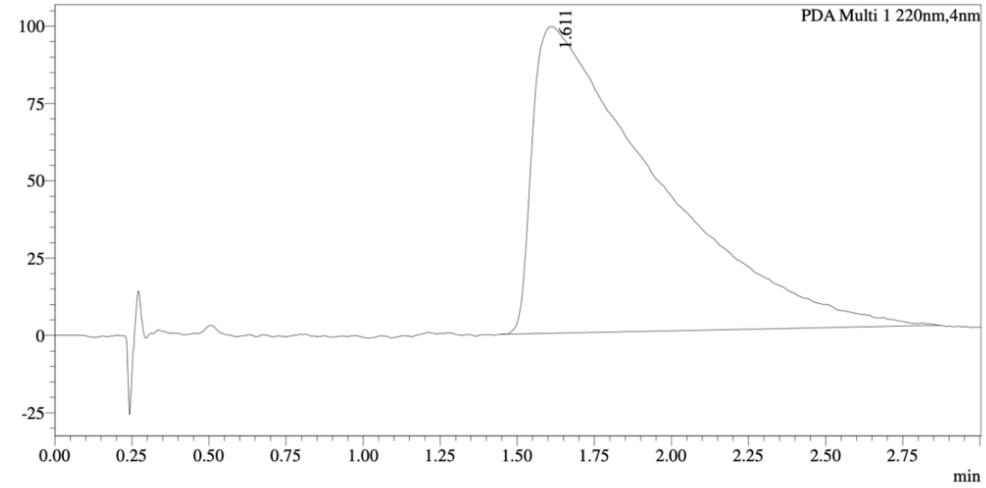


1 PDA Multi 1 / 220nm,4nm

Integration Result

Peak Table

PDA Ch1 220nm

| Peak# | Ret. Time | Height | Height% | Resolution(USP) | Area | Area% |
|---|---|---|---|---|---|---|
| 1 | 1.611 | 99103 | 100.000 | -- | 2794539 | 100.000 |

Mixture of WX-02-32 & WX-02-52

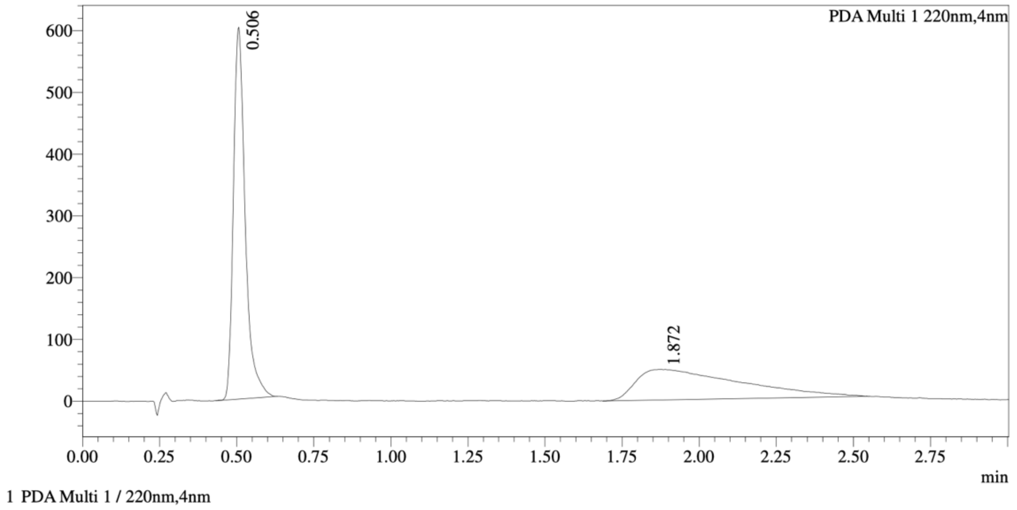


Integration Result

Peak Table

PDA Ch1 220nm

| Peak# | Ret. Time | Height | Height% | Resolution(USP) | Area | Area% |
|---|---|---|---|---|---|---|
| 1 | 0.506 | 584338 | 92.223 | -- | 1603318 | 58.190 |
| 2 | 1.872 | 49273 | 7.777 | 4.102 | 1152016 | 41.810 |

Method details

Column: Chiralcel OD-3, 50 mm length × 4.6 mm internal diameter, 3 μm particle size.
Mobile phase: A: supercritical $CO_2$; B: MeOH (0.05% diethylamine).
Isocratic elution: 40% B.

WX-02-654

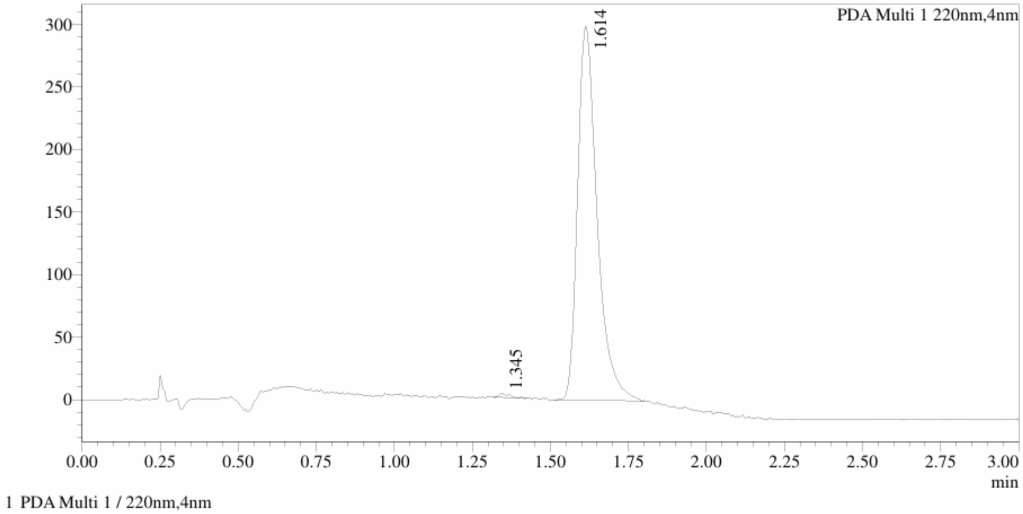


Integration Result

Peak Table

PDA Ch1 220nm

| Peak# | Ret. Time | Height | Height% | Resolution(USP) | Area | Area% |
|---|---|---|---|---|---|---|
| 1 | 1.345 | 2771 | 0.921 | -- | 6885 | 0.526 |
| 2 | 1.614 | 297977 | 99.079 | 3.130 | 1302250 | 99.474 |

WX-02-655

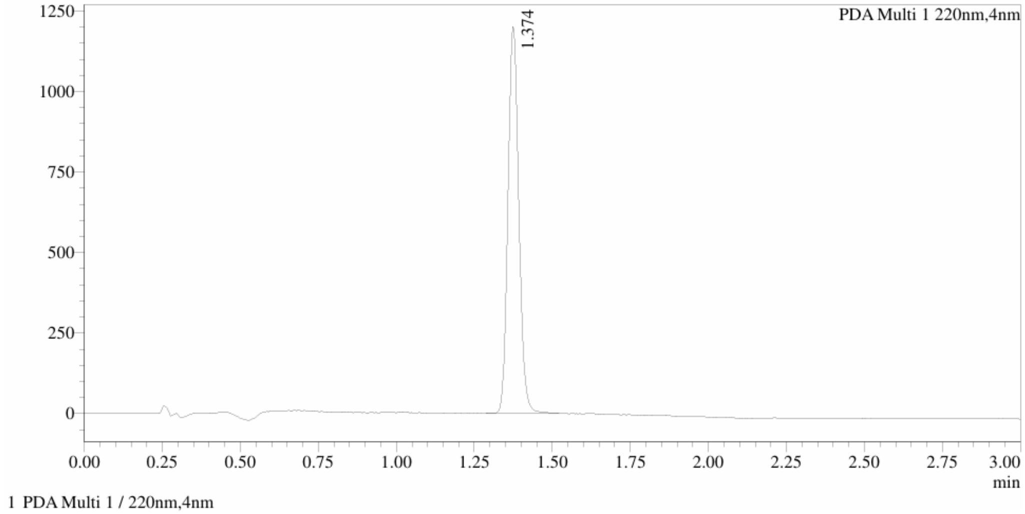


Integration Result

Peak Table

PDA Ch1 220nm

| Peak# | Ret. Time | Height | Height% | Resolution(USP) | Area | Area% |
|---|---|---|---|---|---|---|
| 1 | 1.374 | 1188799 | 100.000 | -- | 2907020 | 100.000 |

Mixture of WX-02-654 & WX-02-655

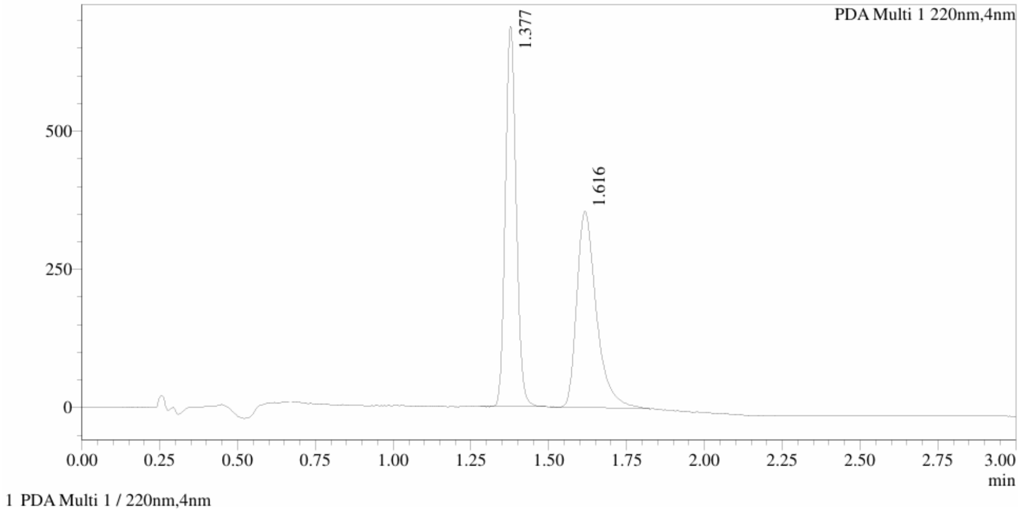


Integration Result

Peak Table

PDA Ch1 220nm

| Peak# | Ret. Time | Height | Height% | Resolution(USP) | Area | Area% |
|---|---|---|---|---|---|---|
| 1 | 1.377 | 664316 | 65.399 | -- | 1660706 | 51.619 |
| 2 | 1.616 | 351472 | 34.601 | 2.670 | 1556532 | 48.381 |

Method details

Column: Chiralpak AD-3, 50 mm length × 4.6 mm internal diameter, 3 μm particle size.
Mobile phase: A: supercritical $CO_2$; B: EtOH (0.05% diethylamine).
Gradient elution: 5 - 40% B.
